# Implications of *TP53* Allelic State for Genome Stability, Clinical Presentation and Outcomes in Myelodysplastic Syndromes

**DOI:** 10.1101/2019.12.19.868844

**Authors:** Elsa Bernard, Yasuhito Nannya, Robert P. Hasserjian, Sean M. Devlin, Heinz Tuechler, Juan S. Medina-Martinez, Tetsuichi Yoshizato, Yusuke Shiozawa, Ryunosuke Saiki, Luca Malcovati, Max F. Levine, Juan E. Arango, Yangyu Zhou, Francesc Sole, Catherine A. Cargo, Detlef Haase, Maria Creignou, Ulrich Germing, Yanming Zhang, Gunes Gundem, Araxe Sarian, Arjan A. van de Loosdrecht, Martin Jädersten, Magnus Tobiasson, Olivier Kosmider, Matilde Y. Follo, Felicitas Thol, Ronald F. Pinheiro, Valeria Santini, Ioannis Kotsianidis, Jacqueline Boultwood, Fabio P.S. Santos, Julie Schanz, Senji Kasahara, Takayuki Ishikawa, Hisashi Tsurumi, Akifumi Takaori-Kondo, Toru Kiguchi, Chantana Polprasert, John M. Bennett, Virginia M. Klimek, Michael R. Savona, Monika Belickova, Christina Ganster, Laura Palomo, Guillermo Sanz, Lionel Ades, Matteo Giovanni Della Porta, Alexandra G Smith, Yesenia Werner, Minal Patel, Agnès Viale, Katelynd Vanness, Donna S. Neuberg, Kristen E. Stevenson, Kamal Menghrajani, Kelly L. Bolton, Pierre Fenaux, Andrea Pellagatti, Uwe Platzbecker, Michael Heuser, Peter Valent, Shigeru Chiba, Yasushi Miyazaki, Carlo Finelli, Maria Teresa Voso, Lee-Yung Shih, Michaela Fontenay, Joop H. Jansen, José Cervera, Yoshiko Atsuta, Norbert Gattermann, Benjamin L. Ebert, Rafael Bejar, Peter L. Greenberg, Mario Cazzola, Eva Hellström-Lindberg, Seishi Ogawa, Elli Papaemmanuil

**Affiliations:** Computational Oncology Service, Department of Epidemiology & Biostatistics, Memorial Sloan Kettering Cancer Center, New York, NY, USA; Center for Hematologic Malignancies, Memorial Sloan Kettering Cancer Center, New York, NY, USA; Department of Pathology and Tumor Biology, Kyoto University, Kyoto, Japan; Department of Pathology, Massachusetts General Hospital, Boston, MA, USA; Department of Epidemiology & Biostatistics, Memorial Sloan Kettering Cancer Center, New York, NY, USA; Department of Molecular Medicine, University of Pavia, Pavia. Italy; Department of Hematology, IRCCS Fondazione Policlinico S. Matteo, Pavia. Italy; MDS Group, Institut de Recerca Contra la Leucèmia Josep Carreras, Barcelona, Spain; Haematological Malignancy Diagnostic Service, St James’s University Hospital, Leeds, United Kingdom; Clinics of Hematology and Medical Oncology, University Medical Center. Göttingen, Germany; Department of Medicine Huddinge, Center for Hematology and Regenerative Medicine, Karolinska Institutet, Karolinska University Hospital, Stockholm, Sweden; Department of Hematology, Oncology and Clinical Immunology, Heinrich Heine University, Düsseldorf, Germany; Department of Pathology, Memorial Sloan Kettering Cancer Center, New York, NY, USA; Department of Hematology, VU University Medical Center Amsterdam, The Netherlands; Department of Hematology, Assistance Publique-Hôpitaux de Paris, Hôpital Cochin and Université de Paris, Université Paris Descartes, Paris, France; Department of Biomedical and Neuromotor Sciences, University of Bologna, Bologna, Italy; Department of Hematology, Hemostasis, Oncology and Stem Cell Transplantation, Hannover Medical School, Hannover, Germany; Drug Research and Development Center, Federal University of Ceara, Ceara, Brazil; MDS Unit, Hematology, AOU Careggi, University of Florence, Italy; Department of Hematology, Democritus University of Thrace Medical School, Alexandroupolis, Greece; Radcliffe Department of Medicine, University of Oxford and Oxford BRC Haematology Theme, Oxford, United Kingdom; Oncology-Hematology Center, Hospital Israelita Albert Einstein, São Paulo, Brazil; Department of Hematology, Gifu Municipal Hospital, Gifu, Japan; Department of Hematology, Kobe City Medical Center General Hospital, Kobe, Japan; Department of Hematology, Gifu University Graduate School of Medicine, Gifu, Japan; Department of Hematology and Oncology, Graduate School of Medicine, Kyoto University, Japan; Department of Hematology, Chugoku Central Hospital, Fukuyama, Japan; Department of Medicine, Chulalongkorn University, King Chulalongkorn Memorial Hospital, Bangkok, Thailand; Lab. Medicine and Pathology, Hematology/Oncology, University of Rochester Medical Center, Rochester, NY, USA; Department of Medicine, Memorial Sloan Kettering Cancer Center, New York, NY, USA; Department of Medicine, Vanderbilt-Ingram Cancer Center, Vanderbilt University School of Medicine, Nashville, TN, USA; Department of Genomics, Institute of Hematology and Blood Transfusion, Prague, Czech Republic; Department of Hematology, Hospital Universitario y Politécnico La Fe, Valencia, Spain; CIBERONC, Instituto de Salud Carlos III, Madrid, Spain; Department of Hematology, hôpital St Louis, and Paris University, Paris, France; Cancer Center, Humanitas Research Hospital & Humanitas University, Milan Italy; Department of Health Sciences, University of York, United Kingdom; Integrated Genomics Operation, Memorial Sloan Kettering Cancer Center, New York, NY, USA; Department of Data Sciences, Dana-Farber Cancer Institute, Boston, MA, USA; Medical Clinic and Policlinic 1, Hematology and Cellular Therapy, University of Leipzig, Leipzig, Germany; Department of Internal Medicine I, Division of Hematology and Hemostaseology and Ludwig Boltzmann Institute for Hematology and Oncology, Medical University of Vienna, Austria; Department of Hematology, Faculty of Medicine, University of Tsukuba, Tsukuba, Japan; Department of Hematology, Atomic Bomb Disease Institute, Nagasaki University, Japan; Institute of Hematology, S.Orsola-Malpighi University Hospital, Bologna, Italy; MDS Cooperative Group GROM-L, Department of Biomedicine and Prevention, Tor Vergata University, Rome, Italy; Chang Gung Memorial Hospital at Linkou, Chang Gung University, Taiwan; Laboratory Hematology, Deptartment LABGK, Radboud University Medical Centre, Nijmegen, The Netherlands; Department of Hematology and Genetics Unit, University Hospital La Fe, Valencia, Spain; Japanese Data Center for Hematopoietic Cell Transplantation, Nagoya, Japan; Department of Medical Oncology and Howard Hughes Medical Institute, Dana-Farber Cancer Center, Boston, MA, USA; UC San Diego Moores Cancer Center, La Jolla, California, USA; Stanford University Cancer Institute, Stanford, CA, USA

**Author notes:** Shared senior authorship (Seishi Ogawa, Elli Papaemmanuil).

## Abstract

*TP53* mutations are associated with poor clinical outcomes and treatment resistance in myelodysplastic syndromes. However, the biological and clinical relevance of the underlying mono- or bi-allelic state of the mutations is unclear. We analyzed 3,324 MDS patients for *TP53* mutations and allelic imbalances of the *TP53* locus and found that 1 in 3 *TP53*-mutated patients had mono-allelic targeting of the gene whereas 2 in 3 had multiple hits consistent with bi-allelic targeting. The established associations for *TP53* with complex karyotype, high-risk presentation, poor survival and rapid leukemic transformation were specific to patients with multi-hit state only. *TP53* multi-hit state predicted risk of death and leukemic transformation independently of the Revised International Prognostic Scoring System, while mono-allelic patients did not differ from *TP53* wild-type patients. The separation by allelic state was retained in therapy-related MDS. Findings were validated in a cohort of 1,120 patients. Ascertainment of *TP53* allelic state is critical for diagnosis, risk estimation and prognostication precision in MDS, and future correlative studies of treatment response should consider *TP53* allelic state.

## INTRODUCTION

*TP53* is the most frequently mutated gene in cancer^1, 2^. In patients with myelodysplastic syndromes (MDS), *TP53* mutations have consistently been associated with high-risk disease features such as complex karyotype^3^, elevated blasts and severe thrombocytopenia^4^. *TP53*-mutated patients have dismal outcomes^5^, rapid transformation^6^ to acute myeloid leukemia (AML) and resistance to conventional therapies^7^. Recent studies suggest that *TP53* mutations are predictive of relapse following hematopoietic stem cell transplantation (HSCT)^8, 9^ and of disease progression during lenalidomide treatment in the context of del(5q)^10^. Upon AML progression, *TP53* mutations demarcate an extremely adverse prognostic group associated with a chemo-refractory disease and less than 2% 5-year survival^11, 12^. Therapy-related MDS with *TP53* mutations is similarly associated with dismal outcomes^8, 13^. These observations illustrate a central role of *TP53* in the pathogenesis of myeloid neoplasms and highlight its relevance as a prognostic and predictive biomarker. However, *TP53* mutations are not yet considered in clinical risk scores such as the Revised International Prognostic Scoring System (IPSS-R)^14^ for MDS.

The majority of *TP53* mutations are missense variants clustering within the DNA binding domain (DBD). Consistent with its role as a tumor suppressor, bi-allelic targeting is mediated by loss of heterozygosity (LOH) involving 17p13 locus, commonly caused by deletion^15^. However, patients present with both mono- and bi-allelic mutations. Functional studies link specific *TP53* mutations with gain of function (GOF) and dominant negative effect (DNE)^16,17,18^, which may explain the diverse presentation of *TP53* mutations. Beyond the profound negative effect of *TP53* mutations, the clinical impact of bi-allelic vs. mono-allelic *TP53* mutations on outcomes and response to therapy has not been fully investigated.

We set out to study profiles of genome stability, clinical phenotypes and outcomes of MDS patients with *TP53* mutations in the context of the allelic state. In collaboration with the International Working Group for Prognosis in MDS (IWG-PM) (Supplementary Table. 1), we analyzed a cohort of 3,324 peri-diagnostic and treatment naive patients with MDS or closely related myeloid neoplasms (Extended Data Table 1 and Supplementary Fig. 1). Patient samples were representative of all MDS WHO subtypes and IPSS-R risk groups, and included 563 (17%) MDS/MPN and 167 (5%) AML/AML with myelodysplasia-related changes (AML-MRC) samples. An additional 1,120 samples derived from the Japanese MDS consortium (Extended Data Table 2) were used as a validation cohort. We described a detailed catalogue of mutagenic processes targeting the *TP53* locus, encompassing acquired mutations, copy-neutral loss of heterozygosity (cnLOH), focal and arm level deletions. We defined distinct *TP53* allelic states and showed that each state is associated with unique profiles of genome stability and clinical presentation. Our findings are of immediate clinical relevance with implications for diagnostic assay development, reporting guidelines and risk stratification of MDS patients.

## RESULTS

### Characterization of genome wide allelic imbalances in MDS

Genetic profiling included conventional G-banding analyses (CBA) and a custom capture next generation sequencing (NGS) panel that covered *TP53* and genome wide copy-number probes. Allele specific copy-number profiles were generated from NGS data using CNACS^9^. CBA data were available for 2,931 (88%) patients. Comparison of NGS-derived ploidy alterations to CBA-derived ones showed highly concordant results between the two assays (Supplementary Fig. 2 and 3a-b), which allowed us to complement the dataset with NGS findings for 393 cases with missing CBA (Supplementary Fig. 3c). Our custom capture approach further enabled the detection of focal (∼3MB) gains or deletions and regions of cnLOH (Supplementary Fig. 4 and Extended Data Fig. 1). Eleven percent of patients (n=360) had at least one cnLOH region, frequently targeting chr17/17p, chr4q, chr7q, chr11q, chr1p and chr14q (Extended Data Fig. 1b). Collectively, 1,571 (47%) patients had one or more chromosomal aberration, of which 329 (10%) had a complex karyotype^19^ and 177 (5%) had a monosomal karyotype^20^ (Supplementary Table 2).

### *TP53* mutation landscape in MDS

We identified 486 mutations in *TP53* across 378 individuals. Mutations in *TP53* were annotated as putative oncogenic as previously described^12, 21, 22^ by consideration of 1. Prior evidence in cancer databases^23,24,25^; 2. Recurrence in myeloid disease^5, 22, 26^; 3. Variant allele frequency (VAF) consistent with somatic representation; 4. Technical controls; and 5. Germline databases^27, 28^. The spectrum of identified *TP53* mutations followed patterns from systematic sequencing studies (Supplementary Fig. 5). As expected, 71% of the mutations were missense variants clustered within the DBD. The 4 most common hotspots (R273, R248, Y220 and R175) accounted for 21% of all mutations.

Among the 378 patients with *TP53* mutations, 274 (72.5%) had a single *TP53* mutation, 100 had two (26.5%) and 4 (1%) had three (Supplementary Fig. 6). We mapped deletions of the *TP53* locus in 97 cases, of which 18 were focal events detected by NGS-based analysis only. We also identified 80 cases with cnLOH which were not detected by CBA (Supplementary Table 3). Approximately half (54%, n=149) of the patients with one *TP53* mutation had loss of the wild-type allele by deletion or cnLOH. In contrast, only 13% (n=14) of patients with ≥2 *TP53* mutations had a concomitant allelic imbalance at the *TP53* locus (OR=8, p<10^-13^ Fisher exact test) (Fig. 1a). According to the number of mutations and the presence of deletion or cnLOH, we defined 4 main *TP53*-mutant subgroups (Fig. 1b): 1. Mono-allelic mutation (n=125, 33% of *TP53*-mutated patients); 2. Multiple mutations without deletion or cnLOH affecting the *TP53* locus (n=90, 24%); 3. Mutation(s) and concomitant deletion (n=85, 22%); 4. Mutation(s) and concomitant cnLOH (n=78, 21%). Additionally, in 24 patients, the *TP53* locus was affected by deletion (n=12), cnLOH (n=2) or isochromosome 17q rearrangement (n=10) without evidence of *TP53* mutations (Fig. 1a).

**Figure 1.**
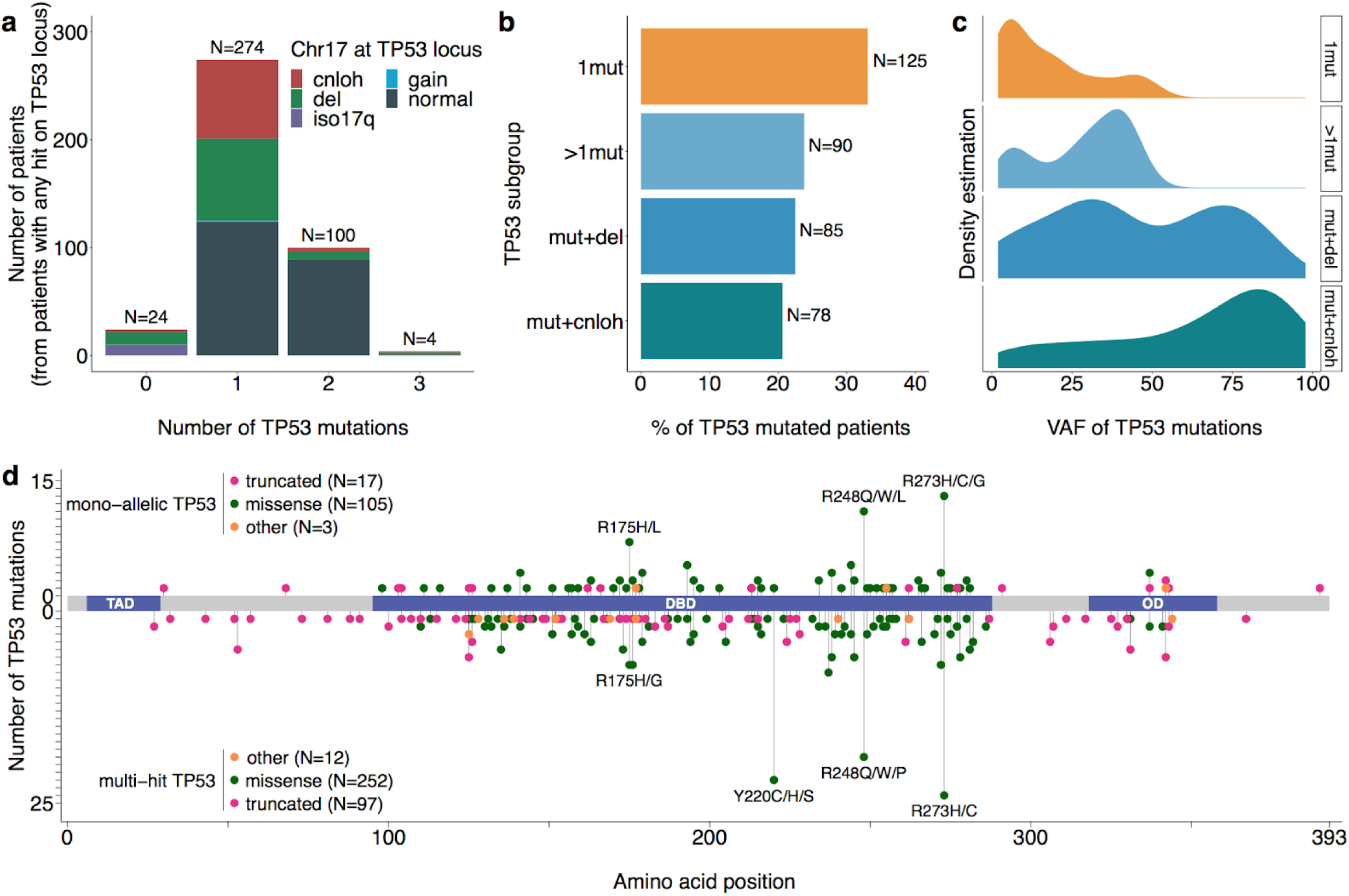
Integration of *TP53* mutations and allelic imbalances at the *TP53* locus identifies *TP53* states with evidence of mono-allelic or bi-allelic targeting. **a**, Number of patients (from patients with any hit at the *TP53* locus) with 0, 1, 2 or 3 *TP53* mutations. Colors represent the status of chromosome 17 at the *TP53* locus, to include copy-neutral loss of heterozygosity (cnloh), deletion (del), isochromosome 17q rearrangement (iso17q), gain or no detected aberration (normal). **b**, Frequency of *TP53* subgroups within *TP53*-mutated patients. *TP53* subgroups are defined as cases with i) single gene mutation (1mut) ii) several mutations with normal status of chromosome 17 at the *TP53* locus (>1mut) iii) mutation(s) and chromosomal deletion at the *TP53* locus (mut+del) and iv) mutation(s) and copy-neutral loss of heterozygosity at the *TP53* locus (mut+cnloh). **c**, Density estimation of variant allele frequency (VAF) of *TP53* mutations across *TP53* subgroups (1mut, >1mut, mut+del, mut+cnloh from top to bottom). **d**, Distribution of *TP53* mutations along the gene body. Mutations from patients with mono-allelic *TP53* per single gene mutation are depicted at the top, mutations from patients with multiple *TP53* hits at the bottom. Missense mutations are shown as green circles. Truncated mutations, including nonsense or nonstop mutations, frameshift deletions or insertions and splice site variants are shown as pink circles. Other types of mutations to include inframe deletions or insertions are shown as orange circles. TAD: transactivation domain; DBD: DNA binding domain; OD: oligomerization domain.

VAF measurements confirmed that the majority of multi-hit cases were indeed bi-allelic. In patients with ≥2 mutations, the VAFs of mutation pairs were strongly correlated (R^2^=0.77, Extended Data Fig. 2a), indicative of bi-allelic state. In 67% (n=60) of those cases, the mutations occurred with certainty in the same cells, with a cumulated VAF exceeding 50% (*the pigeonhole principle*^29^), thus confirming bi-allelic state. In addition, mutations within sequencing read length were systematically observed in trans, i.e., on different alleles (Extended Data Fig. 2b-c). In patients with an allelic imbalance at *TP53*, VAF estimates were enriched for values greater than 50%, consistent with loss of the wild-type allele (Fig. 1c). Taken together, VAF measurements supported bi-allelic targeting of *TP53* on cases with multiple mutations or mutation(s) and allelic imbalances (subgroups 2-4). However, VAF alone was not sufficient to determine allelic state. For example, we identified 19 cnLOH-positive patients with ≤50% *TP53* VAF (median 29%, range 3-49%), suggesting that 24% of cnLOH patients would be misassigned as mono-allelic on the basis of VAF. Therefore, accurate determination of *TP53* allelic state cannot solely rely on *TP53* mutation VAF and should consider LOH mapping, as can be achieved by NGS-based analysis of targeted gene sequencing panels with copy number probes, that are increasingly routine in clinical practice. In mono-allelic cases, VAF densities suggested that *TP53* mutations were enriched for subclonal presentation (median VAF: 13%, median sample purity: 86%) as compared to *TP53* mutations from patients with multiple mutations which were predominantly clonal (median VAF: 32%, median sample purity: 85%) (Fig. 1c).

**Figure 2.**
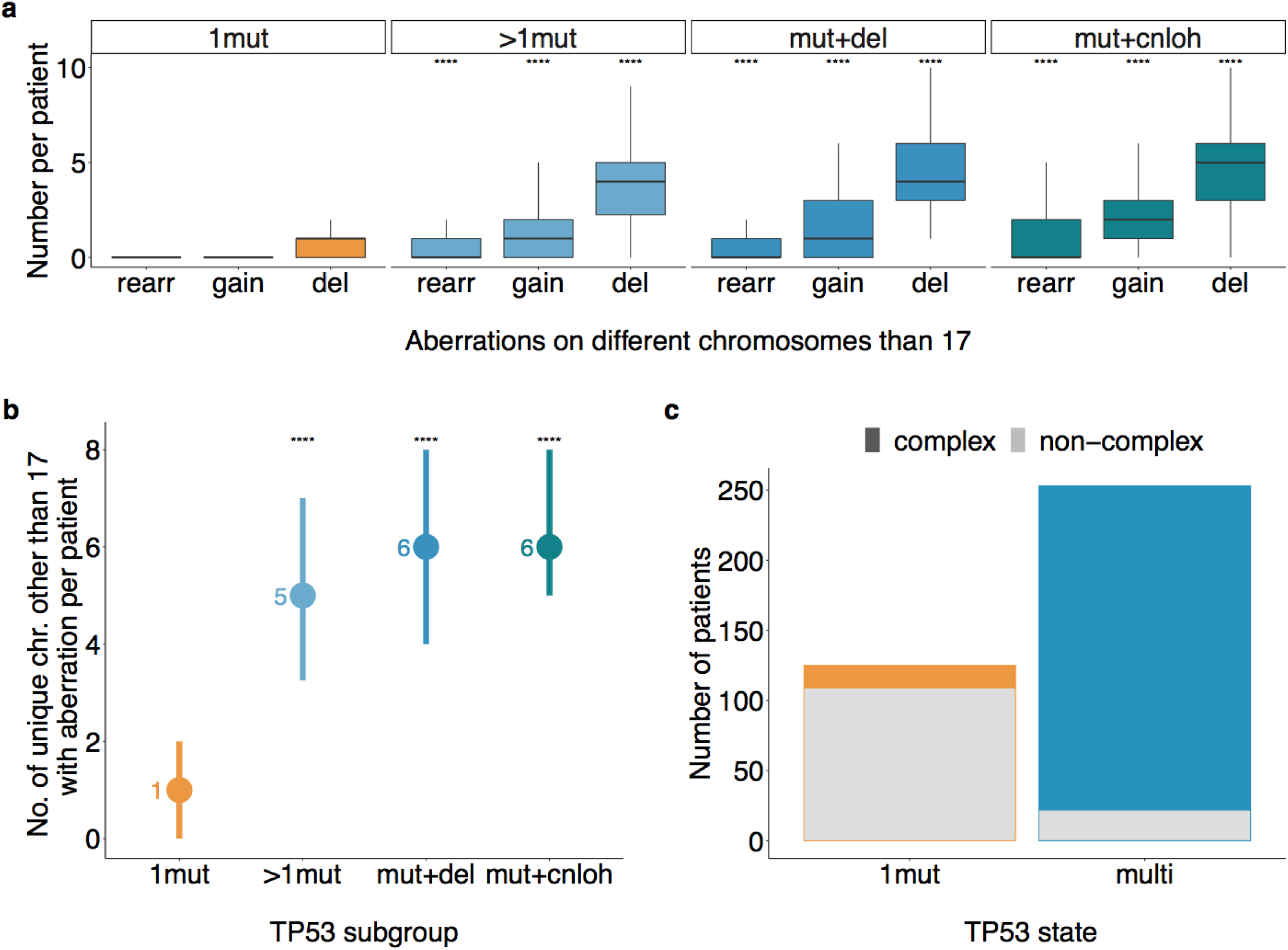
Genome instability within the multi-hit *TP53* subgroups but not the mono-allelic subgroup. **a**, Distribution of the number of chromosomal aberrations on other chromosomes than 17 per patient across *TP53* subgroups and types of aberrations, i.e., rearrangement (rearr), gain or deletion (del). ****p<0.0001, Wilcoxon rank-sum test, each compared to the same aberration within the 1mut group. **b**, Number of unique chromosomes other than 17 affected by a chromosomal aberration (rearrangement, deletion or gain) per *TP53* subgroup. Dots represent the median across patients and lines extend from 25% to 75% quantiles. ****p<0.0001, Wilcoxon rank-sum test, compared to the 1mut group. **c**, Interaction between *TP53* allelic state and complex karyotype. 13% (16/125) of mono-allelic *TP53* patients (1mut) had a complex karyotype. Conversely, 91% (231/253) of multi-hit *TP53* patients (multi) had a complex karyotype (OR=70, 95% CI: 33-150, p<10^-16^ Fisher exact test).

We organized the *TP53*-mutant subgroups into two states: A. mono-allelic *TP53* state representing subgroup 1, and B. multi-hit *TP53* state encompassing subgroups 2-4, with evidence of at least two *TP53* hits in each patient. While the multi-hit state most likely reflects the presence of clones with bi-allelic targeting, we maintained both “bi-allelic” and “multi-hit” terminology.

Overall, the *TP53* allelic states had shared repertoire of mutations (Fig. 1d and Supplementary Fig. 7). Of note, truncating mutations were enriched in the multi-hit state (28% vs. 14%, OR=2.3, p=0.002 Fisher exact test) while hotspot mutations accounted for 25% of mutations in the mono-allelic state and 20% in the multi-hit state (OR=1.38, p=0.2 Fisher exact test). The differing fractions of hotspot and truncating mutations between states might reflect discrete functional impacts of both mutation types, representing dominant negative vs. simple loss-of-function. In addition, *TP53* allelic state, and by extension whether a wild-type *TP53* allele is retained, points towards differential potential for clonal dominance, whereby the mono-allelic state was confined to smaller sub-clones and the multi-hit state was most frequently clonal.

### Implications of *TP53* allelic state to genome stability

The association between *TP53* mutations and chromosomal aneuploidies is well established^3, 9, 11, 12^. Overall, 67% (n=252) of *TP53*-mutated cases had ≥2 chromosomal deletions as compared to 5% (n=158) of wild-type cases (OR=35, p<10^-16^ Fisher exact test). Excluding chr17 (which is linked to state definition), there was a significantly higher number of chromosomal aberrations per patient, across rearrangements, gains and deletions, in all *TP53* subgroups of multiple hits compared to the mono-allelic state (Fig. 2a and Extended Data Fig. 3). This enrichment was most pronounced for deletions (median 4 in multi-hit vs. 1 in mono-allelic state, p<10^-16^ Wilcoxon rank-sum test, Fig. 2a). In particular, deletion of 5q was observed in 85% of multi-hit patients as opposed to 34% of mono-allelic patients (OR=10, p<10^-16^ Fisher exact test, Supplementary Fig. 8). Taken together, we found a median of 6 unique chromosomal aberration in the multi-hit state and 1 in the mono-allelic state (p<10^-16^ Wilcoxon rank-sum test, Fig. 2b). Our data suggest that residual wild-type *TP53* is critical to maintenance of genome stability, and that the association between *TP53* and complex karyotype is specific to the multi-hit state (91% vs. 13% complex karyotype patients within multi-hit or mono-allelic states, OR=70, p<10^-16^ Fisher exact test, Fig. 2c).

### *TP53* allelic state associates with distinct clinical phenotype and shapes patient outcomes

Without discriminating allelic states, previous studies uniformly reported adverse effects of *TP53* mutations on clinical phenotypes and outcome^3, 4^. These were recapitulated in our study (Supplementary Fig. 9 and 10). However, when analyzed separately, the two *TP53* allelic states associated with distinct clinical presentation and outcomes.

Mono-allelic *TP53* patients were less cytopenic (Fig. 3a-c) and had lower percentages of bone marrow blasts compared to multi-hit patients (median 4 vs. 9%, p<10^-10^ Wilcoxon rank sum test, Fig. 3d). There was a higher prevalence of lower risk MDS subtypes such as isolated del(5q) in mono-allelic patients, while the multi-hit state was enriched for higher risk WHO subtypes (Extended Data Fig. 4a). In IPSS-R, 46% of the mono-allelic *TP53* cases classified as good/very-good and 29% as poor/very-poor risk, whereas only 5.5% of multi-hit cases stratified as good/very-good and 89% as poor/very-poor risk (Extended Data Fig. 4b). These observations link *TP53* allelic state to disease presentation, WHO and IPSS-R risk classifications, whereby multi-hit state was enriched in higher risk disease.

**Figure 3.**
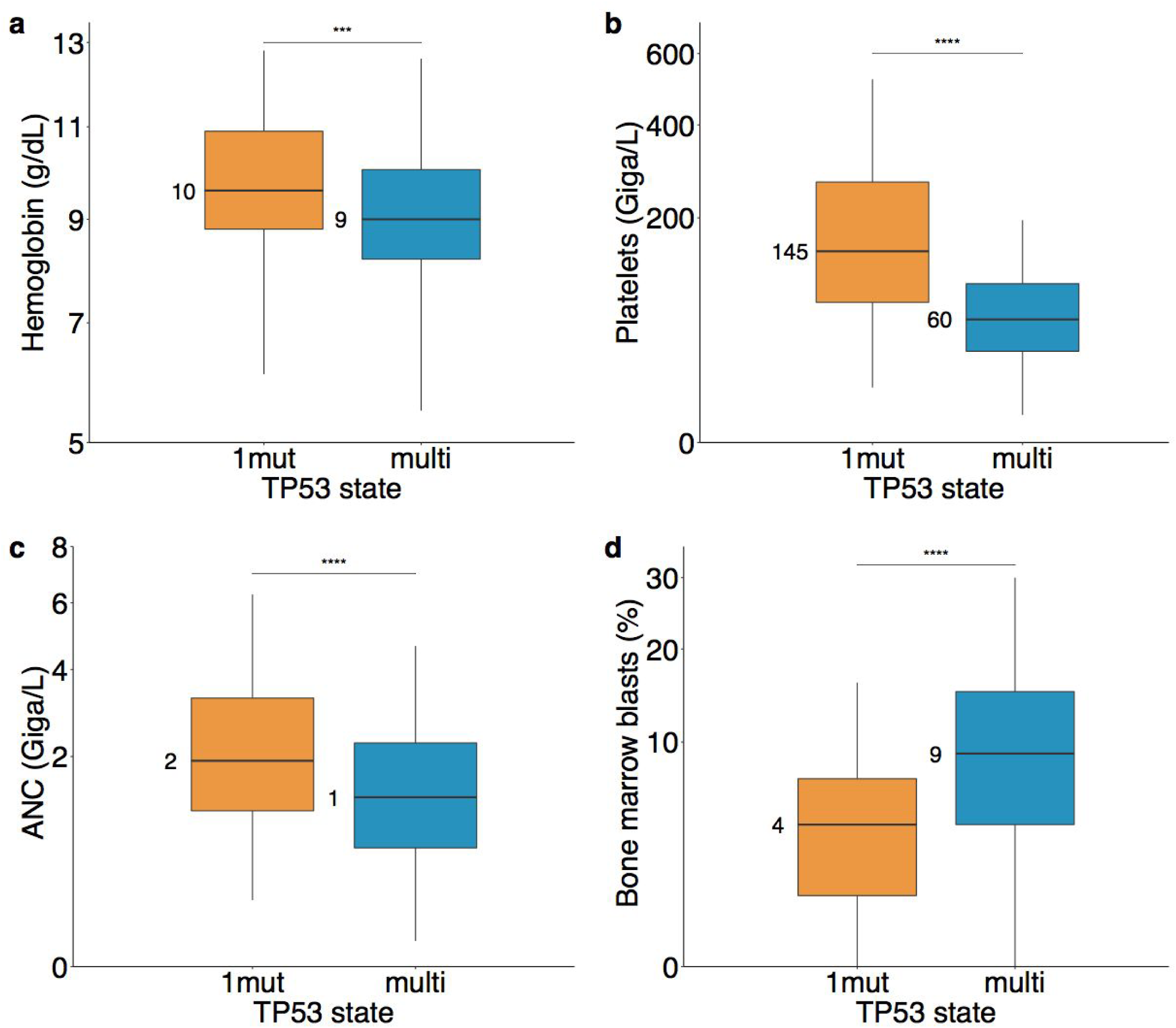
*TP53* allelic state associates with distinct clinical phenotypes. **a-c**. Boxplots indicative of the levels of cytopenias per *TP53* state of single gene mutation (1mut) or multiple hits (multi), respectively hemoglobin in panel a., platelets in panel b. and absolute neutrophil count (ANC) in panel c. Black lines represent the median across patients and filled boxes extend from 25% to 75% quantiles. The y-axis are square-root transformed. **d**. Percentage of bone marrow blasts per *TP53* state of a single gene mutation (1mut) or multiple hits (multi). ***p<0.001, ****p<0.0001, Wilcoxon rank-sum test.

The two allelic states had very different effects on overall survival (OS) and AML transformation. The median OS in mono-allelic *TP53* state was 2.5 years (95% CI: 2.2-4.9 years) and 8.7 months in the multi-hit state (95% CI: 7.7-10.3 months) (HR=3.7, 95% CI: 2.7-5.0, p<10^-16^ Wald test). In comparison, wild-type patients had a median OS of 3.5 years (95% CI: 3.4-3.9 years) (Fig. 4a). The effect of mono-allelic *TP53* on OS was independent and not confounded by del(5q) (Supplementary Fig. 11). The 5-year cumulative incidence of AML transformation in mono-allelic *TP53* was 21% and 44% in the multi-hit state (HR=5.5, 95% CI: 3.1-9.6, p<10^-8^ Wald test) (Fig. 4b). Of note, the different *TP5*3-subgroups defining the multi-hit state (multiple mutations, mutation(s) and deletion or cnLOH) had equally dismal outcomes (Extended Data Fig. 5), illustrating that the various mutagenic processes leading to bi-allelic targeting of *TP53* equally shape the clinical routes of MDS patients.

**Figure 4.**
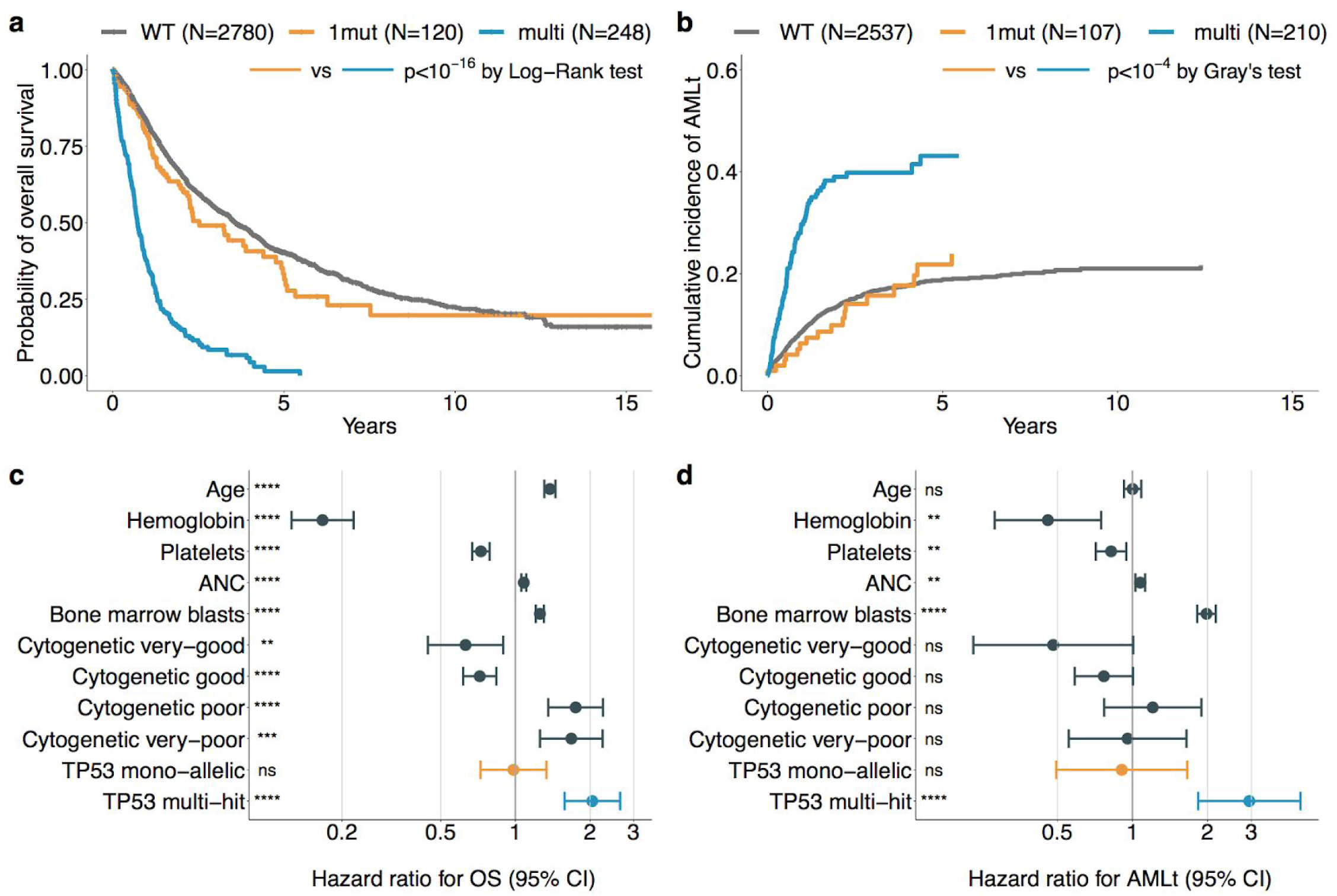
*TP53* allelic state shapes patient outcomes. Kaplan-Meier probability estimates of overall survival **(a)** and cumulative incidence of AML transformation (AMLt) **(b)** per *TP53* state of wild-type *TP53* (WT), mono-allelic *TP53* per single gene mutation (1mut) and multiple *TP53* hits (multi). **c**, Results of Cox proportional hazards regression for overall survival (OS) performed on 2,719 patients with complete data for OS and with 1,290 observed death. Explicative variables are Hemoglobin, Platelets, Absolute neutrophil count (ANC), Bone marrow blasts, Cytogenetic IPSS-R risk scores (very-good, good, intermediate is the reference, poor and very-poor) and *TP53* allelic state (mono-allelic, multi-hit and wild-type is the reference). Hemoglobin, Platelets, ANC and Bone marrow blasts are scaled by their sample mean. Age is scaled by a factor 10. The x-axis is log_10_ scaled. **d**, Results of cause-specific Cox proportional hazards regression for AML transformation (AMLt) performed on 2,464 patients with complete data for AMLt and with 411 observed transformation. Covariates are the same as in c. The x-axis is log_10_ scaled. ****p<0.0001, ***p<0.001, **p<0.01, *p<0.05 Wald test.

The OS separation of the two *TP53* states transcended disease subtypes and was significant across most WHO classes (Extended Data Fig. 6a). It was less pronounced in MDS with excess blasts and AML/AML-MRC arguably because other dominant risk factors exist besides *TP53* allelic state. Differences in outcomes between *TP53* states were also independent of IPSS-R risk groups (Extended Data Fig. 6b), and multi-hit *TP53* state identified patients with poor survival across IPSS-R strata. Notably, 10% of multi-hit patients were classified as IPSS-R good/very-good or intermediate risk. The implication of this finding is that assessment of *TP53* allelic state is critical to identify higher risk patients. In fact, multivariable Cox proportional hazards models that included *TP53* state alongside age of diagnosis, cytogenetic risk score^19^ and established predictive features identified multi-hit *TP53* as an independent predictor for the risk of death and AML transformation (HR_OS_=2.04, 95% CI: 1.6-2.6, p<10^-7^; HR_AML_=2.9, 95% CI: 1.8-4.7, p<10^-5^ Wald test), whereas mono-allelic *TP53* state did not influence OS or AML transformation compared to wild-type *TP53* (Fig. 4c-d). The same conclusion resulted from multivariable models that considered overall IPSS-R score (Supplementary Fig. 12).

Recent studies reported additive risk effects for *TP53* mutations and complex karyotype^3, 11, 12^, highlighting that each independently contribute to a patient risk. In multivariable analyses, multi-hit *TP53* state and complex karyotype, but not mono-allelic *TP53*, were independent predictors of adverse outcome (Supplementary Fig. 13). Despite the strong correlation between multi-hit *TP53* and complex karyotype, the additive risk effects remained in this setting, whereby patients with complex karyotype and multi-hit *TP53* state did worse than patients with either complex or multi-hit *TP53* (Supplementary Fig. 13b-c). In the absence of complex karyotype, mono-allelic *TP53* patients had similar survival than wild type patients while multi-hit *TP53* patients had an increased risk of death (Supplementary Fig. 13c). This emphasizes the importance of mapping *TP53* state alongside complex karyotype for accurate risk estimation.

*TP53* mutation VAF had been reported to be of prognostic significance in MDS^30^. This is likely explained by the strong correlation between high VAF, especially for values exceeding 50%, and bi-allelic targeting. However, we showed that VAF as a criterion cannot accurately capture the entire spectrum of patients with bi-allelic targeting, which includes patients with more than one *TP53* mutations in the absence of allelic imbalance and patients with subclonal cnLOH at VAF ≤50% (Fig. 1b-c and Extended Data Fig. 2). Optimal cut-point analysis^31^ identified that patients with mono-allelic *TP53* mutations and VAF>23% (n=34) had increased risk of death compared to wild-type patients (HR=2.2, 95% CI: 1.5-3.2, p<10^-3^ Wald test), whereas patients with mono-allelic *TP53* mutations and VAF≤23% (n=91) had similar OS than wild-type patients (Supplementary Fig. 14). While we may have missed a second *TP53* hit in the small subset of mono-allelic cases with VAF>23%, this shows that patients with mono-allelic mutations and high VAF should be closely monitored. Conversely, multi-hit patients had poor outcomes across all ranges of VAF, whereby even the subset of patients with low VAF≤10% (n=20) had very dismal outcome (Supplementary Fig. 14). This highlights that VAF alone is not sufficient to determine *TP53* allelic state, which requires assessment of both mutations and allelic imbalances, and that multi-hit *TP53* state identifies very high-risk patients independently of the VAF of *TP53* mutations.

### Effect of *TP53* mutation types in clinical outcomes

The emergence of data in support of DNE^18, 32^ and GOF^33,34,35^ led us to test whether outcomes differed based on the nature of the underlying lesion, i.e., missense, truncated or hotspot mutations. In the multi-hit state, no differences were observed on genome instability levels (Extended Data Fig. 7) and outcomes (Extended Data Fig. 7 and Supplementary Fig. 15a-b) across mutation types, showcasing that it is the loss of both wild-type copies of *TP53* that drive the dismal outcomes of *TP53*-mutated MDS patients rather than the underlying mutation types.

In the mono-allelic state, missense mutations in the DBD as a whole had no effect on patient outcomes compared to wild-type *TP53*. However, there was an increased risk of death of mono-allelic *TP53* patients with hotspot mutations (R175, R248) compared to wild-type patients (HR=1.7, 95% CI: 1.1-2.8, p=0.02 Wald test, Supplementary Fig. 15c-d). This is consistent with either DNE or GOF of the hotspot mutant proteins with increased selection of these mutated residues. But this observation was not uniform across all mono-allelic missense mutations, suggesting that the putative DNE^18^ may not be equivalent across DBD mutations. Definitive conclusions on the possible non-equivalence of mono-allelic missense mutations warrant evaluation in larger datasets and functional studies that extend beyond the mutation hotspots.

### Consequences of *TP53* allelic state in therapy-related MDS

Our cohort included 229 cases with therapy-related MDS (t-MDS), which were enriched^8, 13^ for *TP53*-mutated patients relative to de-novo MDS (18% vs. 6%, OR=3.3, p<10^-11^ Fisher exact test). The *TP53*-mutated t-MDS patients had a higher proportion of multiple hits compared to *TP53*-mutated de-novo patients (84% vs. 65%, OR=2.8, p=0.002 Fisher exact test). Comparison of genome profiles (Supplementary Fig. 16) and clinical outcomes (Fig. 5a) between *TP53* allelic states reiterated observations in de-novo MDS. *TP53*-mutant t-MDS is considered one of the most lethal malignancies with limited treatment options^7^, yet mono-allelic *TP53* had lower risk of death compared to multi-hit *TP53* even in the t-MDS setting (HR=0.39, 95% CI: 0.15-1.0, p=0.05 Wald test).

**Figure 5.**
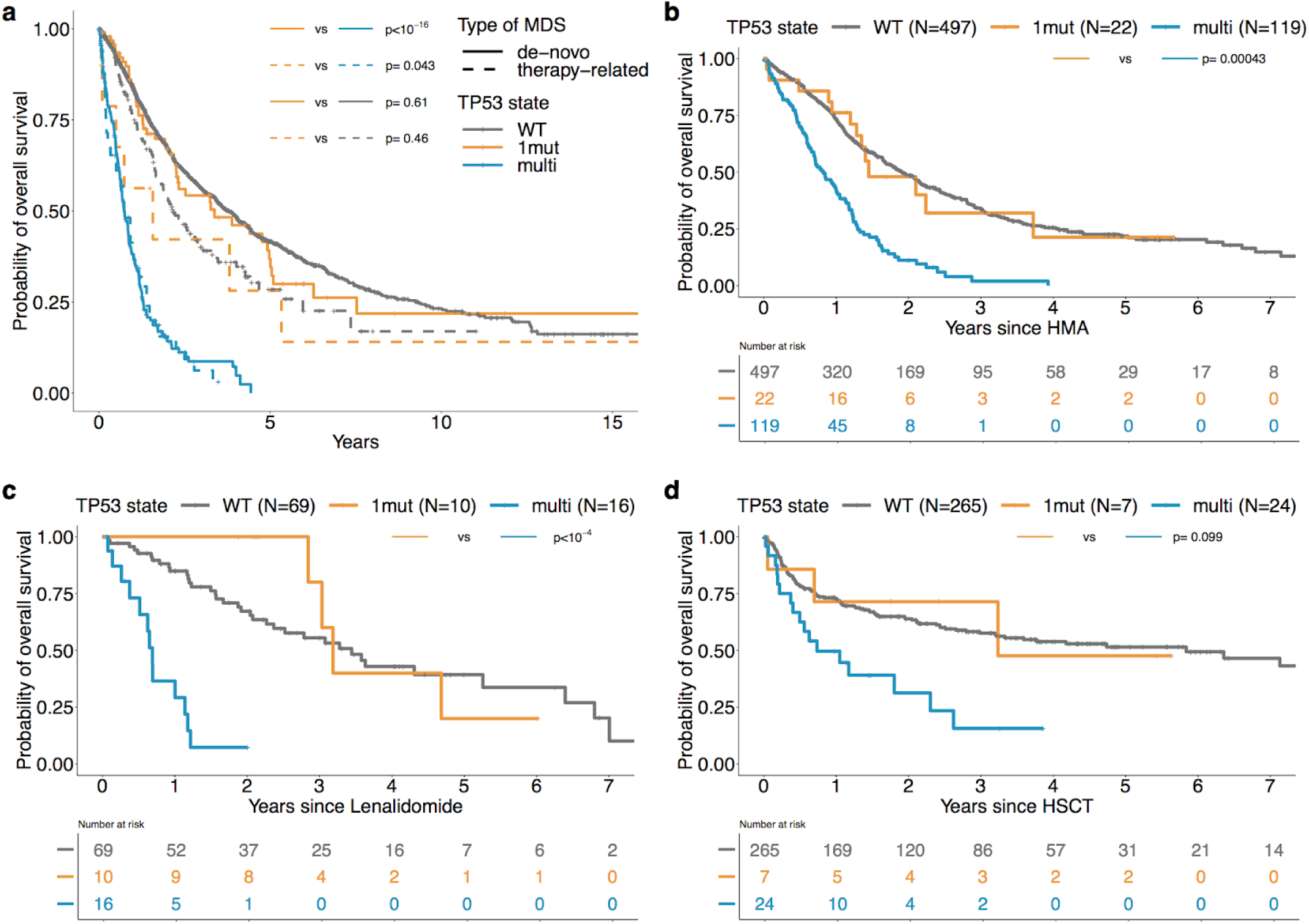
*TP53* allelic state demarcates distinct outcomes in therapy-related MDS and on different therapies. **a**, Kaplan-Meier probability estimates of overall survival per allelic state of wild-type *TP53* (WT), mono-allelic *TP53* per single gene mutation (1mut) and multiple *TP53* hits (multi); and across type of MDS, i.e., de-novo MDS (solid line) or therapy-related MDS (dashed line). Within the therapy-related cases, 10 had a mono-allelic *TP53* mutation (dashed orange line), 52 were multi-hit *TP53* (dashed blue line) and 162 were *TP53* wild-type (dashed grey line). **b-c-d**, Kaplan-Meier probability estimates of overall survival post start of hypomethylating agent (HMA) treatment **(b)** start of Lenalidomide treatment for patients with del(5q) **(c)** hematopoietic stem cell transplantation (HSCT) **(d)** per allelic state of wild-type *TP53* (WT), mono-allelic *TP53* (1mut) and multiple *TP53* hits (multi). In b, c, and d, overall survival was measured from the time of treatment start or HSCT to the time of death from any cause. Patients alive at the last follow-up date were censored at that time. Annotated p-values are from the log-rank test.

### *TP53* allelic state and disease progression

We analyzed serial data from 12 MDS patients of an independent cohort collected from the diagnostic service of St James’s University Hospital (Leeds, United Kingdom)^36, 37^ who progressed to AML with a *TP53* mutation in either disease phase (Supplementary Fig. 17). We found a preponderance of two *TP53* hits at the time of MDS diagnosis (7/12 cases, with a median of 4 months to AML progression) (Supplementary Fig. 17a-g). In 3 patients, bi-allelic targeting occurred during disease progression with evidence of inter-clonal competition and attainment of clonal dominance for the *TP53* clone (Supplementary Fig. 17h-i). The remaining two cases that progressed with a mono-allelic *TP53* mutation had other high-risk mutations in *RUNX1* and *KRAS* or in *CBL* (Supplementary Fig. 17k-l). These data provided further evidence that bi-allelic alteration of *TP53* is a potent driver of disease progression and underscored the importance of assessing *TP53* allelic state at diagnosis and for disease surveillance.

### Validation cohort

We tested our findings in 1,120 MDS patients with comparable molecular annotations. We validated the representation of *TP53* allelic states (Supplementary Fig. S18), genome stability profiles (Supplementary Fig. S19) and differences in clinical phenotypes (Supplementary Fig. S20). Our validation cohort was enriched for higher risk disease subtypes compared to our study cohort (Extended Data Table 1 and 2). Overall, multi-hit patients had significantly increased risk of death than mono-allelic patients (Supplementary Fig. S20e). Within lower-risk disease subgroups, OS of mono-allelic *TP53* patients was similar to that of wild-type patients (Supplementary Fig. S20f).

### *TP53* allelic state on treatment response

Recent studies report poor responses to lenalidomide^10^ and HSCT^8, 9^ for *TP53*-mutated patients, and marked but transient response to HMA^38^. We conducted an exploratory analysis of overall survival per *TP53* state of patients that received hypomethylating agent (HMA) (Fig. 5b), lenalidomide on the subset with del(5q) (Fig. 5c) and following HSCT (Fig. 5d). On HMA and lenalidomide, patients with mono-allelic *TP53* mutations had evidence of longer survival compared to multi-hit patients (Fig. 5b-c). The analysis of our HSCT cohort was limited due to its size, yet we observed a trend for improved survival of mono-allelic patients compared to multi-hit patients following HSCT (Fig. 5d). These observations highlight the importance of mapping allelic state in future correlative studies of *TP53* response to therapy.

## DISCUSSION

The increasing prevalence of molecular profiling in clinical practice calls for an improved mapping of genotype features to clinical outcomes, in order to identify meaningful biomarkers and institute precision medicine practices^39^. Beyond biomarker validation, the delivery of precision medicine practices is increasingly reliant upon precision diagnostics. This study unraveled the distinct effects of the allelic states of *TP53*, the most frequently mutated gene in cancer^1, 2^, with clinical implications for diagnosis, disease surveillance and risk stratification.

We developed a novel framework for the ascertainment of *TP53* mutations, focal deletions and cnLOH from unmatched custom capture sequencing data. Our copy-number tools CNACS is accessible from an open-source software development platform (https://github.com/papaemmelab/toil_cnacs). Through integrative analyses of the mutagenic processes targeting *TP53*, coupled with large sample size and robust clinical annotation, we were able to accurately characterize *TP53* allelic states and evaluate their clinical implications in MDS.

*TP53* is universally considered as an adverse prognostic biomarker associated with genome instability, treatment resistance, disease progression and dismal outcomes^16, 40^. We provided strong and definitive evidence that the multi-hit *TP53* state in MDS, not the bare presence of any *TP53* mutation, underlies these associations. It is therefore critical to accurately assess *TP53* allelic state in the diagnostic workup of MDS patients. Combining information from the number of *TP53* mutations, CBA, VAF and LOH status derived from NGS-based copy-number analysis or SNP arrays would allow clinical laboratories to discriminate between the vast majority of bi-allelic and mono-allelic *TP53* mutations. Fig. 6 suggests an easily implementable workflow for the assessment of *TP53* allelic state in routine clinical practice. We propose that bi-allelic *TP53* state should be distinguished from mono-allelic *TP53* mutations in future revisions of the IPSS-R and correlative studies of treatment response. This is meaningful for clinical practice as one in three *TP53*-mutated patients is mono-allelic. In our cohort, mono-allelic patients did not differ from *TP53* wild-type patients with regards to genome stability, response to therapy, overall survival and progression to AML. Although *TP53* is the most scrutinized cancer gene, our study materializes to our knowledge the first assessment of the impact of the allelic state of *TP53* on disease biology and clinical outcomes in large MDS patient cohorts. Given the importance of *TP53* in cancer, these findings warrant further investigation across cancer indications.

**Figure 6.**
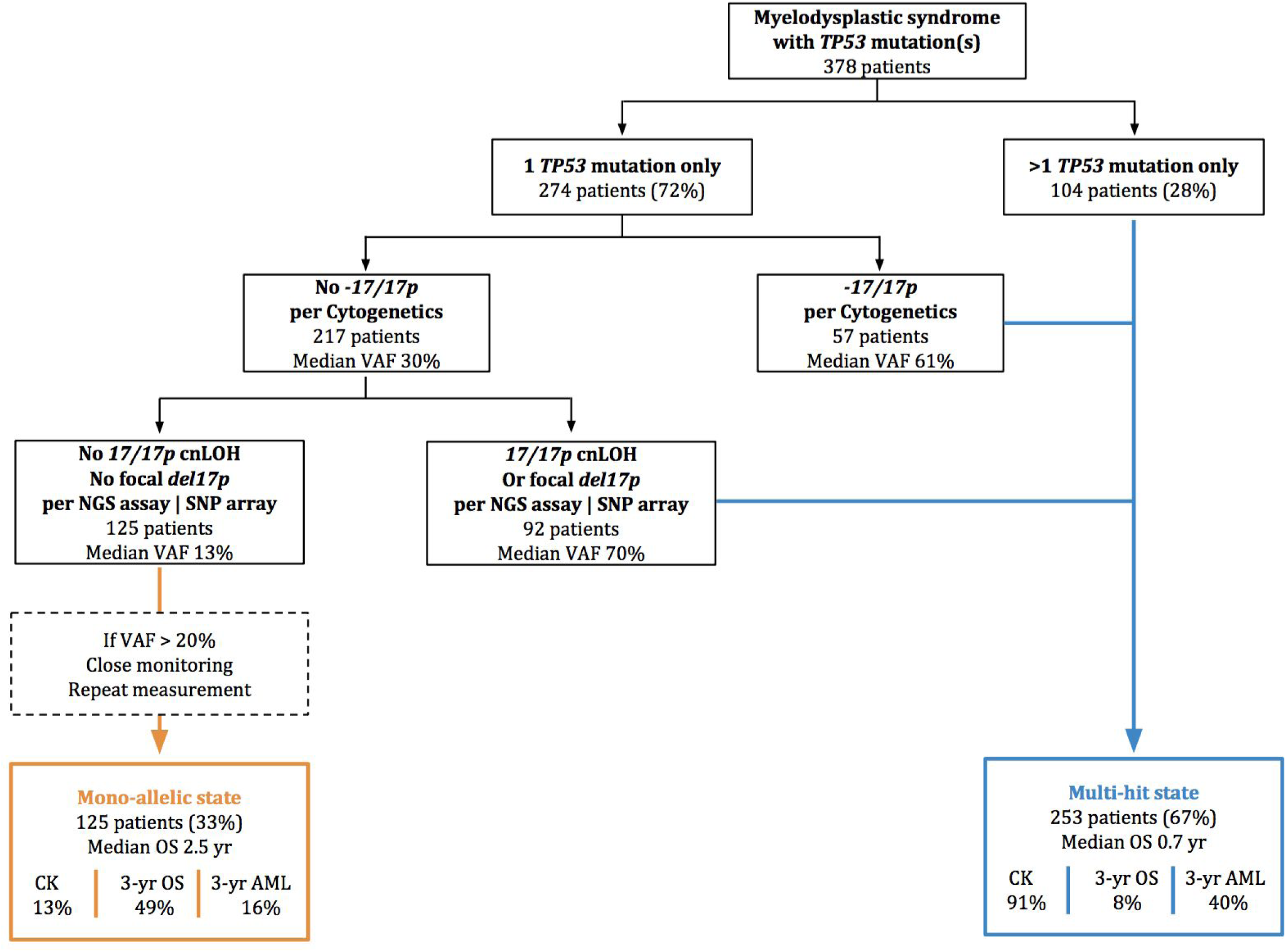
Clinical workflow for the assessment of *TP53* allelic state. Schematic of a simple clinical workflow based on the number of *TP53* mutations, the presence or absence of deletion 17p per cytogenetic analysis, and the presence or absence of cnLOH at 17p or focal deletion per NGS based assay or SNP array. Mutations were considered if VAF≥2%. VAF: variant allele frequency; CK: complex karyotype; OS: overall survival; AML: transformation to acute myeloid leukemia.

## METHODS

### Patient samples

The IWG-PM cohort originated from 24 MDS centers (Supplementary Table 1) that contributed peri-diagnosis MDS, MDS/MPN and AML/AML-MRC patient samples to the study. Upon quality control (Supplementary Fig. 1), 3,324 samples were included in the study (Extended Data Table 1). The source for genomic DNA was bone marrow or peripheral blood. The median time from diagnosis to sampling was 0 days (1st quartile: 0 days, 3rd quartile: 113 days). The validation cohort consisted of 1,120 samples from the Japanese MDS consortium (Extended Data Table 2). Samples were obtained with informed consent in accordance with the Declaration of Helsinki and appropriate Ethics Committee approvals.

### Clinical data

Diagnostic clinical variables were provided by the contributing centers and curated to ensure uniformity of metrics across centers and countries. Clinical variables included i) Sex ii) Age at diagnosis iii) WHO disease subtype iv) MDS type i.e., de-novo, secondary or therapy-related MDS v) Differential blood counts to include hemoglobin, platelets, white blood cell, neutrophil and monocyte vi) Percentage of bone marrow and peripheral blood blasts vii) Cytogenetic data and viii) Risk score as per the IPSS-R^14^. Clinical outcomes included the time of death from any cause or last follow-up from sample collection, and the time of AML transformation or last follow-up from sample collection. Detailed cohort characteristics are provided in Extended Data Table 1 and 2.

#### Cytogenetic data

CBA data were available for 2,931 patients and karyotypes were described in accordance to the International System for Human Cytogenetic Nomenclature^41^. CBA data were risk stratified according to the IPSS-R guidelines^19^ using both algorithmic classification and manual classification by an expert panel of cytogeneticists.

#### WHO subtypes

Contributing centers provided for the vast majority disease classification as per WHO 2008. Pathology review was performed uniformly on the entire cohort, to ensure concordance between disease classification and diagnostic variables, and to update the classification as per WHO 2016.

#### IPSS-R risk scores

IPSS-R risk scores were uniformly calculated based on the IPSS-R cytogenetic risk scores and on the values for hemoglobin, platelets, absolute neutrophil count and percentage of bone marrow blasts.

### Targeted sequencing

#### Panel design

The panel used for targeted sequencing included 1,118 genome wide single nucleotide polymorphism (SNP) probes for copy number analysis, with on average one SNP probe every 3Mb. Bait tiling was conducted at 2x. Baits were designed to span all exonic regions of *TP53* across all transcripts, as described in RefSeq (NM_001276761, NM_001276695, NM_001126114, NM_00112611), and included 20bp intronic flanking regions.

#### Library preparation and sequencing

For library construction, 11-800ng of genomic DNA was used using the KAPA Hyper Prep Kit (Kapa Biosystems KK8504) with 7-12 cycles of PCR. After sample barcoding, 10-1610ng of each library were pooled and captured by hybridization. Captured pools were sequenced with paired-end Illumina HiSeq at a median coverage of 730x per sample (range 127-2480x). Read length was 100bp or 125bp.

We also sequenced 48 samples on the panel, with the same sequencing conditions as the tumor samples, from young individuals who did not have hematological disease; to help further filtering of sequencing artefacts and germline SNPs.

Sequencing was performed in an unmatched setting i.e., without a matched normal tissue control per patient, so that variants had to be curated accordingly (see section “*TP53* variant annotation” below).

#### Alignment

Raw sequence data were aligned to the human genome (NCBI build 37) using BWA^42^ version 0.7.17. PCR duplicate reads were marked with Picard tools (https://broadinstitute.github.io/picard/) version 2.18.2. For alignment, we used the pcap-core dockerized pipeline version 4.2.1 available at https://github.com/cancerit/PCAP-core/wiki/Scripts-Reference-implementations.

### Sample quality control

Quality control (QC) of the fastq data and bam data were performed with FastQC (http://www.bioinformatics.babraham.ac.uk/projects/fastqc/) version 0.11.5 and Picard tools respectively.

In addition, a number of downstream QC steps were performed, to include:

- Fingerprinting, i.e., evaluation of the similarity between all pairs of samples based on the respective genotype on 1,118 SNPs. Duplicate samples were excluded from the study.
- Evaluation of concordance between the patient sex from the clinical data and the coverage on the sex chromosomes. Discordant cases were discussed with the contributed centers to rule out patients with Klinefelter syndrome and filter out erroneous samples appropriately.
- Evaluation of concordance between CBA data and NGS derived copy-number profiles (see section “Copy number analysis” below). A typical discordant case is a case where CBA reports a given deletion or gain in a high number of metaphases and NGS profile clearly shows other abnormalities but not the one reported by CBA. All discordant cases were reviewed by a panel of experts through the IWG cytogenetic committee.

Finally, samples that passed QC but were found not to be treatment naive i.e., the patients received disease modifying treatment before sample collection were excluded from the study. Supplementary Fig. 1 summarizes the QC workflow.

### *TP53* variant calling

Variants in *TP53* were called using a combination of variant callers. For single nucleotide variants (SNVs), we used Caveman (http://cancerit.github.io/CaVEMan/) version 1.7.4, Mutect^43^ version 4.0.1.2 and Strelka^44^ version 2.9.1. For small insertions and deletions, we used Pindel^45^ version 1.5.4, Mutect version 4.0.1.2 and Strelka version 2.9.1. VAFs were uniformly reported across all called variants using a realignment procedure (https://github.com/cancerit/vafCorrect).

Likely artefact variants were filtered out based on:

- The number of callers calling a given variant and the combination of filters from the triple callers.
- Variants with VAF<2%, less than 20 total reads or less than 5 mutant supporting reads were excluded.
- Recurrence and VAF distribution of the called variants on a panel of normals.
- Off-target variants, i.e., variants called outside of the panel target regions were not considered.

### *TP53* variant annotation

All called variants were annotated with VAGrENT (https://github.com/cancerit/VAGrENT) version 3.3.0 and Ensembl-VEP (https://github.com/Ensembl/ensembl-vep) with Ensembl version 91 and VEP release 94.5.

After pre-filtering of artifactual variants, likely germline SNPs were filtered out by consideration of:

- VAF density of variants consistent with germline SNP.
- Presence in the Genome Aggregation Database (gnomAD)^46^.
- Recurrence in panel of normals.

All remaining likely somatic *TP53* variants were manually inspected with the Integrative Genomics Viewer (IGV)^47^ to rule out residual artefacts.

From the list of likely somatic *TP53* variants, putative oncogenic variants were distinguished from variants of unknown significance (VUS) based on:

- The inferred consequence of a mutation; where nonsense and splice SNVs, and frameshift insertions and deletions were considered oncogenic.
- Recurrence in the Catalogue Of Somatic Mutations in Cancer (COSMIC)^24^, in myeloid disease samples registered in cBioPortal^24, 48^ or in the study dataset.
- Presence in pan-cancer hotspot analysis as described in^49^ and^50^.
- Annotation in the human variation database ClinVar^28^.
- Annotation in the precision oncology knowledge database OncoKB^23^.
- Functional annotation in the International Agency for Research on Cancer (IARC) *TP53* database^25^.
- *TP53* functional classification prediction scores using PHANTM^51^.
- Recurrence with somatic presentation in a set of in-house data derived from >6,000 myeloid neoplasms^12, 21, 22^.

### Copy number analysis

We assessed chromosomal alterations based on NGS sequencing data using CNACS^9^. CNACS enables the detection of arm level and focal copy-numbers changes as well as regions of cnLOH. CNACS has been optimized to run in the unmatch setting and uses a panel of normals for calibration. CNACS^9^ is available as a python toil workflow engine at https://github.com/papaemmelab/toil_cnacs, where release v0.2.0 was used in this study.

Supplementary Fig. 2 provides examples of characterization of allelic imbalances (gains, deletions and regions of cnLOH) using CNACS, with concordant copy-number change findings between CBA and CNACS, focal deletions exclusively detected with CNACS and, as expected, regions of cnLOH exclusively detected by CNACS. Supplementary Fig 3a-b provides a genome-wide characterization of allelic imbalances on 2,931 MDS patients and compares the levels of detection from CBA and CNACS. Note that for genome-wide analysis, we restricted the CNACS gain, deletion or cnLOH segments to be bigger than 3Mb. Supplementary Fig 4 provides examples of characterization of allelic imbalances by CNACS and SNP arrays on 21 selected samples, with extremely concordant findings between the two assays.

In addition to CNACS, we also run CNVkit^52^ version 0.9.6 on the study cohort. CNVkit does not infer allele specific copy-numbers, so that it does not allow to mark regions of cnLOH, but it estimates copy-number changes. The integration of two copy-number tools allowed to increase specificity and sensitivity of the calling.

On 2,931 patients with CBA data, we performed a detailed comparison of CBA and CNACS results (Supplementary Fig. 3). Along the annotation of regions of cnLOH, we supplemented the presence of copy-number changes on those patients when it was clear on the NGS results but missed by CBA. For the 393 patients with missing CBA data, we used the NGS results to annotate copy-number changes. As our NGS assay did not allow to detect translocations, inversions, whole genome amplification and the presence of marker or ring chromosomes, those specific alterations were statistically imputed from other molecular markers on the 393 patients with missing CBA.

#### Complex Karyotype

From the 2,931 patients with CBA data, 310 had a complex karyotype according to the CBA results, where complex karyotype was defined as 3 or more independent chromosomal abnormalities. Within the 2,931 patients with CBA data, CNACS results helped to identify complex karyotype in an additional 15 patients. Within the 393 cases with missing CBA data, 13 had a complex karyotype according to NGS copy-number profiles (Supplementary Fig. 3c). Overall, 329 patients had complex karyotype representing 10% of the study cohort.

### Survival analysis

All statistical analyses were conducted using the R statistical platform (R Core Team 2019) (https://www.r-project.org/).

#### Overall survival

OS was measured from the time of sample collection to the time of death from any cause. Patients alive at the last follow-up date were censored at that time. Survival probabilities over time were estimated using Kaplan-Meier methodology, and comparisons of survival across *TP53*-mutant subgroups were conducted using the logrank test.

Multivariable models of OS were performed with Cox proportional hazards regressions. Hazard ratios and 95% confidence intervals were reported for the covariates along the p-values from the Wald test. Covariates included in the multivariable model of OS shown in Fig. 3c were age, hemoglobin, platelets, absolute neutrophil count (ANC), bone marrow blasts, cytogenetic risk group and *TP53* allelic state. Hemoglobin, platelets, ANC and bone marrow blasts were treated as continuous variables and were scaled by their sample mean. Age was treated as a continuous variable and was scaled by a factor 10. Cytogenetic risk group was treated as a categorical variable with the intermediate risk group as the reference group. *TP53* allelic state was treated as a categorical variable with the wild-type state as the reference group relative to the mono-allelic and the multi-hit groups. Note that those covariates correspond to all covariates included in the age-adjusted IPPS-R model in addition to *TP53* allelic state.

#### AML transformation

In univariate analysis of AML transformation (AMLt), time to AMLt was measured from the time of sample collection to the time of transformation, with death without transformation treated as a competing risk. Patients alive without AMLt at the last contact date were censored at that time. Cumulative incidence functions were used to estimate the incidence of AMLt and comparisons of cumulative incidence function across *TP53*-mutant subgroups were conducted using the Gray’s test.

Multivariable models of AMLt were performed using cause-specific Cox proportional hazards regressions, where patients who did not transform but died were censored at the time of death. Hazard ratios and 95% confidence intervals were reported for the covariates along the p-values from the Wald test. Covariates included in the multivariable model of AMLt shown in Fig. 3d were the same as the ones included in the model of OS described above.

## AUTHOR CONTRIBUTIONS

E.B. and E.P. designed the study. E.B., Y.N. and T.Y. performed statistical analysis. S.D. and E.P. supervised statistical analysis. L.M., B.L.E, R.B., P.L.G., M. Cazzola, E.H-L., S.O. and E.P. supervised research. P.L.G. and E.P. coordinated the study. L.M., F.S., C.A.C., M. Creignou, U.G., A.A.L., M.J., M.T., O.K., M.Y.F., F.T., R.F.P., V.S., I.K., J.B., F.P.S.S., S.K., T.I., T.H., A.K.T., T.K., C.P., V.M.K., M.R.S., M.B., C.G., L.P., L.A., M.G.D.P., P.F., A.P., U.P., M.H., P.V., S.C., Y.M., C.F., M.T.V., L-Y.S., M.F., J.H.J., J.C., Y.A., N.G., M. Cazzola, E.H-L. and S.O. provided clinical data and DNA specimens. E.B., Y.W., M.P. and E.P. coordinated sample acquisition. A.V. and K.V. performed sample preparation and sequencing. E.B., R.P.H., H.T. and M. Creignou curated clinical data. R.P.H. and J.M.B. performed pathology review. E.B. and H.T. processed cytogenetic data. F.S., D.H. and J.S. performed cytogenetic review. E.B., Y.N., J.S.M-M, T.Y., A.S. and G.G. performed bioinformatic analysis. J.S.M-M, M.F.L., J.E.A. and J.Z. supported sequence data pipelines. Y.S. and R.S. developed copy-number algorithm CNACS. M.F.L. generated copy-number profiles. Y.Z. performed SNP array analysis. E.B. and Y.N. prepared figures and tables. E.B., S.O. and E.P. wrote the manuscript. All authors reviewed the manuscript during its preparation.

## ACKNOWLEDGEMENTS

Supported in part by grants from Celgene Corporation through and MDS Foundation, Inc, Yardville, NJ; J.B. and A.P. acknowledge funding from Bloodwise grant 13042; P.V. was supported by the Austrian Science Fund (FWF) grant F4704-B20; M.Y. F. was supported by Italian MIUR-PRIN grants; L.M. was supported by the Associazione Italiana per la Ricerca sul Cancro (AIRC, Milan, Italy) 5×1000 project #21267 and IG #20125; M.T.V. is supported by AIRC 5permille N. 21267 and patients are recruited through the GROM-L clinical network; E.B. was supported by the Francois Wallace Monahan Fellowship; E.P. is a Josie Robertson Investigator and is supported by the European Hematology Association, American Society of Hematology, Gabrielle’s Angels Foundation, V Foundation and The Geoffrey Beene Foundation. We thank Tracey Iraca for logistic support.

## EXTENDED DATA LEGENDS

**Extended Data Figure 1.**
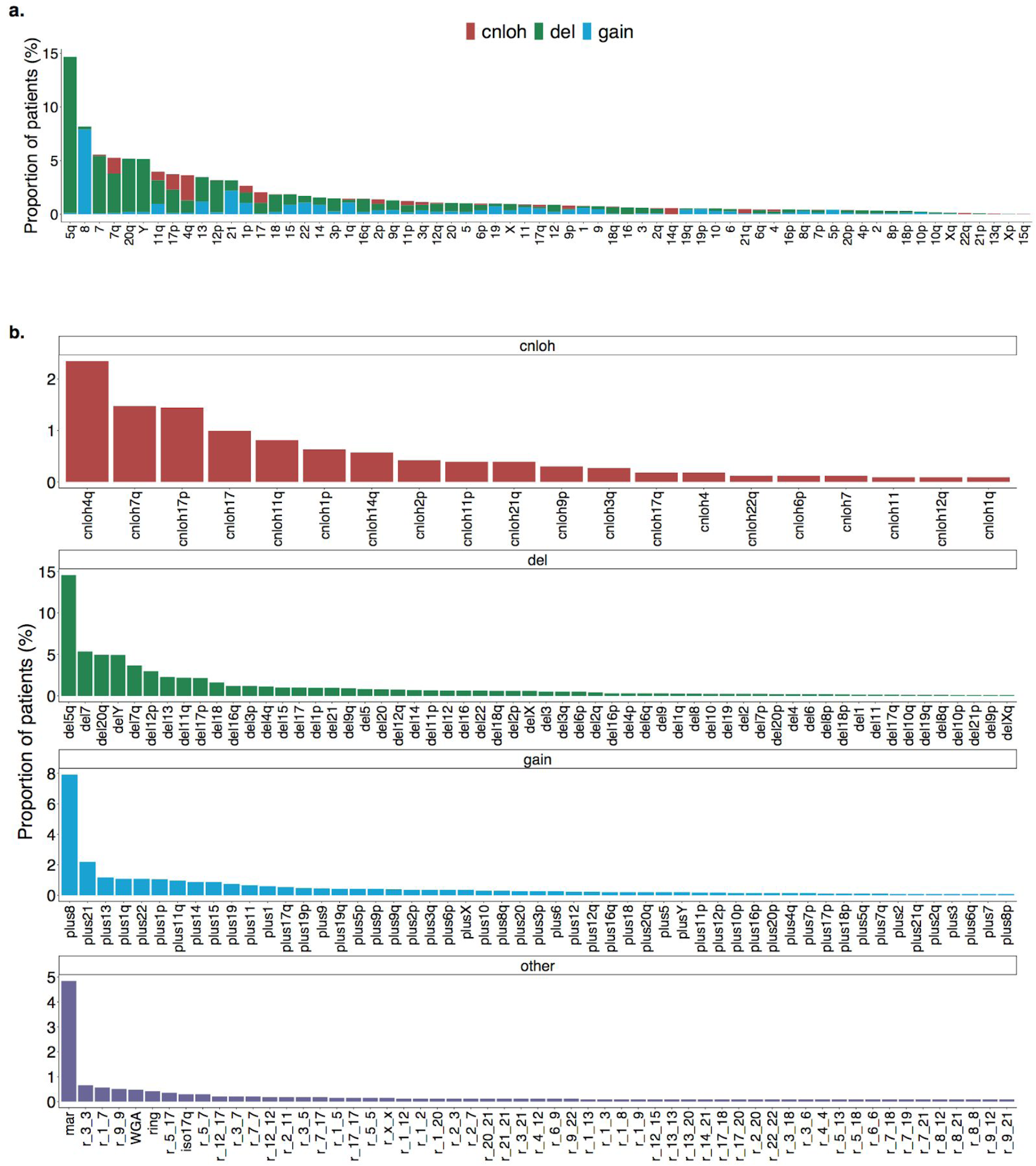
Landscape of chromosomal aberrations in MDS. **a**. Landscape of chromosomal arm-level aberrations across 3,324 patients. Aberrations include copy-neutral loss-of-heterozygosity (cnloh), deletion (del) and gain. The x-axis indicates chromosome arms or entire chromosomes affected by aberrations. Aberrations were assessed using the integration of conventional G-banding analysis (CBA) data and NGS derived copy-number profiles. NGS aberrant segments were restricted to segments larger than 3 megabases. **b**. Frequency distribution of chromosomal aberrations across 3,324 patients ordered by type of aberrations. First top three plots represent arm-level copy-neutral loss-of-heterozygosity (cnloh), deletion (del) and gain. Fourth bottom plot represents other types of aberrations to include the presence of marker chromosome (mar), rearrangements where r_i_j denotes a rearrangement between chromosome i and j, isochromosome 17q (iso17q), whole genome amplification (WGA) and presence of ring chromosome (ring). All aberrations observed in more than 2 patients are depicted. Of note, cnloh is detectable with NGS but not with CBA. On the opposite, rearrangements, presence of marker or ring chromosome and WGA were only assessed from CBA data. On the 393 cases with missing CBA data, those specific aberrations were imputed from other molecular markers.

**Extended Data Figure 2.**
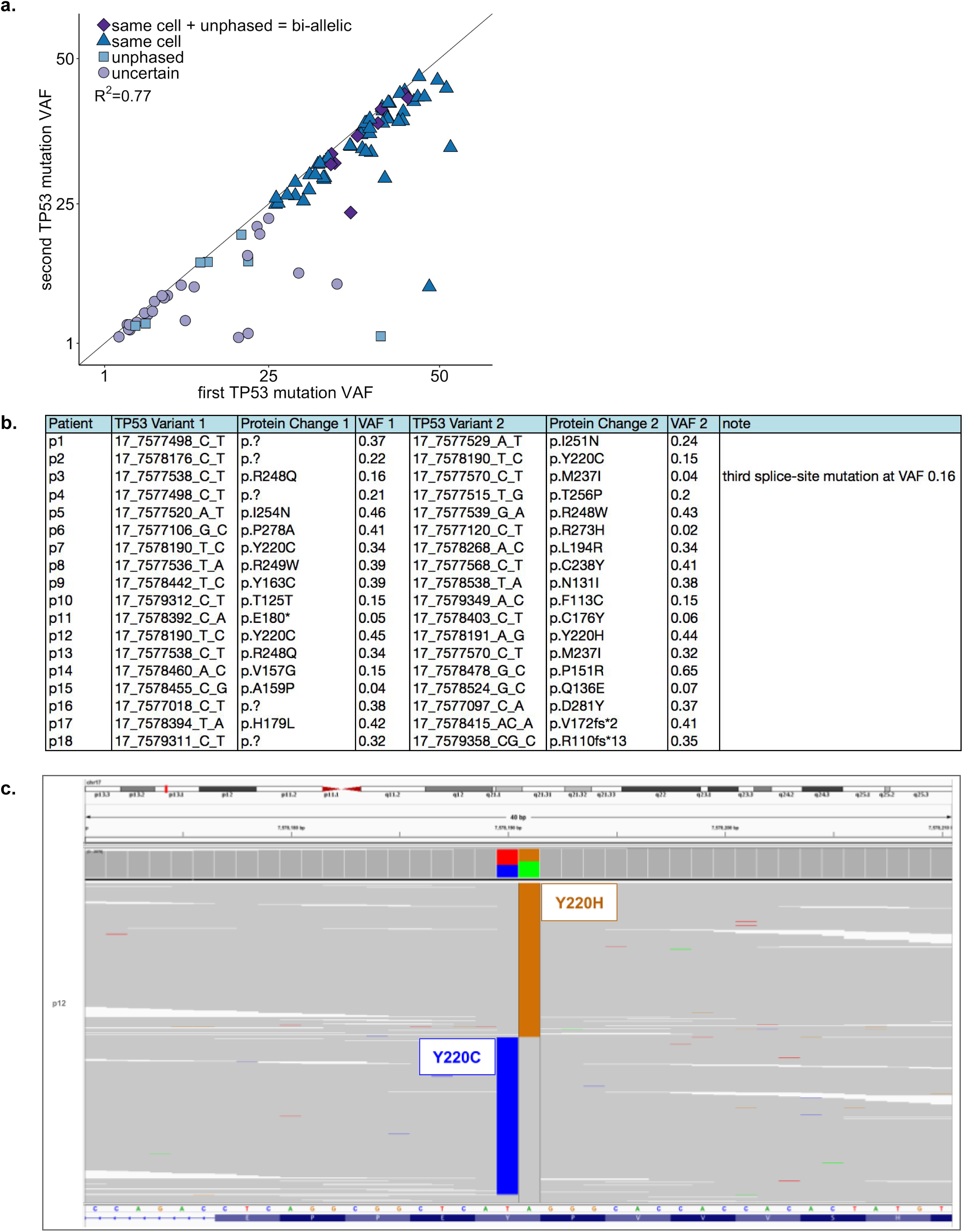
Evidence of bi-allelic *TP53* targeting in the cases with multiple *TP53* mutations. **a**. Scatter plot of the two maximum *TP53* variant allele frequency (VAF) values from the cases with multiple *TP53* mutations and no copy-neutral loss-of-heterozygosity or deletion (N=90). Points are annotated according to the level of information of the mutation pairs. If the sum of the two VAFs exceeded 50%, the mutations were considered to be in the same cells, which happened in 67% (n=60) of the cases (triangle and diamond points). In some specific cases where the genomic distance between two mutations was smaller than the read length, it was possible to phase the mutations. In the 18 cases where possible to assess, mutations were all observed to be unphased, i.e., in trans (square and diamond points). Within those 18 pairs of unphased mutations, 10 pairs had a sum of VAFs above 50%, i.e., mutations were necessarily on different alleles and in the same cells, implying bi-allelic targeting (diamond points). **b**. Table of pairs of *TP53* mutations from the same patients that could be phased. All pairs were in trans, i.e., mutations were supported by different alleles. **c**. Representative IGV example of unphased mutations (patient p12 from table b.).

**Extended Data Figure 3.**
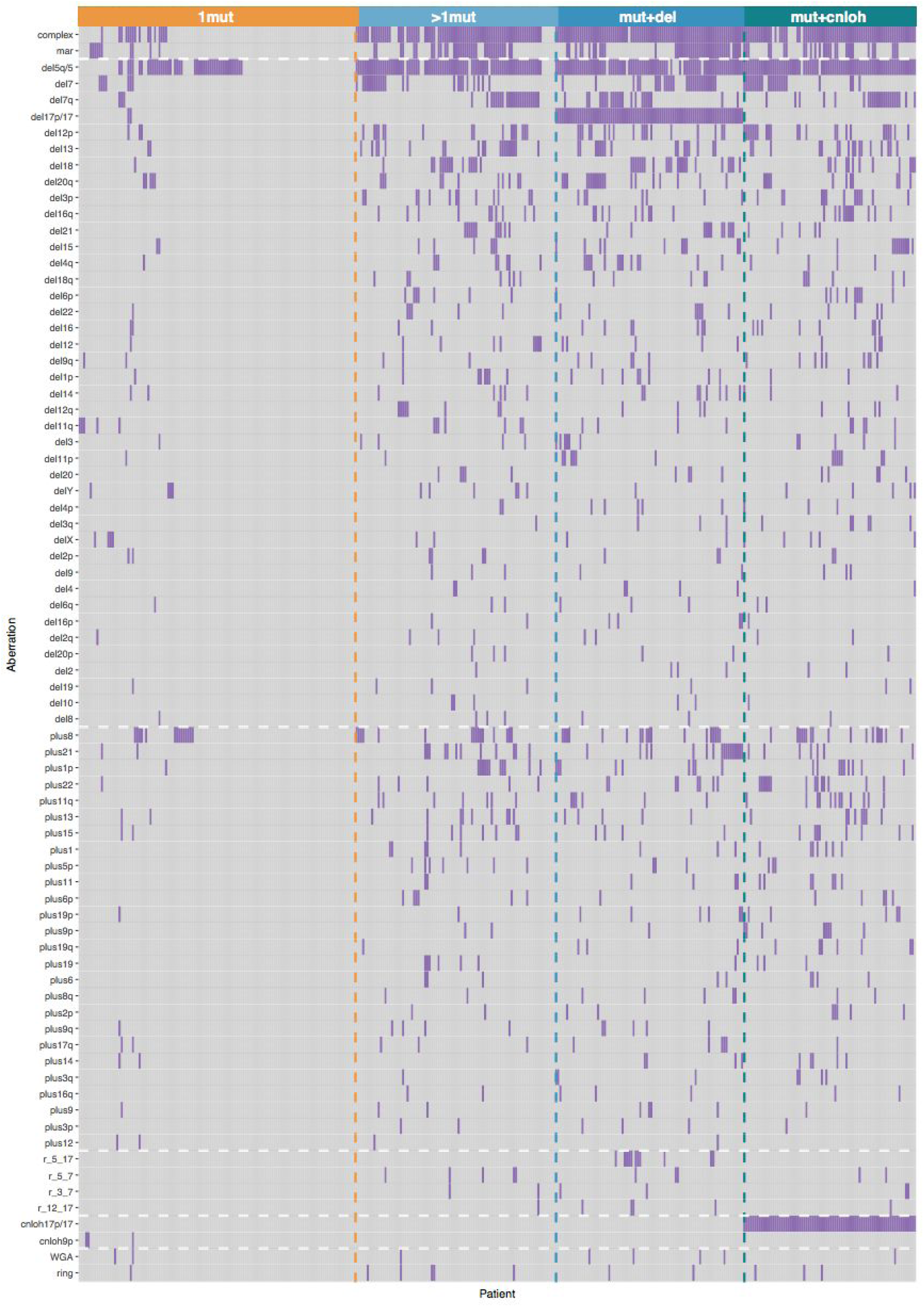
Heatmap of chromosomal aberrations per *TP53* allelic state. Each column represents a patient from the *TP53* subgroups of single gene mutation (top orange band, 1mut), multiple mutations (top light blue band, >1mut), mutation(s) and deletion (top blue band, mut+del) and mutation(s) and copy-neutral loss of heterozygosity (top dark blue band, mut+cnloh). Aberrations observed at a frequency higher than 2% in either mono-allelic or multi-hit *TP53* state are depicted on the y-axis. Aberrations include from top to bottom the annotation of complex karyotype (complex), the presence of marker chromosome (mar), deletion (del), gain (plus), rearrangement (with r_i_j rearrangement between chromosome i and j), copy-neutral loss of heterozygosity (cnloh), whole genome amplification (WGA) and the presence of ring chromosome (ring). Note that the deletions of 17p of two cases from the 1mut *TP53* subgroup did not affect the *TP53* locus.

**Extended Data Figure 4.**
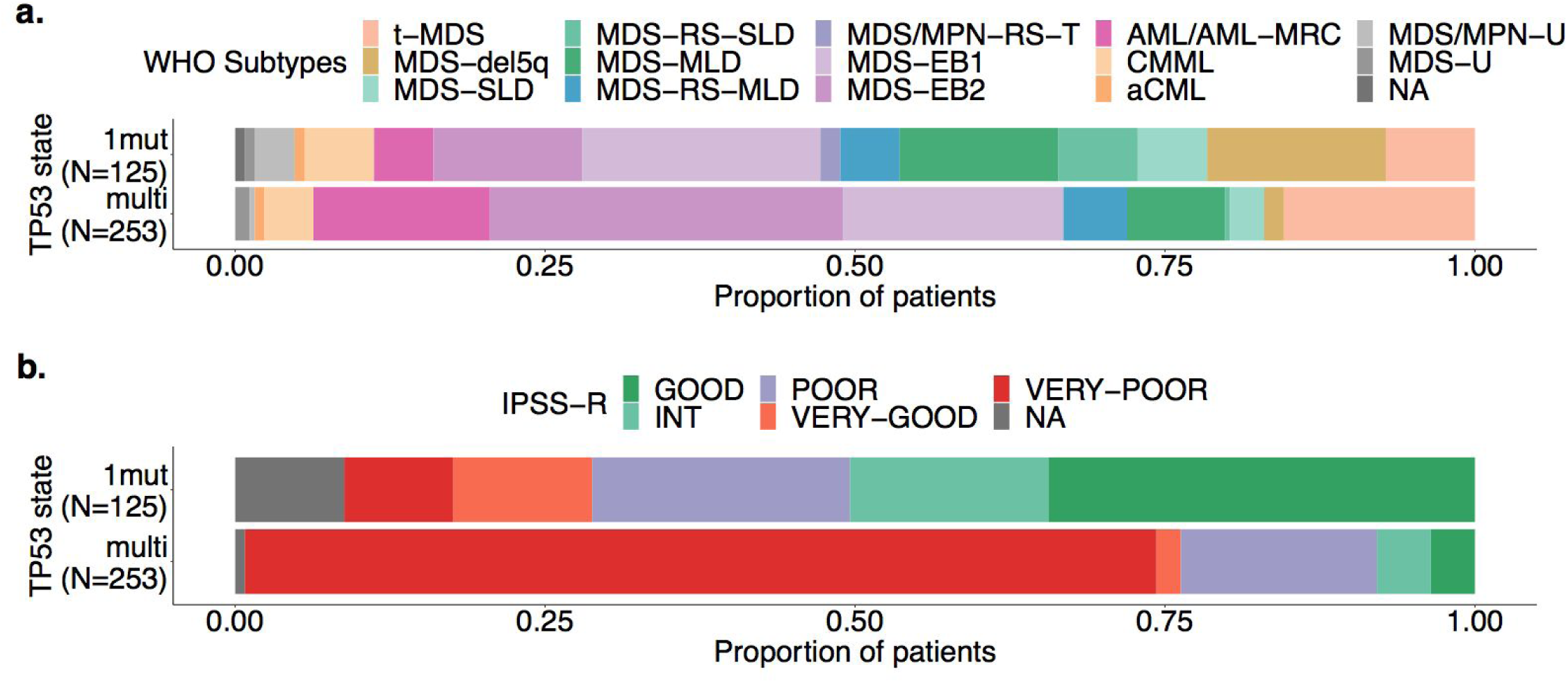
Representation of WHO subtypes and IPSS-R risk groups per *TP53* allelic state. **a**. Proportion of WHO subtypes per *TP53* state of single gene mutation (1mut) and multiple hits (multi). t-MDS: therapy-related MDS; SLD: single lineage dysplasia; RS: ring sideroblast; MLD: multiple lineage dysplasia; EB: excess blasts; AML-MRC: AML with myelodysplasia-related changes; U: unclassified. Compared to cases with single gene mutation (1mut), multi-hit *TP53* was enriched for t-MDS (21% vs. 8%, OR=2.9, p=0.002 Fisher exact test) and MDS-EB2 (31% vs. 13%, OR=3.1, p<10^-4^). Contrarily, mono-allelic *TP53* (1mut) was enriched for MDS-del5q (15% vs. 2%, OR=8.4, p<10^-5^). **b**. Proportion of IPSS-R risk groups per *TP53* state of single gene mutation (1mut) and multiple hits (multi). Multi-hit *TP53* was strongly enriched for the very-poor category compared to mono-allelic *TP53* state (74% vs. 9%, OR=27, p<10^-16^).

**Extended Data Figure 5.**
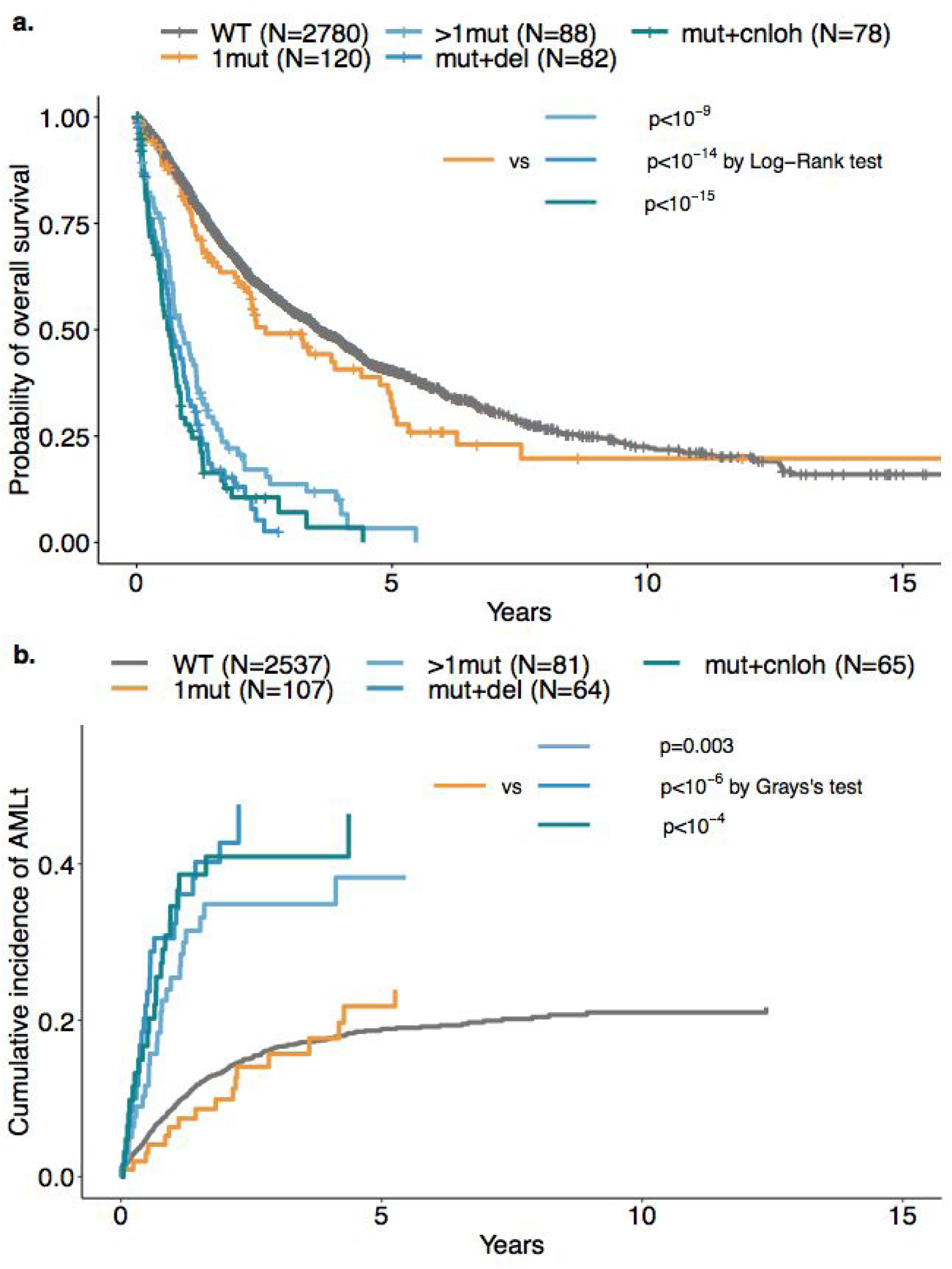
Outcomes across *TP53* subgroups. **a**. Kaplan-Meier probability estimates of overall survival across *TP53* subgroups of wild-type *TP53* (WT), single *TP53* mutation (1mut), multiple *TP53* mutations (>1mut), *TP53* mutation(s) and deletion (mut+del), *TP53* mutation(s) and copy-neutral loss of heterozygosity (mut+cnloh). **b**. Cumulative incidence of AML transformation (AMLt) across *TP53* subgroups.

**Extended Data Figure 6.**
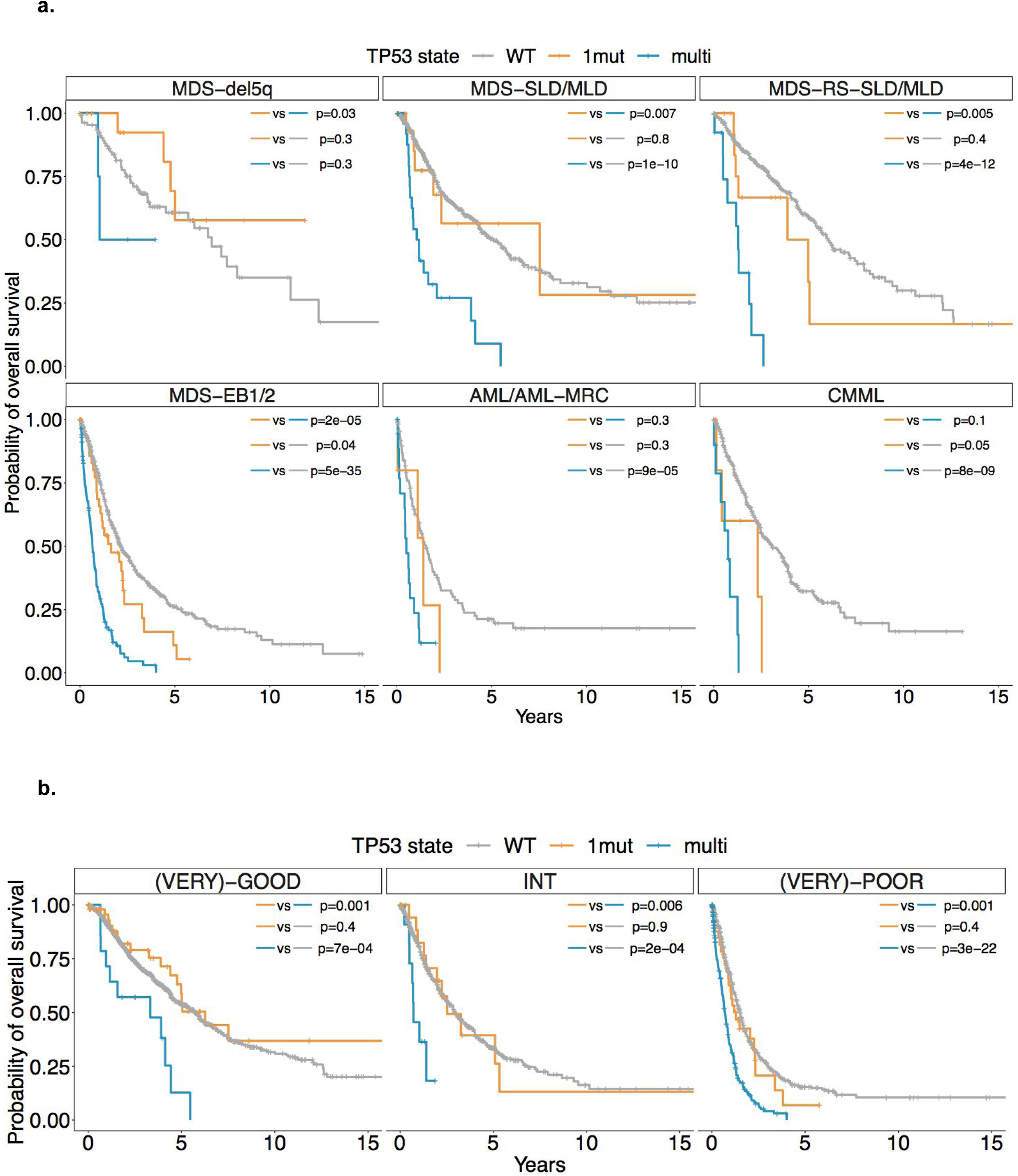
*TP53* allelic state segregates patient outcomes across WHO subtypes and IPSS-R risk groups. **a**. Kaplan-Meier probability estimates of overall survival across main WHO subtypes per *TP53* state of wild-type *TP53* (WT), single *TP53* mutation (1mut) or multiple *TP53* hits (multi). WHO subtypes MDS-SLD and MDS-MLD are merged together as MDS-SLD/MLD and WHO subtypes MDS-EB1 and MDS-EB2 are merged together as MDS-EB1/EB2. **b**. Kaplan-Meier probability estimates of overall survival across IPSS-R risk groups per *TP53* state of wild-type *TP53* (WT), single *TP53* mutation (1mut) and multiple *TP53* hits (multi). IPSS-R very-good and good risk groups are merged together (leftmost panel), and IPSS-R very-poor and poor risk groups are merged together as well (rightmost panel). Annotated p-values are from the log-rank test.

**Extended Data Figure 7.**
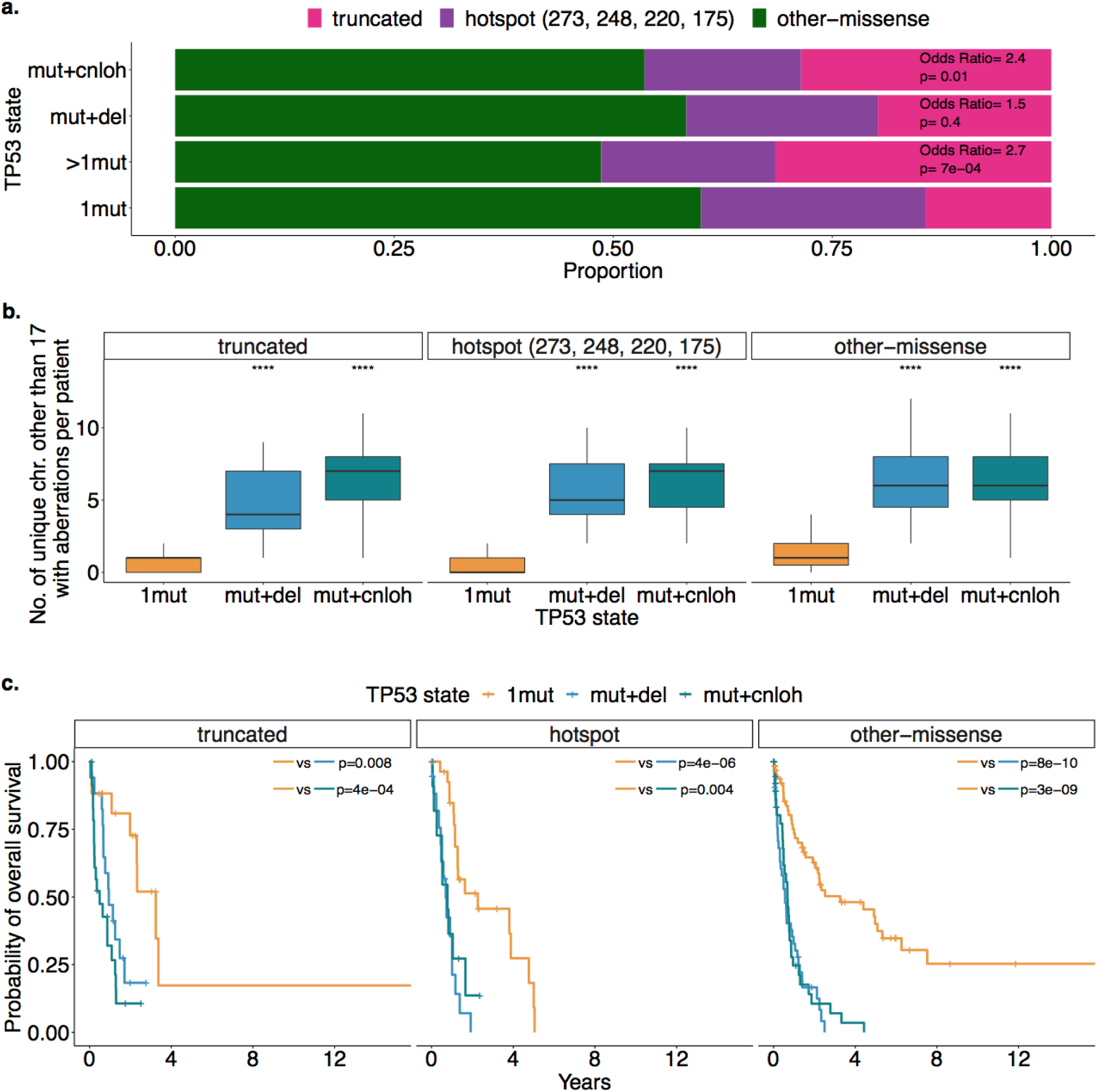
Maintained differences in genome instability levels and outcomes per *TP53* state across mutation types. **a**. Proportion of mutation types across *TP53* subgroups. Truncated mutations (pink) include frameshift indels, nonsense or nonstop mutations and splice-site variants. Mutations annotated as hotspot (purple) are missense mutations at amino acid positions 273, 248, 220 and 175. Mutations annotated as other-missense (green) are additional missense mutations or inframe indels. Odds ratio and Fisher’s test p-values for the proportion of truncated versus non-truncated mutations between the multi-hit *TP53* subgroups and the mono-allelic *TP53* subgroup (1mut) are indicated in the pink parts of the barplot. **b**. Distribution of the number per patient of unique chromosomes other than 17 with aberrations per *TP53* subgroup of single gene mutation (1mut), mutation and deletion (mut+del) and mutation and copy-neutral loss of heterozygosity (mut+cnloh) and across mutation types. Note that 5 patients with both several mutations and deletion or cnloh with ambiguity between the mutation type categories have been excluded for this analysis. ****p<0.0001, Wilcoxon rank-sum test, each compared to the same aberration within the 1mut group. **c**. Kaplan-Meier probability estimates of overall survival per *TP53* subgroup across mutation type. Annotated p-values are from the log-rank test.

**Extended Data Table 1.**
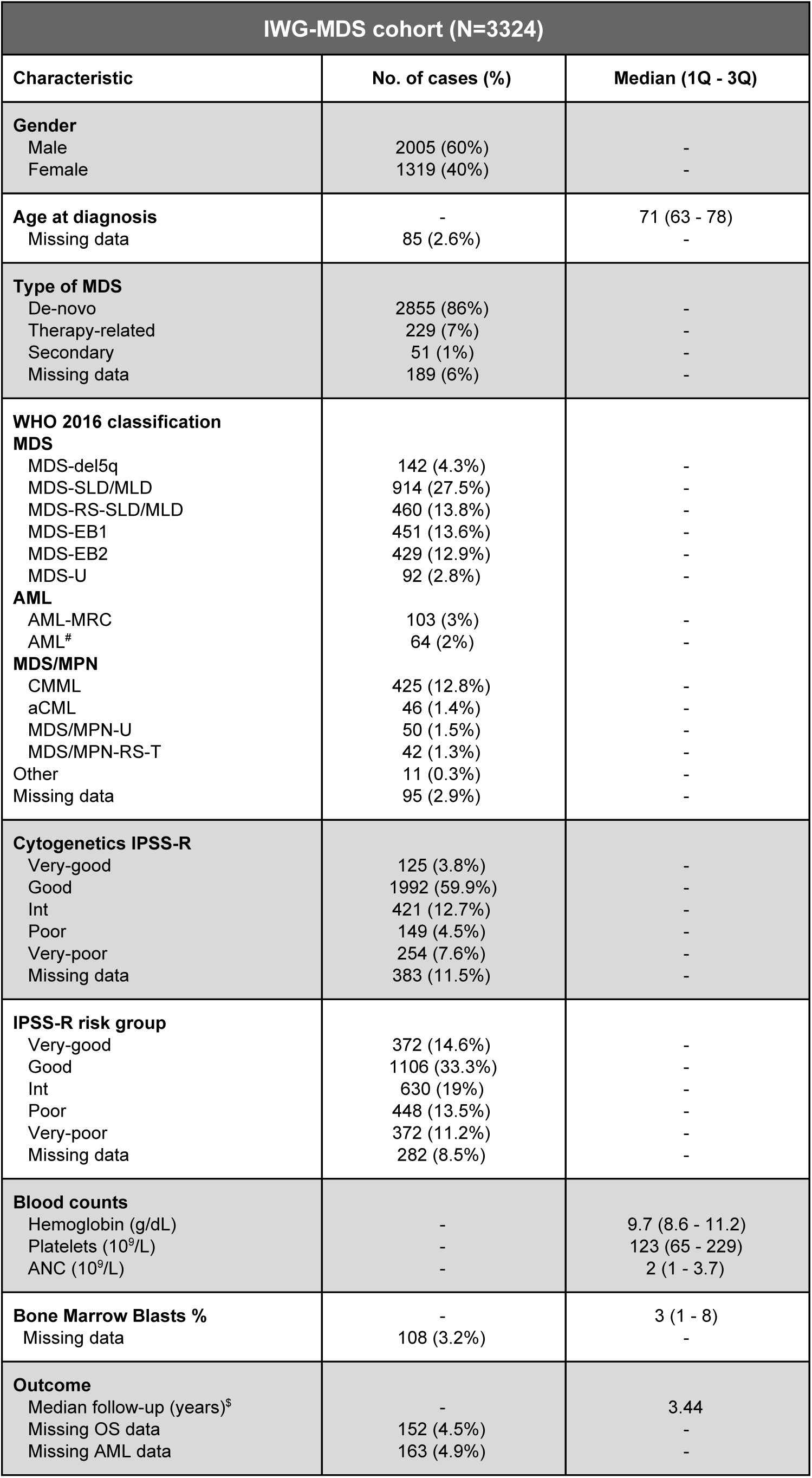
Study cohort characteristics. Table describing the baseline characteristics of the study cohort. 1Q: first quartile; 3Q: third quartile; #: AML classification per WHO 2016 and previously RAEB-T cases. $: Median follow-up time is calculated for censored patients.

**Extended Data Table 2.** Validation cohort characteristics. Table describing the baseline characteristics of the validation cohort. 1Q: first quartile; 3Q: third quartile; $: Median follow-up time is calculated for censored patients.

| Validation cohort (N=1120) |  |  |
| --- | --- | --- |
| Characteristic | No. of cases (%) | Median (1Q - 3Q) |
| **Cohort** |  |  |
| Clinical sequencing | 627 (56%) | - |
| JMPD | 314 (28%) | - |
| JALSG MDS212 | 179 (16%) | - |
| **Gender** |  |  |
| Male | 751 (67%) | - |
| Female | 369 (33%) | - |
| **Age at diagnosis** | - | 65 (54 - 75) |
| Missing data | 121 (11%) | - |
| **WHO 2016 classification** |  |  |
| **MDS** |  |  |
| t-MDS | 9 (0.9%) | - |
| MDS-del5q | 7 (0.6%) | - |
| MDS-SLD | 169 (15.1%) | - |
| MDS-MLD | 100 (8.9%) | - |
| MDS-RS-SLD/MLD | 34 (3%) | - |
| MDS-EB1/2 | 437 (39%) | - |
| MDS-U | 15 (1.3%) | - |
| **AML** |  |  |
| AML-MRC | 121 (10.8%) | - |
| **MDS/MPN** |  |  |
| CMML | 43 (3.8%) | - |
| aCML | 4 (0.4%) | - |
| MDS/MPN-U | 6 (0.5%) | - |
| MDS/MPN-RS-T | 4 (0.4%) | - |
| Missing data | 171 (15.3%) | - |
| **IPSS-R risk group** |  |  |
| Very-good | 22 (4.2%) | - |
| Good | 60 (11.4%) | - |
| Int | 77 (14.6%) | - |
| Poor | 101 (19.1%) | - |
| Very-poor | 166 (31.4%) | - |
| Missing data | 102 (19.3%) | - |
| **Blood counts** |  |  |
| Hemoglobin (g/dL) | - | 8.4 (7.4 - 10.0) |
| Platelets (10<sup>9</sup>/L) | - | 76 (39 - 138) |
| ANC (10<sup>9</sup>/L) | - | 1.2 (0.5 - 2.4) |
| **Bone Marrow Blasts %** | - | 6.8 (2 - 15) |
| Missing data | 554 (49%) | - |
| **Outcome** |  |  |
| Median follow-up (years)<sup>s</sup> | - | 1.1 |
| Missing OS data | 241 (22%) | - |

**Extended Data Table 3.** Characteristics of treated cohort subsets. Table describing the baseline characteristics of the subset of patients that i) received hypomethylating agent (HMA), ii) received Lenalidomide in the context of del(5q) or iii) underwent hematopoietic stem cell transplantation (HSCT).

| Treated cohort subsets |  |  |  |
| --- | --- | --- | --- |
|  | HMA cohort<br>(N=656) | Lenalidomide cohort<br>(N=101) | HSCT cohort<br>(N=310) |
| Characteristic | No. of cases (%) |  |  |
| **TP53 allelic state** |  |  |  |
| Wild type | 511 (78%) | 72 (73%) | 274 (88%) |
| Mono-allelic | 24 (4%) | 12 (12%) | 7 (2%) |
| Multi-hit | 121 (18%) | 17 (15%) | 29 (9%) |
| **TP53 allelic state,**<br>**with outcome data** |  |  |  |
| Wild type | 497 | 69 | 265 |
| Mono-allelic | 22 | 10 | 7 |
| Multi-hit | 119 | 16 | 24 |
| **Gender** |  |  |  |
| Male | 428 (65%) | 35 (35%) | 188 (61%) |
| Female | 228 (35%) | 66 (65%) | 122 (39%) |
| **WHO 2016 classification** |  |  |  |
| MDS-del5q | 4 (0.6%) | 50 (50%) | 5 (1.6%) |
| MDS-SLD/MLD | 84 (13%) | 3 (3%) | 59 (19%) |
| MDS-RS-SLD/MLD | 31 (5%) | 3 (2%) | 21 (6.6%) |
| MDS-EB1/2 | 351 (54%) | 28 (28%) | 144 (46%) |
| MDS-U | 6 (1%) | 4 (4%) | 7 (2%) |
| AML/AML-MRC | 63 (10%) | 6 (7%) | 26 (9%) |
| MDS/MPN | 113 (17%) | 6 (6%) | 45 (14%) |
| Missing data | 4 (0.6%) | 1 (0.9%) | 3 (0.9%) |
| **Cytogenetics IPSS-R** |  |  |  |
| Very-good | 11 (2%) | - | 3 (0.9%) |
| Good | 340 (51%) | 63 (63%) | 177 (56%) |
| Int | 100 (16%) | 6 (6%) | 46 (15%) |
| Poor | 58 (9%) | 3 (3%) | 35 (11%) |
| Very-poor | 111 (17%) | 18 (18%) | 25 (9%) |
| Missing data | 36 (5%) | 11 (11%) | 24 (8%) |
| **IPSS-R risk group** |  |  |  |
| Very-good | 19 (3%) | 3 (3%) | 11 (3%) |
| Good | 95 (14%) | 44 (44%) | 60 (19%) |
| Int | 151 (23%) | 25 (25%) | 89 (28%) |
| Poor | 199 (30%) | 7 (7%) | 87 (28%) |
| Very-poor | 163 (25%) | 17 (17%) | 50 (17%) |
| Missing data | 29 (4%) | 5 (5%) | 13 (4%) |

