## Supplementary for "Implications of *TP53* Allelic State for Genome Stability, Clinical Presentation and Outcomes in Myelodysplastic Syndromes"

#### Table of Contents

**Supplementary Tables** **2**

Supplementary Table 1: International Working Group for Prognosis in MDS 3
Supplementary Table 2: Summary of number of patients with chromosomal aberrations 4
Supplementary Table 3: Assessment of the status of chromosome 17 at the TP53 locus 5

**Supplementary Figures** **6**

Supplementary Figure 1: Sample quality control 7
Supplementary Figure 2: Ascertainment of allelic imbalances by NGS 8
Supplementary Figure 3: Comparison of CBA and NGS 10
Supplementary Figure 4: Comparison of NGS and SNP Array 11
Supplementary Figure 5: Distribution of TP53 mutations 13
Supplementary Figure 6: TP53 molecular landscape 14
Supplementary Figure 7: TP53 mutation distribution per TP53 subgroup 15
Supplementary Figure 8: Frequency distribution of chromosomal aberrations per TP53 state 16
Supplementary Figure 9: TP53 clinical correlates 17
Supplementary Figure 10: Complex karyotype and TP53 mutations 18
Supplementary Figure 11: Overall survival per del(5q) status and TP53 state 19
Supplementary Figure 12: Multivariate Cox models with TP53 state alongside IPSS-R risk group 20
Supplementary Figure 13: Complex karyotype and TP53 state 21
Supplementary Figure 14: Overall survival per TP53 state and VAF stratification 23
Supplementary Figure 15: Overall survival per hotspot status and TP53 state 24
Supplementary Figure 16: Genome stability across de-novo or therapy-related MDS and TP53 state 26
Supplementary Figure 17: Clonal evolution from MDS to AML of TP53 mutated patients 27

a. Patient 128470 28
b. Patient 129056 30
c. Patient 129363 31
d. Patient 129525 32
e. Patient 129602 33
f. Patient 129662 34
g. Patient 129763 35
h. Patient 128715 36
i. Patient 128829 37
j. Patient 129119 38
k. Patient 129047 39
l. Patient 130333 40

Supplementary Figure 18: Representation of TP53 subgroups and states in the validation cohort 42
Supplementary Figure 19: Implications of TP53 state to genome stability in the validation cohort 43
Supplementary Figure 20: Clinical correlates of TP53 state in the validation cohort 45

**Supplementary References** **47**

### Supplementary Tables

### Supplementary Table 1: International Working Group for Prognosis in MDS

Table listing the collaborative centers of the International Working Group for Prognosis in MDS (IWG-MDS) that provided samples and contributed to the final study cohort of 3,324 peri-diagnostic treatment naive MDS samples. Quality control processing steps are described in the Methods and Supplementary Figure 1.

| **Collaborating Center** | **Center Code** | **No. of samples sequenced (excluding serial data) N=4,105** | **No. of samples post-QC that are included in the study cohort N=3,324** |
|---|---|---|---|
| KAROLINSKA INSTITUTE | KI | 1051 | 898 |
| DUSSELDORF MDS REGISTRY | DUS | 534 | 459 |
| UNIVERSITY OF PAVIA | PV | 341 | 316 |
| LA FE UNIVERSITY HOSPITAL | GESMD | 256 | 246 |
| RADBOUDUMC MEDICAL CENTER NIJMEGEN<br>AMSTERDAM UMC/VU UNIVERSITY MEDICAL CENTER | RMCN | 209 | 199 |
| COCHIN HOSPITAL | CCH | 166 | 159 |
| CHANG GUNG MEMORIAL HOSPITAL | CGM | 109 | 107 |
| GRUPPO ROMANO LAZIALE MDS | ROM | 116 | 104 |
| UNIVERSITY OF BOLOGNA | UOB | 97 | 88 |
| MEDICAL UNIVERSITY OF VIENNA | MUV | 83 | 83 |
| HANNOVER MEDICAL SCHOOL | HMS | 244 | 83 |
| UNIVERSITY HOSPITAL DRESDEN | TUD | 77 | 73 |
| FEDERAL UNIVERSITY OF CEARA | FUCE | 83 | 73 |
| INSTITUT JOSEP CARRERAS | ICO | 72 | 71 |
| AOU CAREGGI HOSPITAL | FLO | 78 | 68 |
| DEMOCRITUS UNIVERSITY OF THRACE | DUTH | 67 | 66 |
| UNIVERSITY OF OXFORD | UOXF | 58 | 50 |
| HOSPITAL ISRAELITA ALBERT EINSTEIN | HIAE | 49 | 47 |
| MEMORIAL SLOAN KETTERING CANCER CENTER | MSK | 38 | 36 |
| VANDERBILT UNIVERSITY | VU | 36 | 33 |
| INSTITUTE OF HEMATOLOGY AND BLOOD TRANSFUSION | IHBT | 36 | 33 |
| UNIVERSITY MEDICINE GOTTINGEN | UMG | 26 | 26 |
| RETE EMATOLOGICA LOMBARDA | REL | 228 | 6 |
| SAINT LOUIS HOSPITAL | SLS | 51 | - |

#### Supplementary Table 2: Summary of number of patients with chromosomal aberrations

Summary table of number of patients with different types of chromosomal aberrations, also broken down for *TP53* wild-type or *TP53*-mutated patients. cnLOH: copy neutral loss-of-heterozygosity. $: complex karyotype is defined as 3 independent chromosomal abnormalities, as explained in (Schanz et al. 2012). #: monosomal karyotype is defined as 2 or more autosomal monosomies or 1 autosomal monosomy and structural abnormalities, as suggested in (Breems et al. 2008).

| | **Overall cohort N=3324** | ***TP53* wild type N=2946** | ***TP53* mutated N=378** | **Odds ratio (95 CI)** | **Fisher exact test p-value** |
|---|---|---|---|---|---|
| Number (%) of cases with any chromosomal aberration (including cnLOH) | 1571 (47%) | 1247 (42%) | 327 (86%) | 8 (6-12) | $<10^{-16}$ |
| Number (%) of cases with any deletion | 1075 (32%) | 765 (26%) | 310 (82%) | 13 (10-17) | $<10^{-16}$ |
| Number (%) of cases with any deletion on different chr. than 17 | 1064 (32%) | 754 (26%) | 310 (82%) | 13 (10-17) | $<10^{-16}$ |
| Number (%) of cases with any gain | 575 (18%) | 361 (12%) | 214 (56%) | 10 (8-12) | $<10^{-16}$ |
| Number (%) of cases with any rearrangement | 282 (8%) | 162 (6%) | 120 (32%) | 8 (6-10) | $<10^{-16}$ |
| Number (%) of cases with any cnLOH | 360 (11%) | 267 (9%) | 93 (24%) | 3 (2-4) | $<10^{-15}$ |
| Number (%) of cases with any cnLOH on different chr. than 17 | 278 (8%) | 261 (9%) | 17 (4%) | 0.5 (0.3-0.8) | 0.003 |
| Number (%) of cases with more than 3 chr. affected with aberrations (including cnLOH) | 378 (11%) | 128 (4%) | 250 (66%) | 42 (32-56) | $<10^{-16}$ |
| Number (%) of cases classified as complex karyotype$^{\$}$ | 329 (10%) | 82 (3%) | 247 (65%) | 63 (46-87) | $<10^{-16}$ |
| Number (%) of cases classified as monosomal karyotype$^{\#}$ | 177 (5%) | 22 (0.7%) | 155 (41%) | 91 (57-153) | $<10^{-16}$ |

### Supplementary Table 3: Assessment of the status of chromosome 17 at the *TP53* locus

Table describing the status of chromosome 17 at the *TP53* locus on the study cohort of 3,324 MDS patients. cnLOH: copy-neutral loss-of-heterozygosity, i(17q): isochromosome 17q, CBA: conventional G-banding analysis, NGS: next generation sequencing, NA: not applicable.

| **Status of chromosome 17 at *TP53* locus** | **No. of cases** | **No. of cases with 1 *TP53* mutation** | **No. of cases with more than 1 *TP53* mutation** | **No. of cases with evidence from CBA and NGS** | **No. of cases with evidence from CBA only** | **No. of cases with evidence from NGS only** |
|---|---|---|---|---|---|---|
| cnLOH | 80 | 73 | 5 | NA | NA | 80 |
| Deletion | | | | | | |
| -17 / del17p | 97 | 76 | 9 | 60 | 11 | 26 (18 focal deletions) |
| i(17q) | 10 | 0 | 0 | 7 | 1 | 2 |
| Gain | 1 | 1 | 0 | 1 | 0 | 0 |

### Supplementary Figures

### Supplementary Figure 1: Sample quality control

Schematic of the sample quality control (QC) workflow to build the IWG-PM cohort.

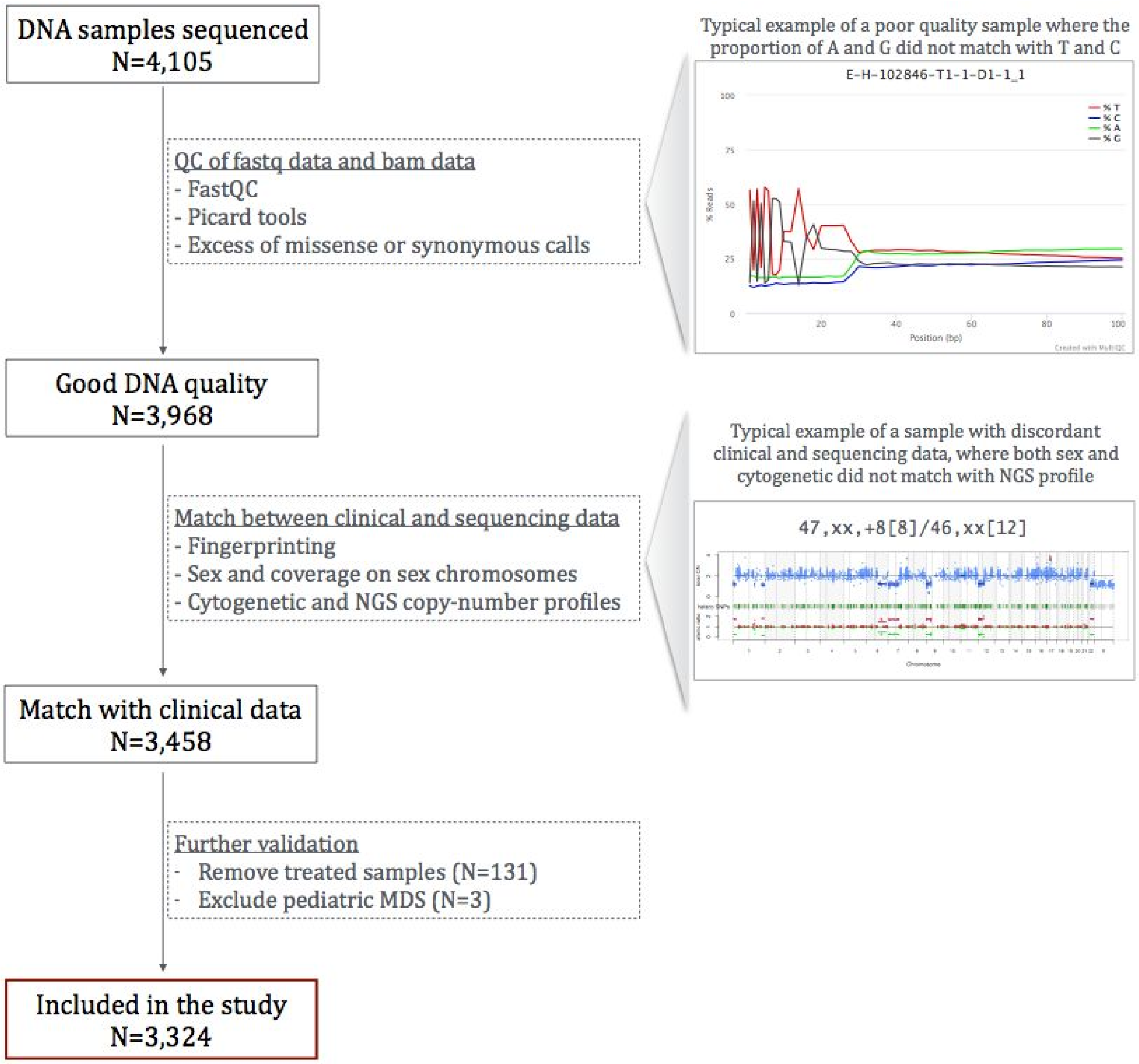

#### Supplementary Figure 2: Ascertainment of allelic imbalances by NGS

Examples of total copy number (CN) and allele specific CN states in representative cases of the study cohort, derived with CNACS algorithm (Yoshizato et al. 2017). The x-axis corresponds to genome wide chromosomal coordinates. On the upper panel, y-axis represents total CN, whereas on the lower panel y-axis represents allele specific CNs. Blue dots show total CN on each coverage probe, red and green dots show major and minor CNs, respectively, on each heterozygous SNP probe. Aberrant segments called by CNACS are underlined with solid lines. Conventional G-banding analysis (CBA) data are also provided for comparison. **a**. Example of a case with arm and chromosome level deletions detected by both CNACS and CBA. **b**. Representative case with concordant findings between CNACS and CBA for -7 and -18, and further detection with CNACS of a focal deletion on chromosome 17 overlapping the *TP53* locus. **c.** Example of a case with concordant arm and chromosome level findings between CBA and CNACS and a copy-neutral LOH at chromosome 17p.

**a.**

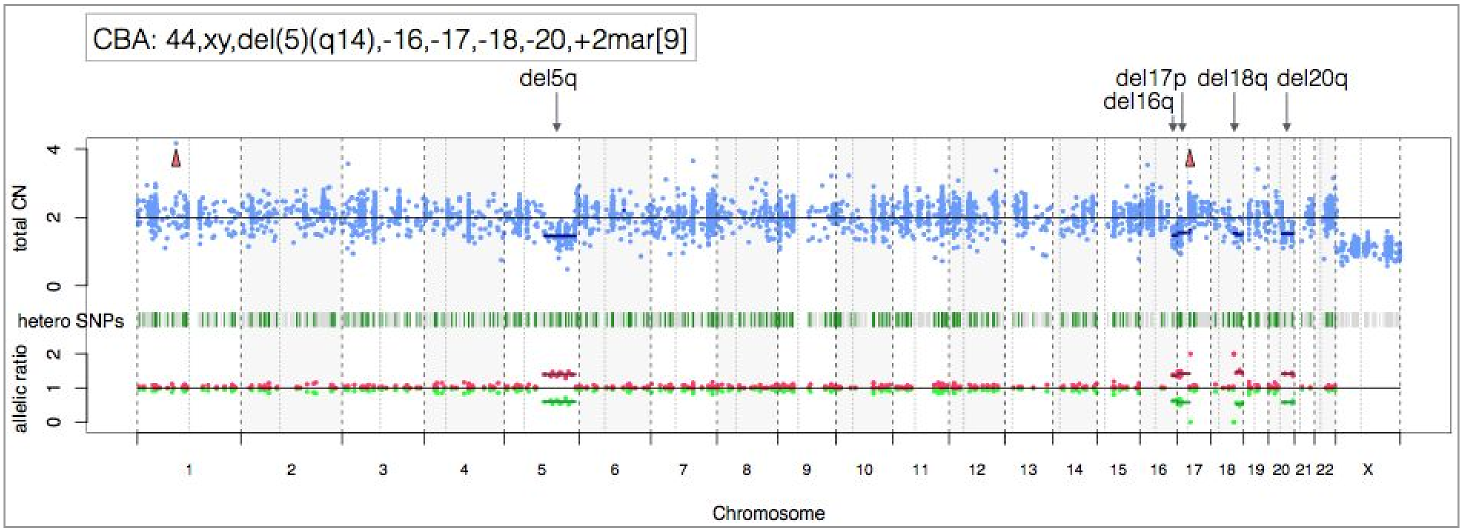


**b.**

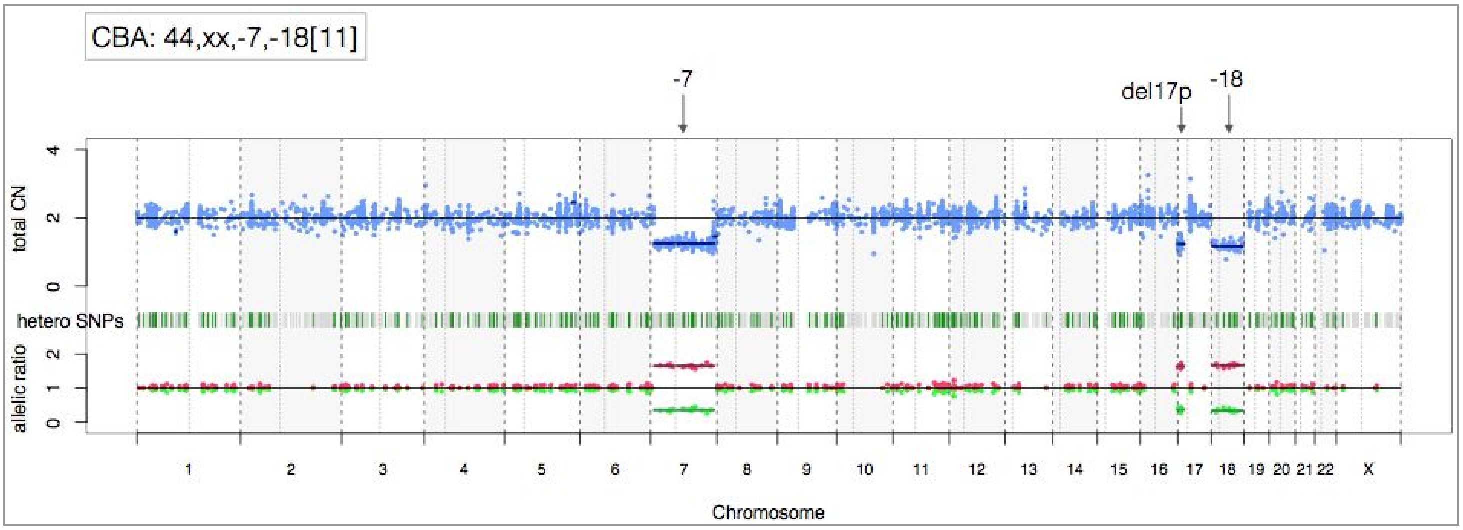

**c.**

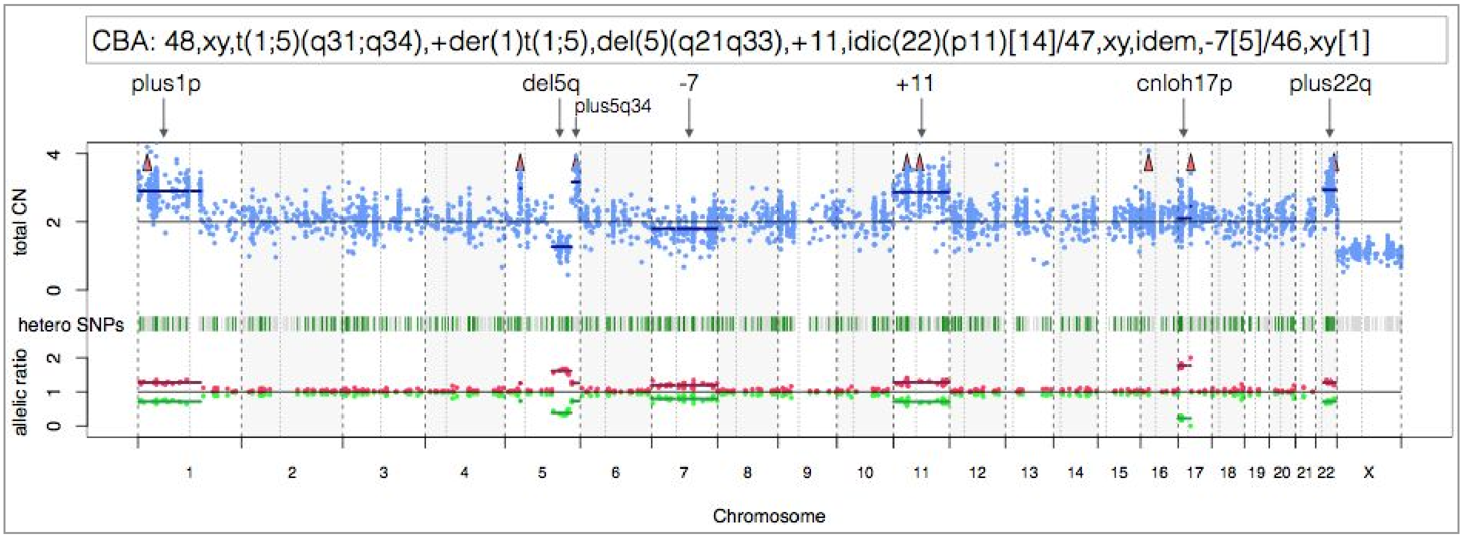
CBA: 48,xy,t(1;5)(q31;q34),+der(1)t(1;5),del(5)(q21q33),+11,idic(22)(p11)[14]/47,xy,idem,-7[5]/46,xy[1]
plus1p
del5q
plus5q34
-7
+11
cnloh17p
plus22q
total CN
hetero SNPs
allelic ratio
Chromosome

### Supplementary Figure 3: Comparison of CBA and NGS

**a.** Comparison of ploidy aberrations on autosomal chromosomes across 2,931 patients with both CBA data and NGS derived copy-number profiles available. Aberrations observed at frequency >0.5% are depicted on the x-axis. Colors represent the combination of assays that detect the aberrations: both CBA and NGS in blue, CBA only in red, NGS only in yellow. **b.** Comparison of Kaplan-Meier (KM) probability estimates of overall survival from patients with del5q (left), plus8 (middle) or del7 (right) detected with CBA only (red), both CBA and NGS (blue) or NGS only (yellow). KM curves follow the same trend within each panel with no statistical differences. **c.** Evaluation of KM probability estimates of overall survival across 3,324 patients annotated as complex karyotype or not. Cases annotated as complex karyotype from CBA data (N=310, 299 with outcome data) are shown in dark green, cases not marked as complex from CBA data but showing evidence of complex from NGS copy-number profiles (N=15, 14 with outcome data) are shown in intermediate green, and cases with missing CBA data but annotated as complex from the NGS copy-number profiles (N=13, 10 with outcome data) are shown in light green. The 3 complex karyotype categories follow the same trend with no statistical differences.

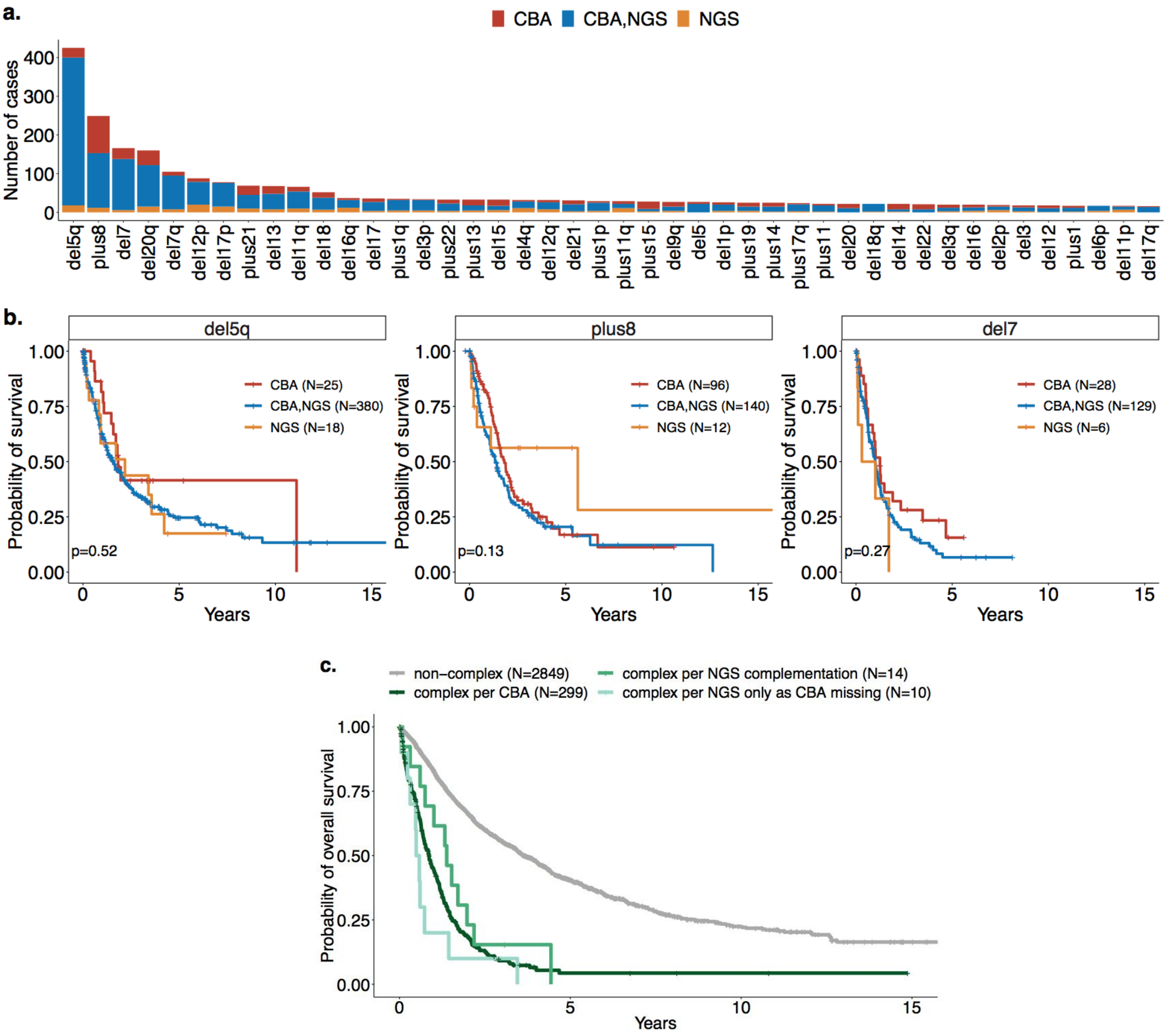

### Supplementary Figure 4: Comparison of NGS and SNP Array

We performed a comparison of NGS derived chromosomal aberrations and SNP array results on 21 samples. SNP arrays were performed with Affymetrix Cytoscan HD array with 2.67 million probes including 780K SNP markers. Samples were selected based on the identification with CNACS of regions with copy-neutral loss of heterozygosity (cnloh) or focal deletions. 4 cases were selected based on the identification of focal deletion of 17p at the *TP53* locus, and were complex karyotype (CK). 5 cases were selected based on the identification of cnloh of 17p at the *TP53* locus and were CK. One of them also had cnloh of 11q at low level. 4 cases were selected based on the identification of cnloh of 11q, 3 others based on cnloh of 4q, and 3 additional cases based on cnloh of 7q. One more case based on both cnloh of 7q and 21q and one last case based on both cnloh of 4q and 7q.

The concordance between findings from NGS profiles and SNP array profiles was extremely concordant. With the exception of one aberration that we annotated as cnloh of 7q whilst SNP array showed a focal deletion, all aberrations between the two assays were concordant across the selected samples.

**a**. Example of a case with focal terminal deletion of 17p at the *TP53* locus along 3 other deletions. Concordant findings are observed between CNACS profile (top) and SNP array profile (bottom). **b.** Representative sample with cnloh of 17p at the *TP53* locus along 4 other deletions and subclonal cnloh of 11q. Concordant findings are observed between CNACS profile (top) and SNP array profile (bottom). **c.** Case with cnloh of 4q along deletion 20q observed in both CNACS profile (top) and SNP array profile (bottom).

**a.**

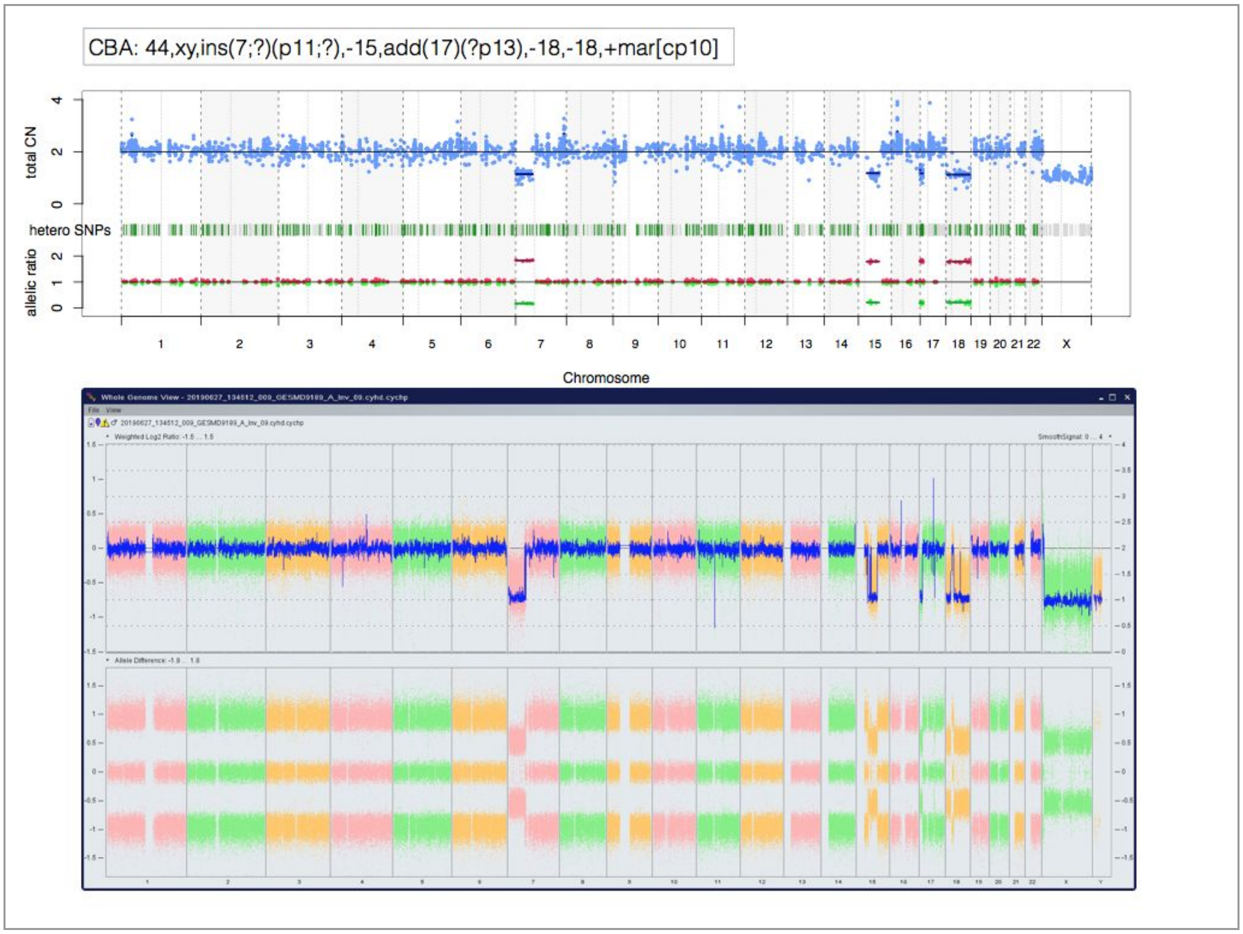

**b.**

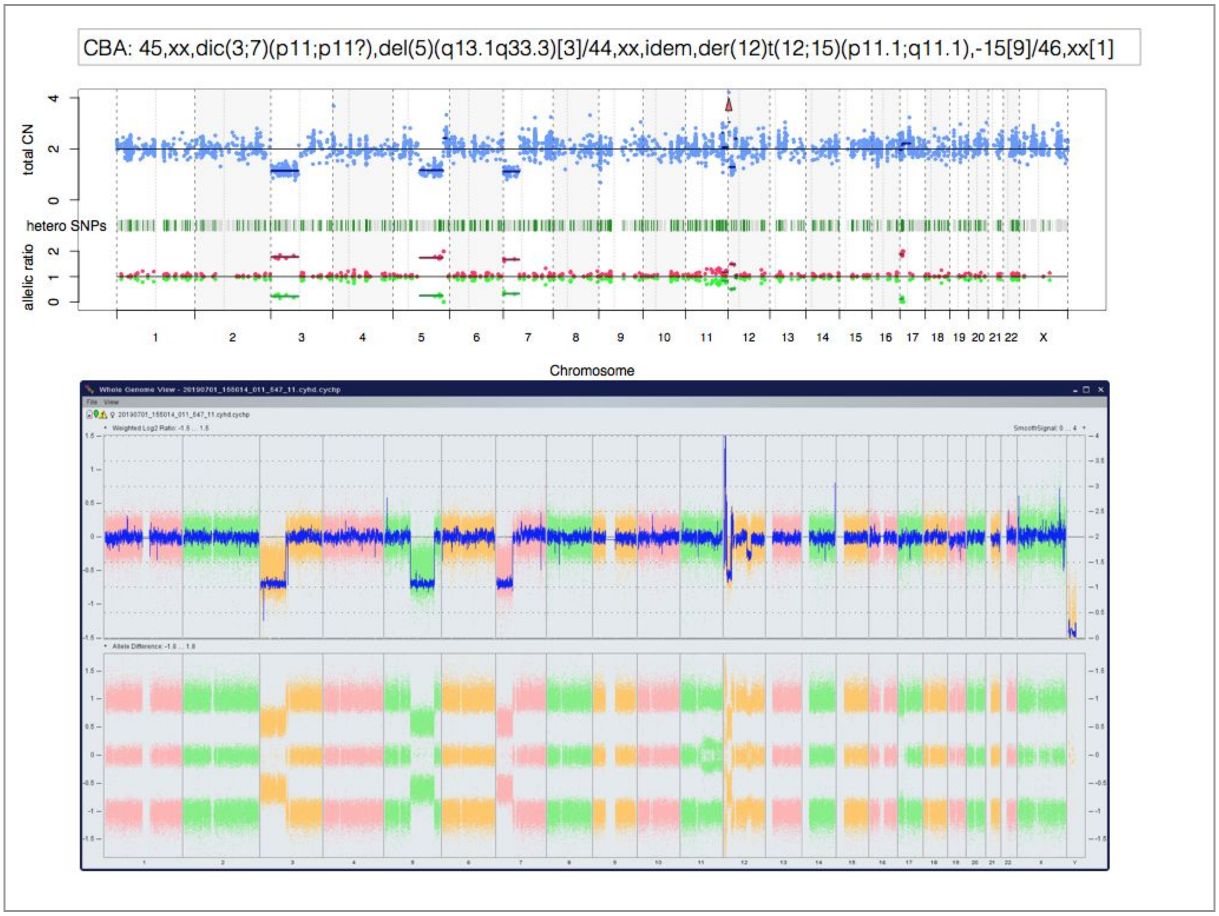
CBA: 45,xx,dic(3;7)(p11;p11?),del(5)(q13.1q33.3)[3]/44,xx,idem,der(12)t(12;15)(p11.1;q11.1),-15[9]/46,xx[1]
total CN
hetero SNPs
allelic ratio
Chromosome

**c.**

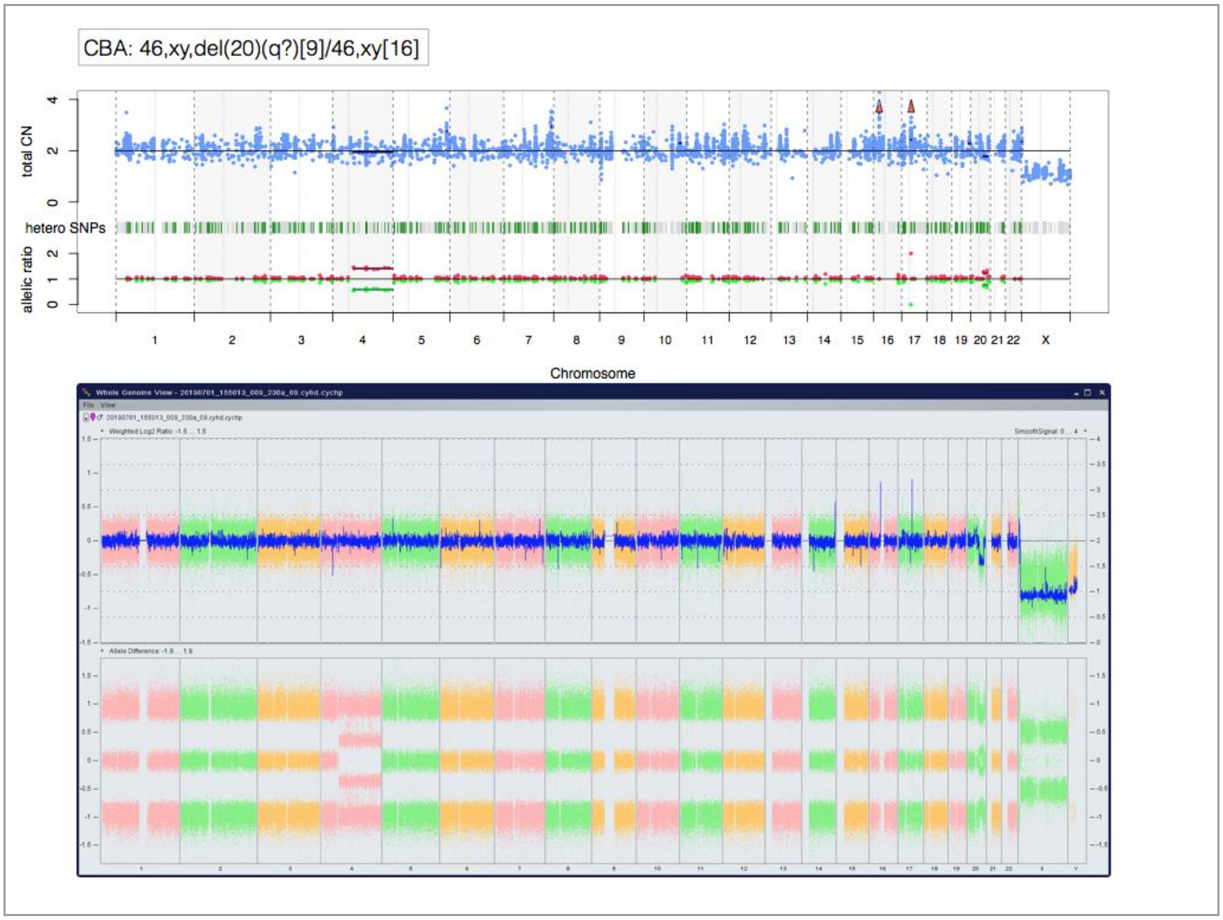
CBA: 46,xy,del(20)(q?)[9]/46,xy[16]
total CN
hetero SNPs
allelic ratio
Chromosome

### Supplementary Figure 5: Distribution of *TP53* mutations

**a.** Distribution of 486 mutations along *TP53* gene body from 378 MDS patients. Mutations annotated as truncated (shown in black) include nonsense, nonstop, frameshift deletion, frameshift insertion and splice site mutations. P53 DNA-binding domain is depicted in red, P53 tetramerization motif in blue. **b.** Distribution of 110 mutations along *TP53* gene body as presented in cBioPortal (Cerami et al. 2012) from all publicly available data in myeloid disease (n=2514 patients, 8 studies in total). cBioPortal: https://cbioportal.mskcc.org.

**a.**

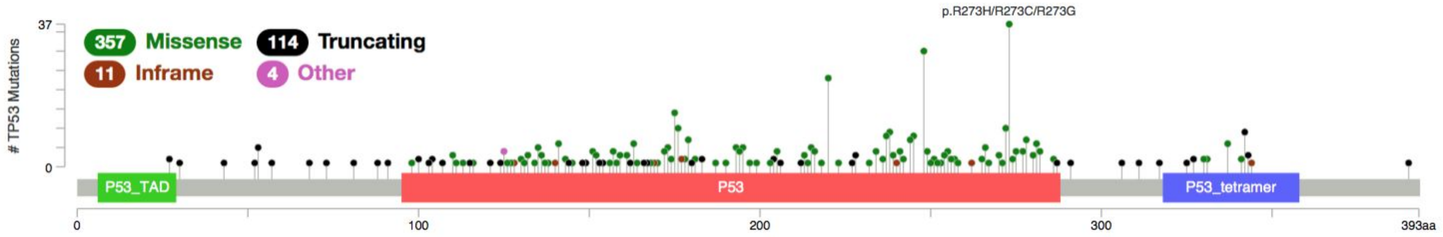


**b.**

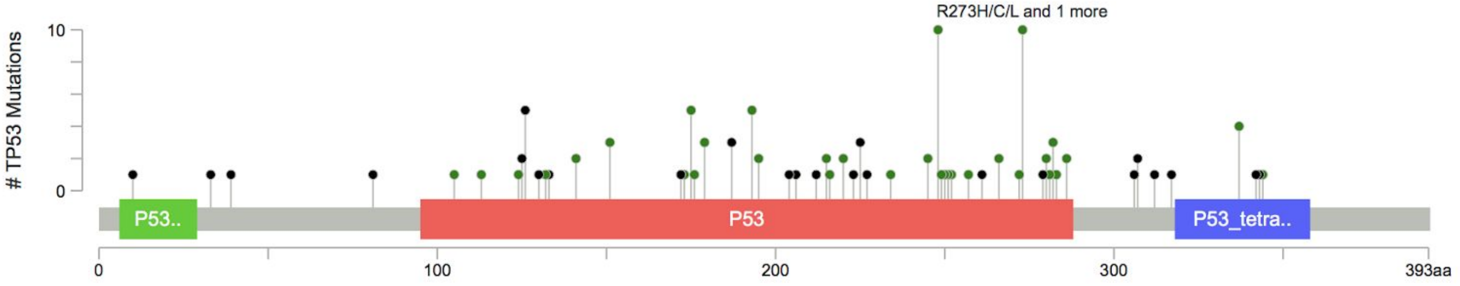

#### Supplementary Figure 6: *TP53* molecular landscape

**a.** Number of patients with 1, 2 or 3 mutations in *TP53*. Within the *TP53* mutated patients, 72% (N=274) have 1 single mutation while 28% (N=104) have multiple mutations. **b.** Density estimation of variant allele frequency (VAF) of 486 *TP53* mutations from 378 patients. The estimated density is trimodal with modes at approximately VAF 10%, 35% and 80%. The high values of VAF distributed around the third 80% mode are indicative of loss-of-heterozygosity. Of note, 97 mutations have a VAF above 55%. **c.** Number of unique *TP53* mutations with observed recurrence across the study cohort. If the recurrence equals 1 the mutation is observed in a single patient, if the recurrence equals, say, 4 the mutation is observed in 4 patients. A mutation is defined here as chromosome, position, reference allele, alternate allele. Colors are indicative of the effect of mutations. Most recurrent protein changes are annotated in text.

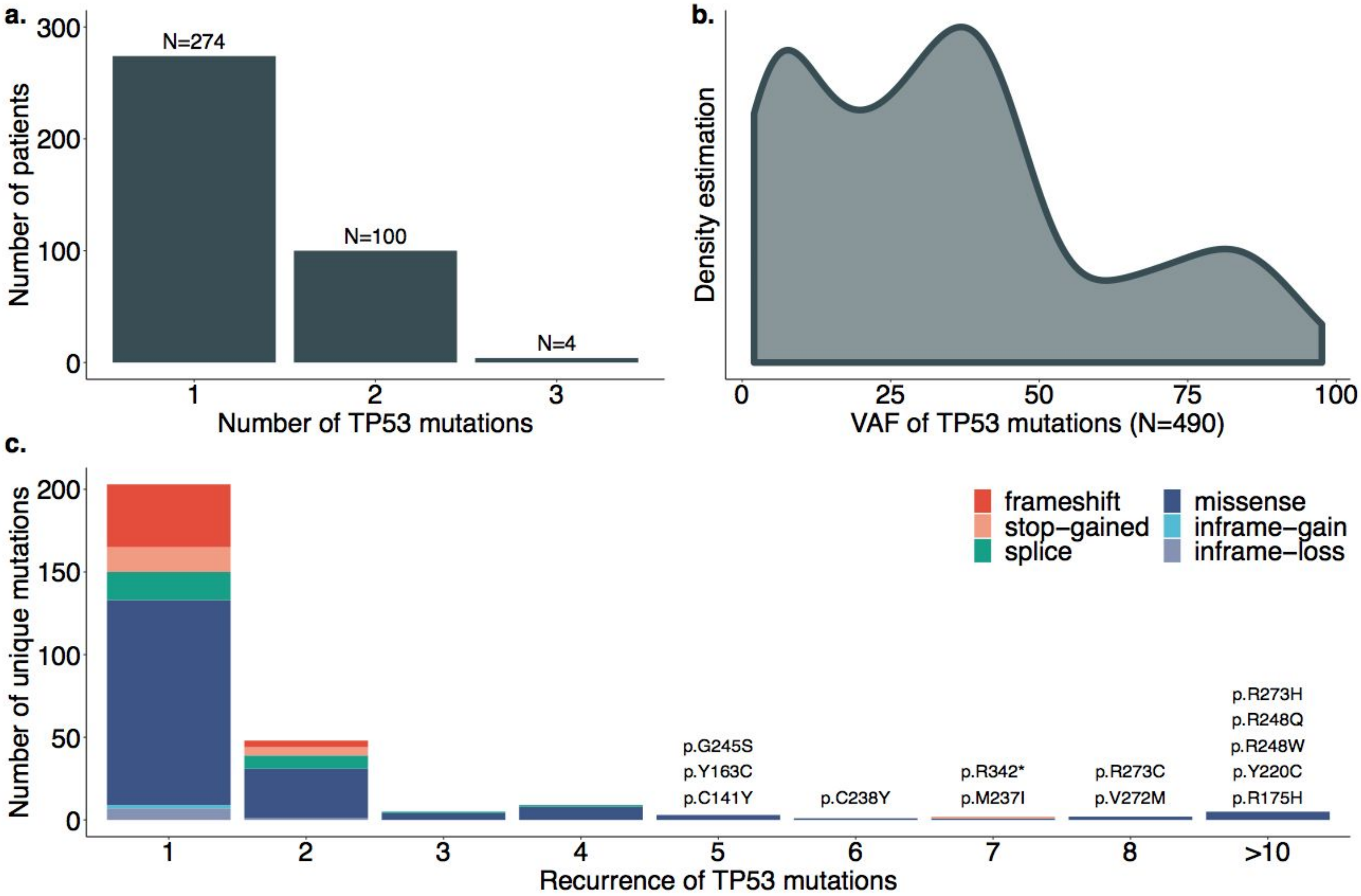

#### Supplementary Figure 7: *TP53* mutation distribution per *TP53* subgroup

Distribution of *TP53* mutations along the gene body per *TP53* subgroup. From top to bottom: *TP53* subgroup of single gene mutation (1mut, N=125 mutations), multiple mutations (>1mut, N=181 mutations), mutation(s) and deletion (mut+del, N=96 mutations), mutation(s) and copy-neutral loss-of-heterozygosity (mut+cnloh, N=84 mutations). Mutations annotated as truncated include nonsense, nonstop, frameshift deletion, frameshift insertion and splice site mutations. P53 DNA-binding domain is depicted in red, P53 tetramerization motif in blue.

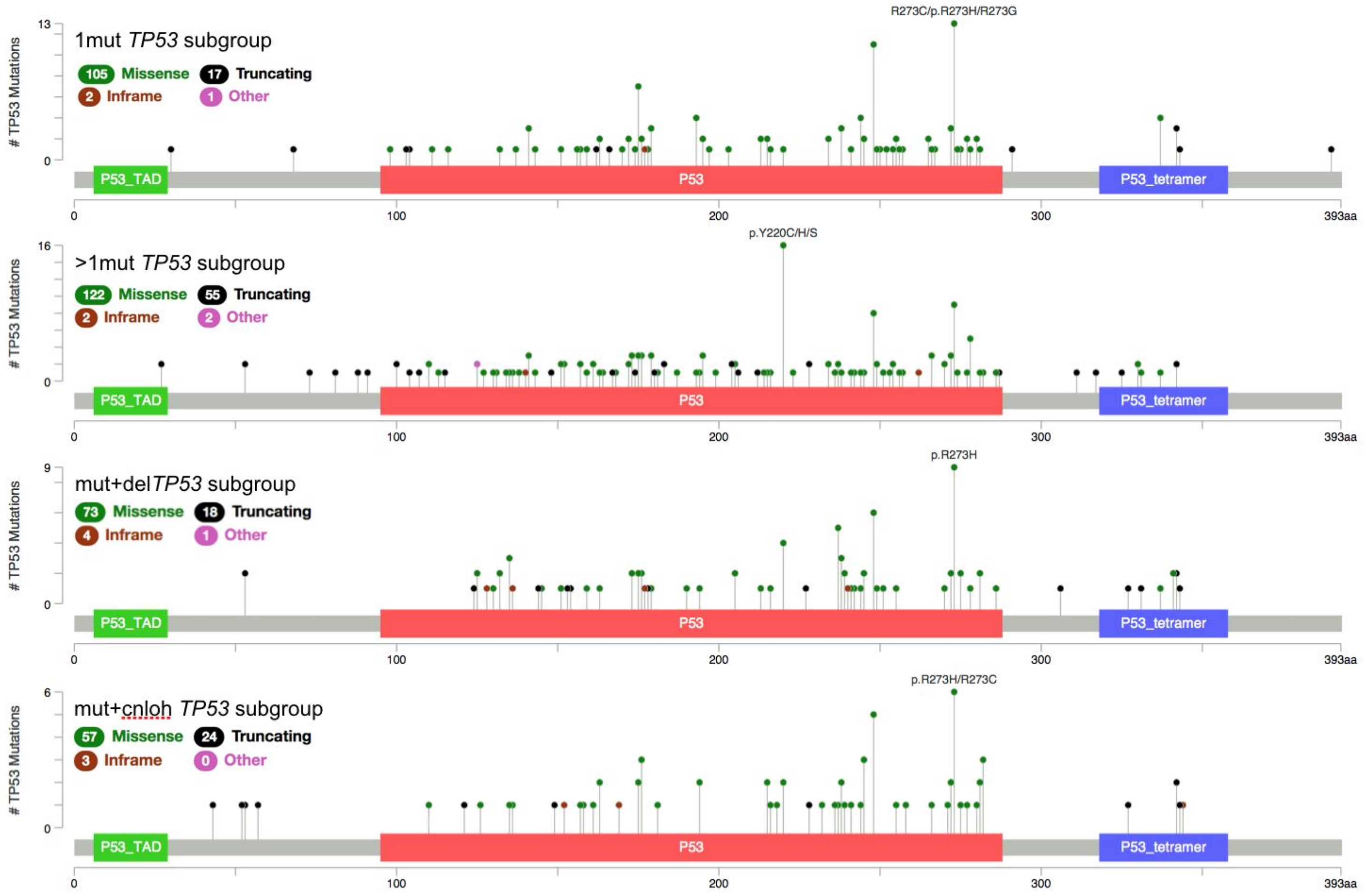

### Supplementary Figure 8: Frequency distribution of chromosomal aberrations per *TP53* state

**a.** Frequency distribution of chromosomal aberrations within the *TP53* state of single gene mutation. The x-axis shows aberrations observed in more than 2% of patients with one *TP53* mutation. **b.** Frequency distribution of chromosomal aberrations within the *TP53* state of multiple hits. The x-axis shows aberrations observed in more than 2% of patients with multiple *TP53* hits. Colors represent the contributions of specific *TP53* subgroups of multiple *TP53* mutations (>1mut), mutation(s) and deletion (mut+del) and mutation(s) and copy-neutral loss-of-heterozygosity (mut+cnloh). Aberrations include deletion (del), gain (plus), copy-neutral loss of heterozygosity (cnloh), presence of marker chromosome (mar), presence of ring chromosome (ring), rearrangement (r_i_j denotes a rearrangement between chromosome i and j) and whole genome amplification (WGA). Of note, the y-axis in panels a and b are on different scales, such that del5q reaches 34% of patients with one single *TP53* mutation while it reaches 85% of patients with multiple *TP53* hits (OR=10, $p<10^{-16}$ Fisher exact test).

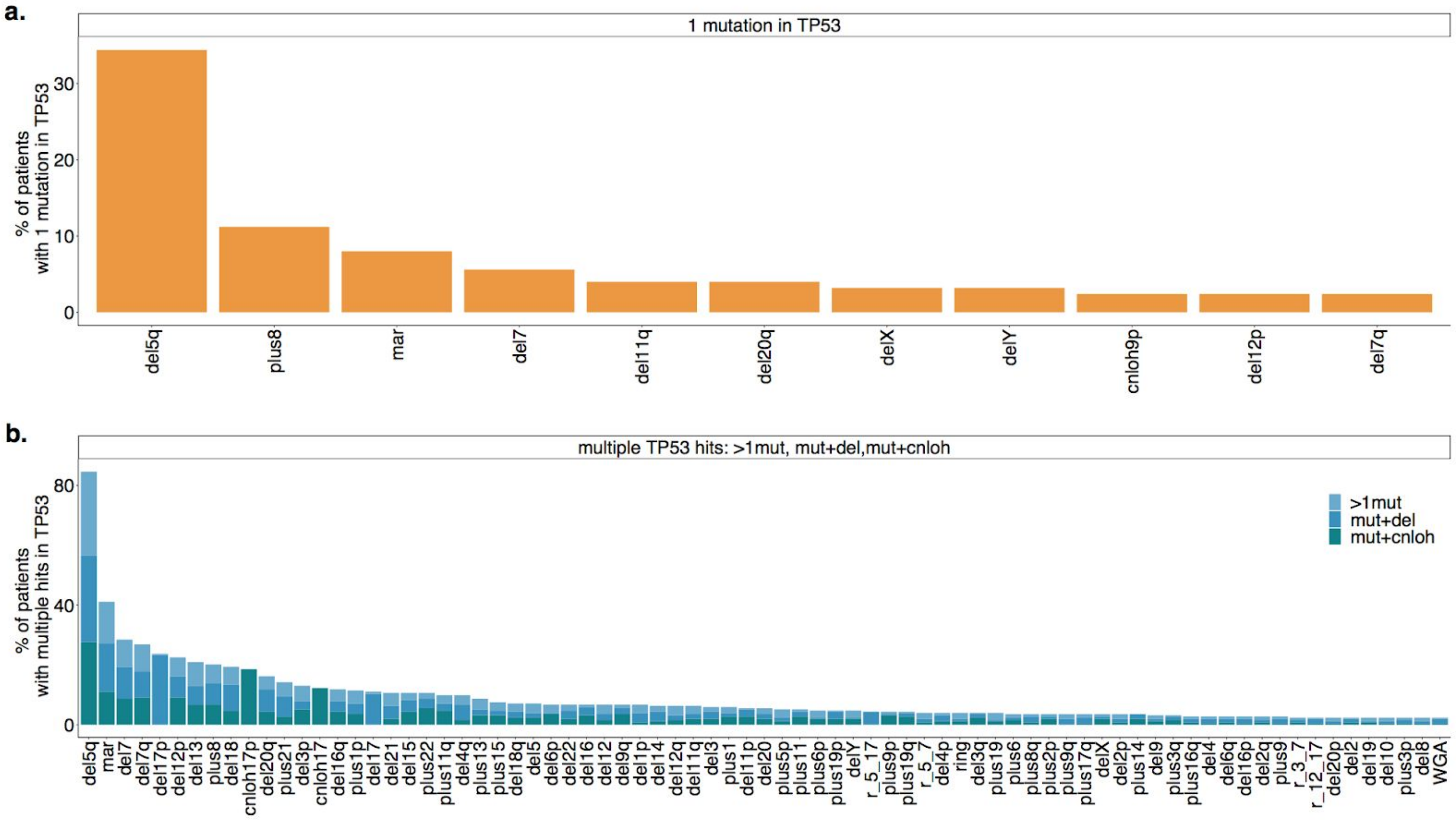

### Supplementary Figure 9: *TP53* clinical correlates

**a-c.** Boxplots indicative of the levels of cytopenias of the *TP53* mutated patients (MUT) compared to *TP53* wild-type patients (WT). Panel **a.** shows hemoglobin level, panel **b.** shows platelets count and panel **c.** shows absolute neutrophil count (ANC). Black lines represent the median level and filled boxes extend from 25% to 75% quantiles. The y-axis are root-squared transformed. **d.** Distribution of bone marrow blasts percentage of the *TP53* mutated (MUT) and wild-type (WT) patients. ****p<0.0001, Wilcoxon rank-sum test. **e-f.** Comparison of outcomes of *TP53* mutated and wild-type patients. Panel e. shows Kaplan-Meier probability estimates of overall survival (HR=2.7, 95%CI: 2.4-3.1, $p<10^{-16}$ Wald test). Panel f. shows cumulative incidence function of AML transformation (AMLt) (HR=2.9, 95%CI: 2.4-3.7, $p<10^{-16}$ Wald test).

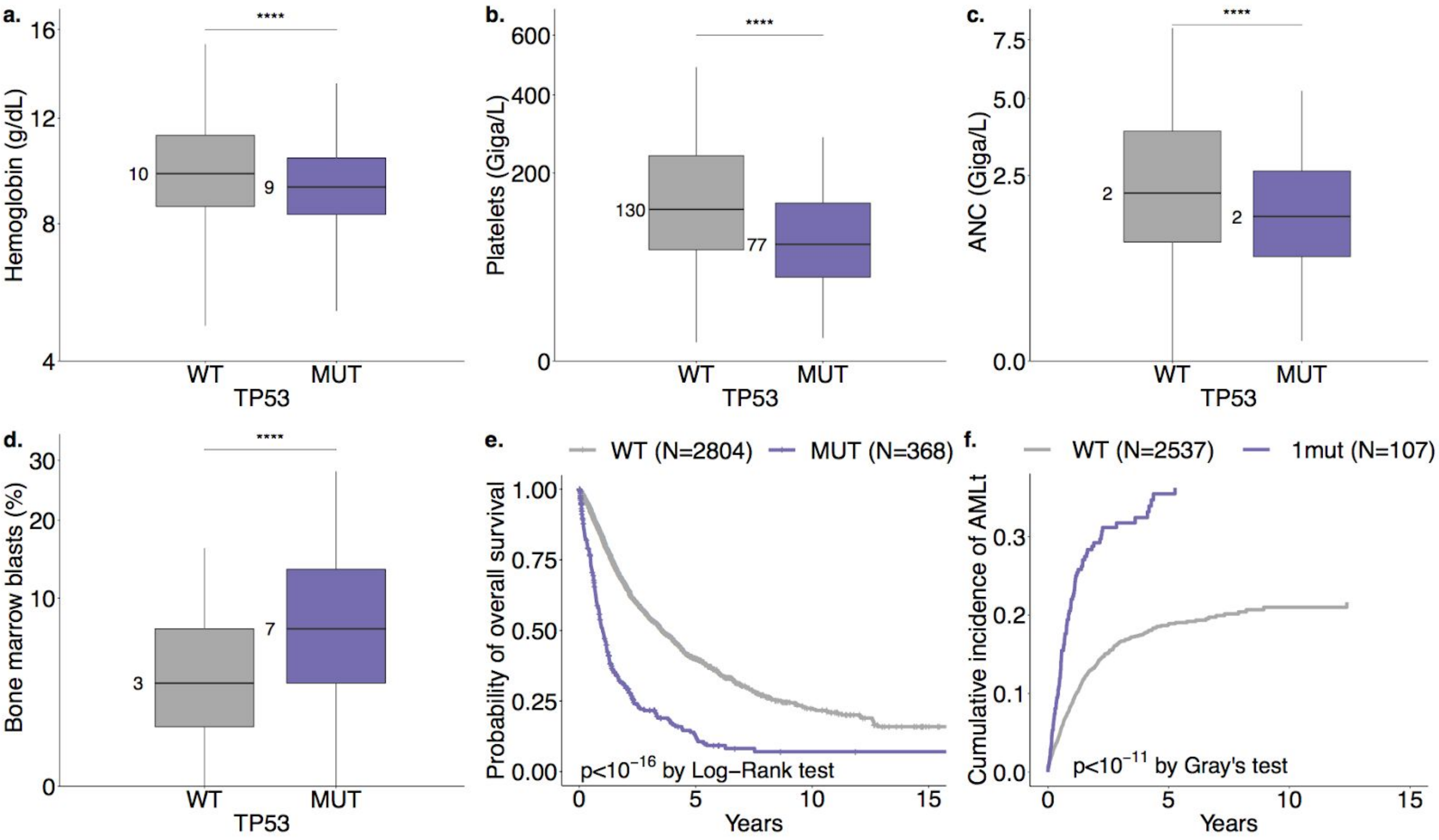

### Supplementary Figure 10: Complex karyotype and *TP53* mutations

Kaplan-Meier probability estimates of overall survival across 329 patients with complex karyotype and with *TP53* wild-type (WT) (N=247, 242 with survival data) or mutated (MUT) (N=82, 81 with survival data). Patients with complex karyotype and mutated *TP53* have significantly worse overall survival than patients with complex karyotype and wild-type *TP53* (HR=2.1, 95%CI: 1.6-2.9, $p<10^{-5}$ Wald test).

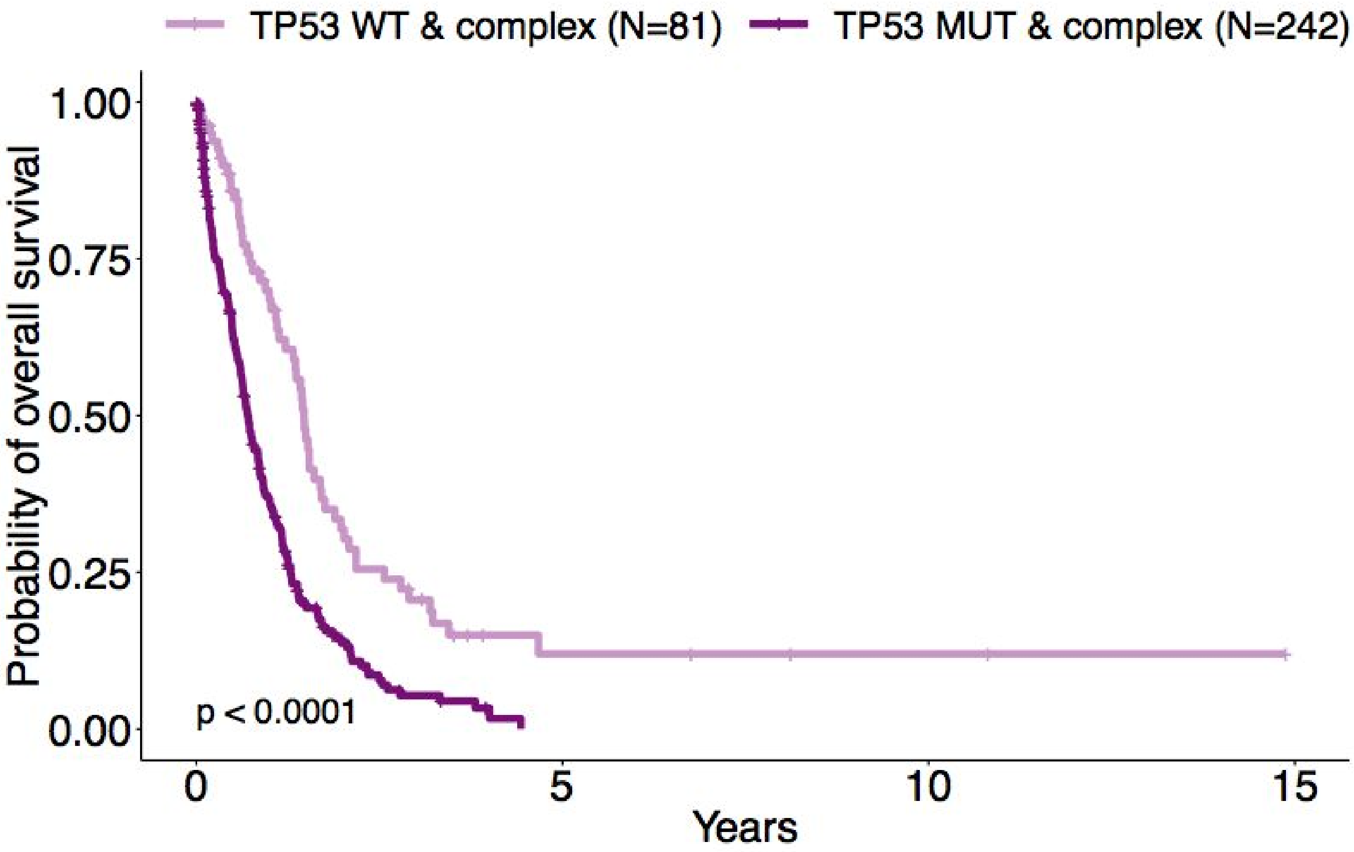

### Supplementary Figure 11: Overall survival per del(5q) status and *TP53* state

Kaplan-Meier probability estimates of overall survival per *TP53* state of wild-type *TP53* (WT), single *TP53* gene mutation (1mut) and multiple *TP53* hits (multi), and across presence (dashed line) or absence (solid line) of deletion 5q (del5q). Among the 2,922 WT cases, 223 have del(5q). Among the 125 1mut cases, 43 have del(5q). Among the 253 multi cases, 214 have del(5q). P-values annotated on the panels are from the log-rank test.

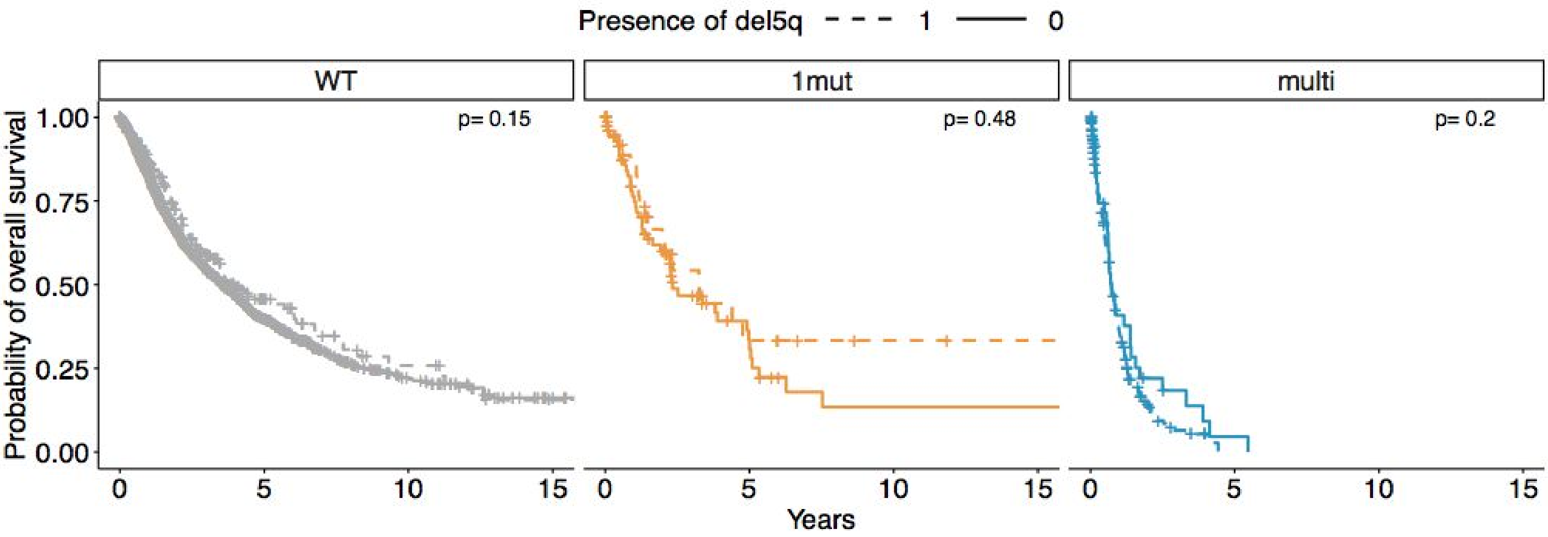

#### Supplementary Figure 12: Multivariate Cox models with *TP53* state alongside IPSS-R risk group

**a.** Results of Cox proportional hazard regression for overall survival (OS). OS is measured from the time of sampling to the death from any cause or to the time of censoring. Explicative variables are the IPSS-R risk group (very-good, good, intermediate is the reference, poor and very-poor) and *TP53* allelic state (mono-allelic, multi-hit and wild-type is the reference). The x-axis is $\log_{10}$ scaled. **b.** Results of cause-specific Cox proportional hazard regression for AML transformation (AMLt) with the same covariates as in a. Time to AML transformation is measured from the time of sampling to the transformation or to the time of censoring, and patients that died without transformation are censored at the time of death. The x-axis is $\log_{10}$ scaled. ****p<0.0001, ***p<0.001, **p<0.01, *p<0.05 Wald test.

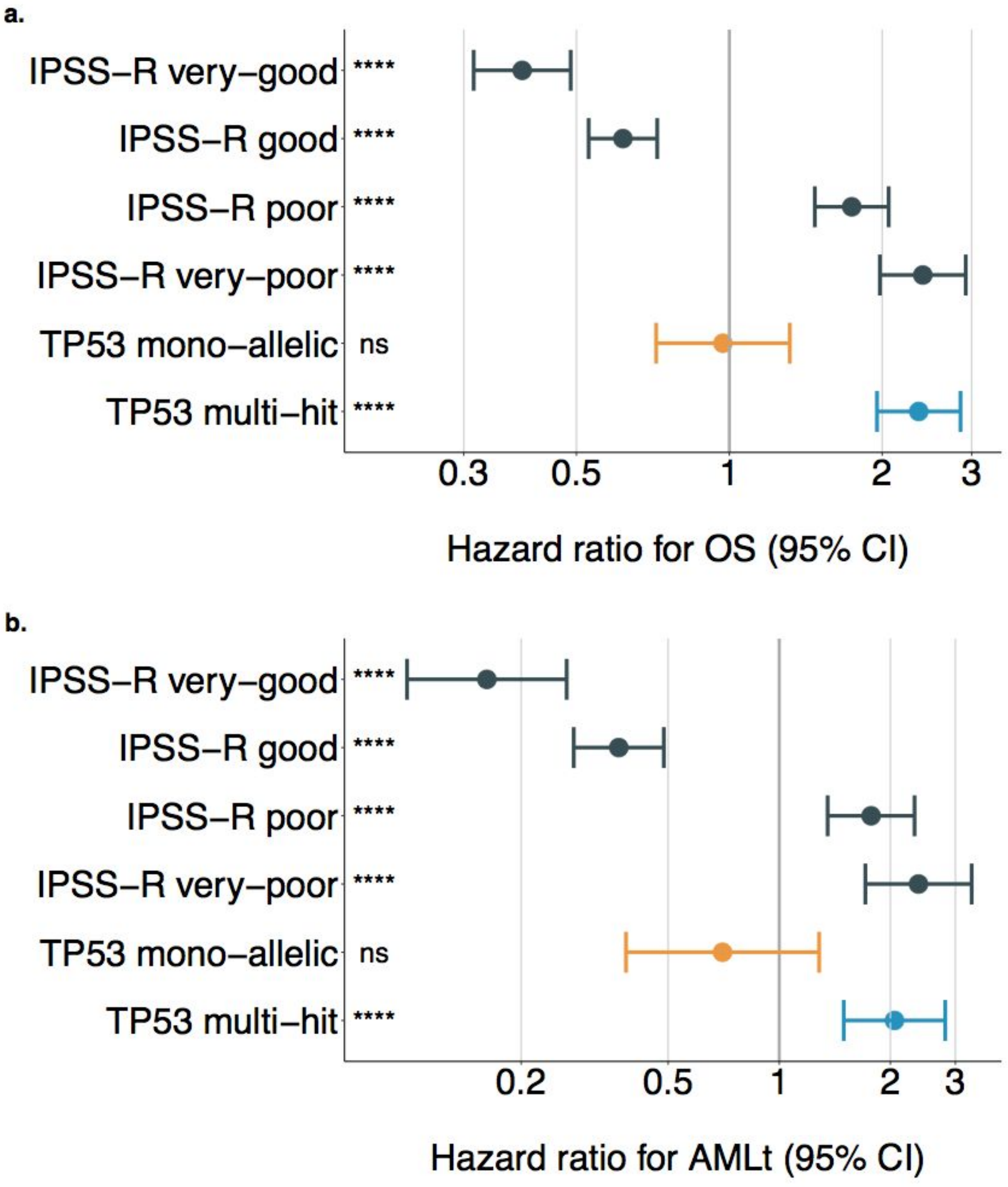

### Supplementary Figure 13: Complex karyotype and *TP53* state

**a.** Interaction between *TP53* state and complex karyotype. 13% (16/125) of mono-allelic *TP53* patients (1mut) have a complex karyotype. Conversely, 91% (231/253) of multiple *TP53* hits patients (multi) have a complex karyotype (91% vs. 13%, OR=70, 95% CI: 33-150, $p<10^{-16}$ Fisher exact test). **b.** Kaplan-Meier probability estimates of overall survival per karyotype status as complex (dashed line) or non-complex (solid line), for cases with multi-hit *TP53* (left panel) or cases with mono-allelic *TP53* (right panel). Multi-hit *TP53* patients with complex karyotype have an increased risk of death compared to multi-hit *TP53* patients without complex karyotype (left panel, HR=3.0, 95% CI: 1.7-5.3, $p<10^{-4}$ Wald test). Mono-allelic *TP53* patients with complex karyotype have an increased risk of death compared to mono-allelic *TP53* patients without complex karyotype (right panel, HR=2.0, 95% CI: 0.99-4.1, p=0.06 Wald test). **c.** Kaplan-Meier probability estimates of overall survival per *TP53* state of wild-type *TP53* (WT), single *TP53* mutation (1mut) and multiple *TP53* hits (multi), for cases with complex karyotype (left panel) or cases without complex karyotype (right panel). Within the cases with complex karyotype (left panel), multi-hit *TP53* patients have an increased risk of death compared to wild-type *TP53* patients (HR=2.3, 95% CI: 1.7-3.2, $p<10^{-6}$ Wald test) and mono-allelic *TP53* patients (HR=2.3, 95% CI: 1.2-4.4, p=0.02 Wald test). Mono-allelic *TP53* patients have similar survival than wild type patients (HR=1.01, 95% CI: 0.48-2.01). Similarly, within the cases with non-complex karyotype (right panel), multi-hit *TP53* patients have an increased risk of death compared to wild-type *TP53* patients (HR=1.9, 95% CI: 1.2-3.2, p=0.01 Wald test) and mono-allelic *TP53* patients (HR=1.6, 95% CI: 0.91-2.9, p=0.1 Wald test). Mono-allelic *TP53* patients have similar survival than wild type patients (HR=1.1, 95% CI: 0.86-1.5). P-values annotated on the panels are from the log-rank test. **c.** Results of Cox proportional hazards regression for overall survival (OS) performed on 3,172 patients with OS data and with 1547 observed death. Explicative variables are the status of the karyotype (complex vs. non-complex) and *TP53* allelic state (mono-allelic, multi-hit and wild-type is the reference). **d.** Results of cause-specific Cox proportional hazards regression for AML transformation (AMLt) performed on 2,875 patients with AMLt data and with 476 observed transformation. Covariates are the same as in b. ****p<0.0001, ***p<0.001, **p<0.01, *p<0.05 Wald test.

a.

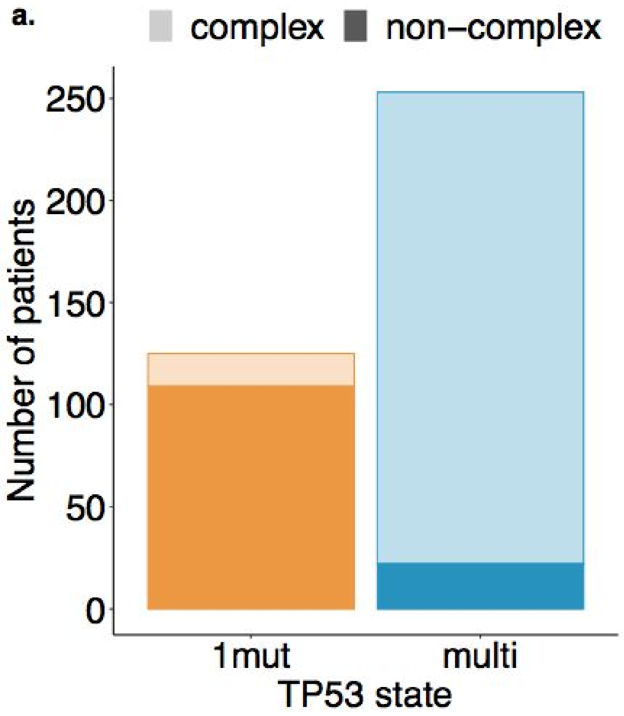

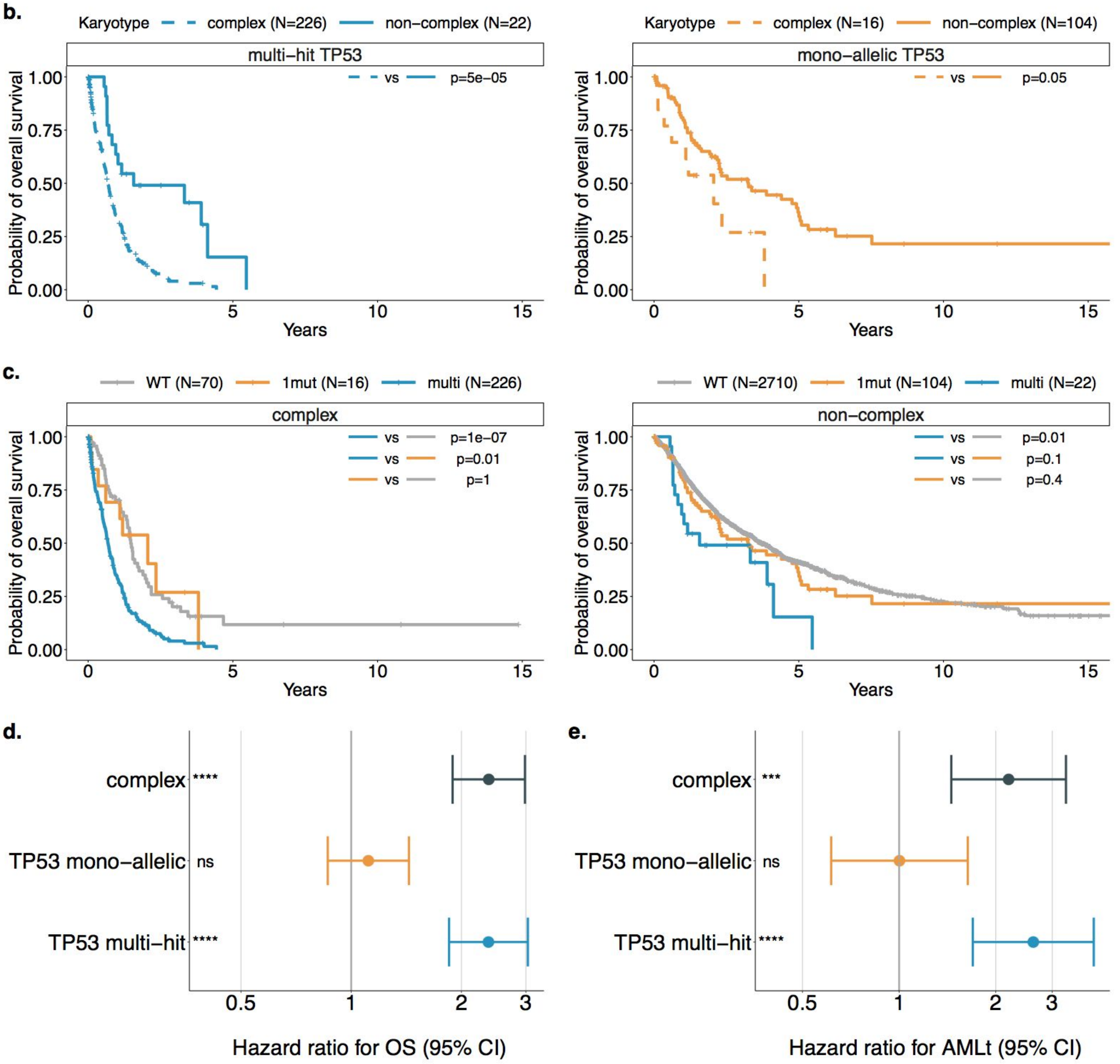
b.
Karyotype
complex (N=226)
non−complex (N=22)
multi−hit TP53
vs
p=5e−05
Probability of overall survival
Years
Karyotype
complex (N=16)
non−complex (N=104)
mono−allelic TP53
vs
p=0.05
c.
WT (N=70)
1mut (N=16)
multi (N=226)
complex
p=1e−07
p=0.01
p=1
WT (N=2710)
1mut (N=104)
multi (N=22)
non−complex
p=0.01
p=0.1
p=0.4
d.
complex ****
TP53 mono−allelic ns
TP53 multi−hit ****
Hazard ratio for OS (95% CI)
e.
complex ***
TP53 mono−allelic ns
TP53 multi−hit ****
Hazard ratio for AMLt (95% CI)

### Supplementary Figure 14: Overall survival per *TP53* state and VAF stratification

Kaplan-Meier probability estimates of overall survival per *TP53* allelic state and different ranges of variant allele frequency (VAF). Optimal VAF cut-points were estimated from the mono-allelic group using the "maxstat" R package (https://cran.r-project.org/web/packages/maxstat/index.html) as implemented in the "survminer" R wrapper package (https://cran.r-project.org/web/packages/survminer/index.html). Patients with mono-allelic *TP53* mutations and VAF>23% (dark orange line) have increased risk of death compared to wild-type patients (HR=2.2, 95% CI: 1.5-3.2, $p<10^{-3}$ Wald test). Multi-hit patients have dismal outcomes across the ranges of VAF. For the multi-hit cases with several mutations, we took at the maximum VAF across the mutations.

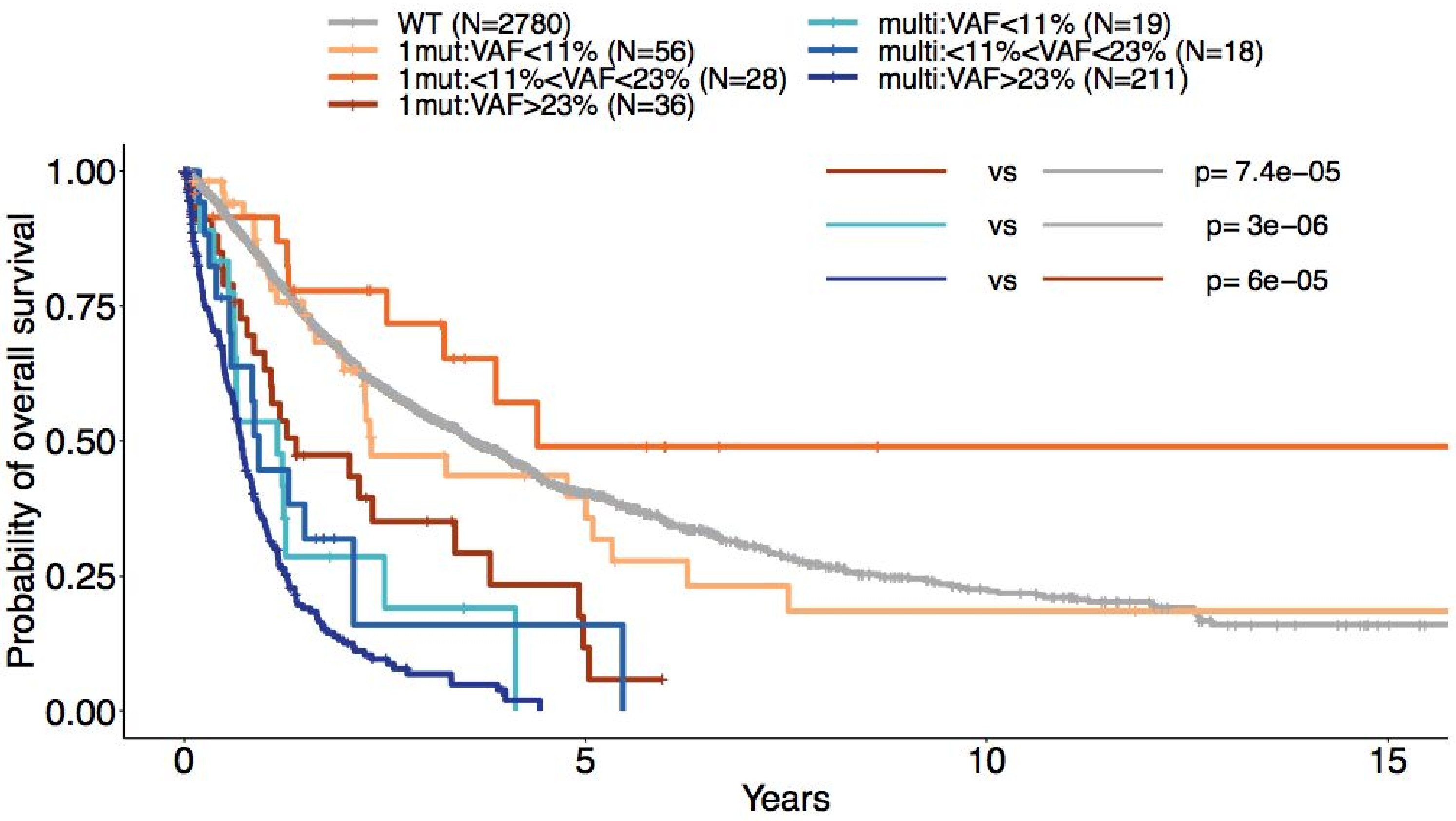

### Supplementary Figure 15: Overall survival per hotspot status and *TP53* state

**a.** Kaplan-Meier probability estimates of overall survival of patients with wild-type *TP53* and with multiple *TP53* hits broken down per hotspot status, i.e., patients with one hotspot mutation alongside another *TP53* alteration (hotspot+other-missense in purple, hotspot+cnloh in pink, hotspot+truncated in orange, hotspot+del in yellow) and patients with multiple hits without hotspot mutation (multi-hit without hotspot, blue). No differences are found between the latter category and any of the hotspot categories. **b.** Kaplan-Meier probability estimates of overall survival of *TP53* wild-type (WT) patients and of multi-hit *TP53* patients with cnloh and truncated mutation (N=23, pink), hotspot mutation at position 273 (N=6, light purple), hotspot mutation at position 248 (N=5, purple) or other missense mutation (N=40, green). **c.** Kaplan-Meier probability estimates of overall survival of *TP53* wild-type (WT) patients and of mono-allelic *TP53* patients with truncated mutations (N=17, pink), with hotspot mutations (N=32, 30 with outcome data, purple) from any position 273, 248, 220 or 175, and with other missense mutations or inframe indels (N=76, 74 with outcome data, green). **d.** Kaplan-Meier probability estimates of overall survival of *TP53* wild-type (WT) patients and of mono-allelic *TP53* patients with hotspot mutations at position 273 (N=13, 11 with outcome data, light purple), 248 (N=11, purple) or 175 (N=7, dark purple). Annotated p-values are from the log-rank test.

**a.**

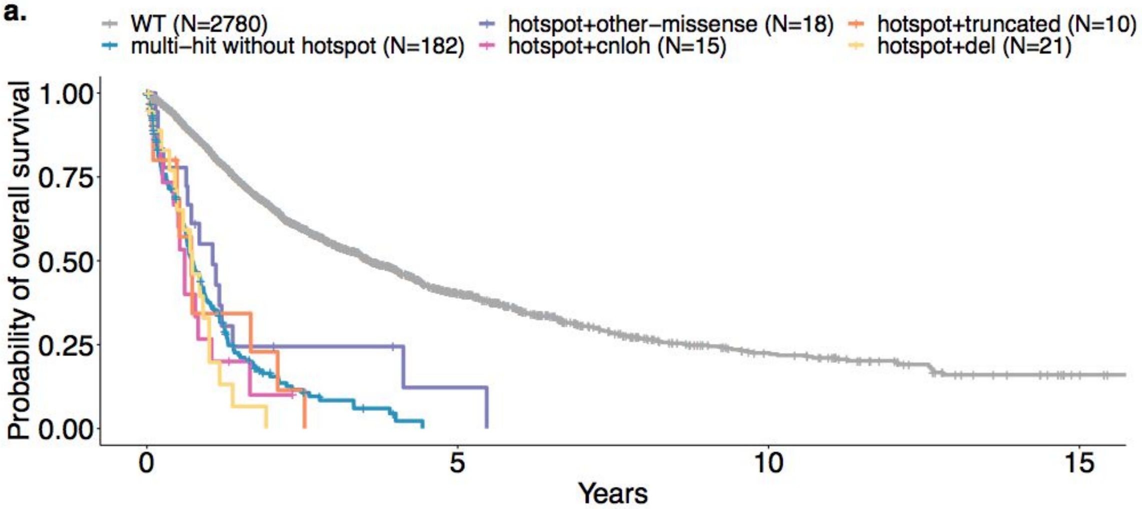


**b.**

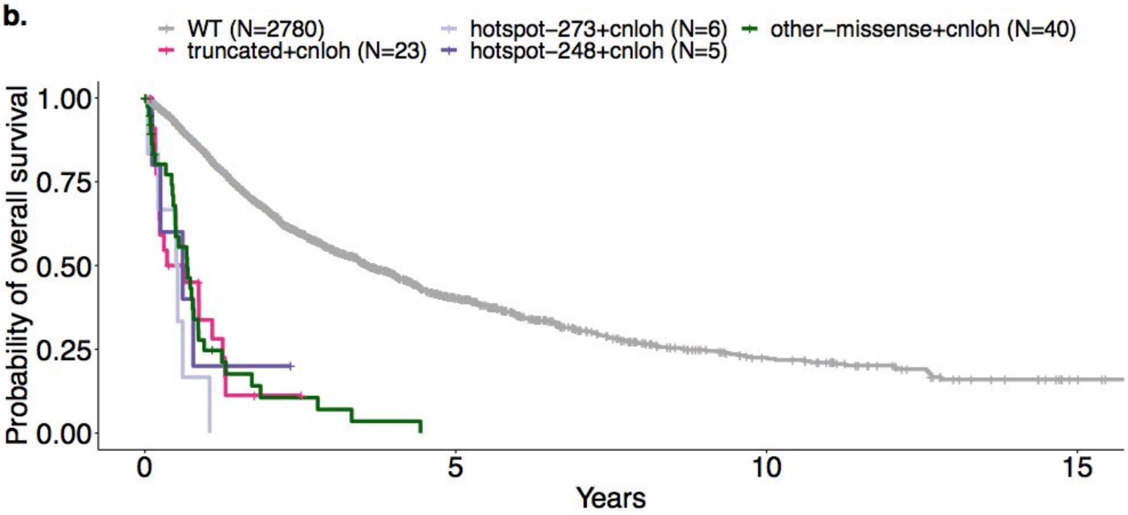

**c.**

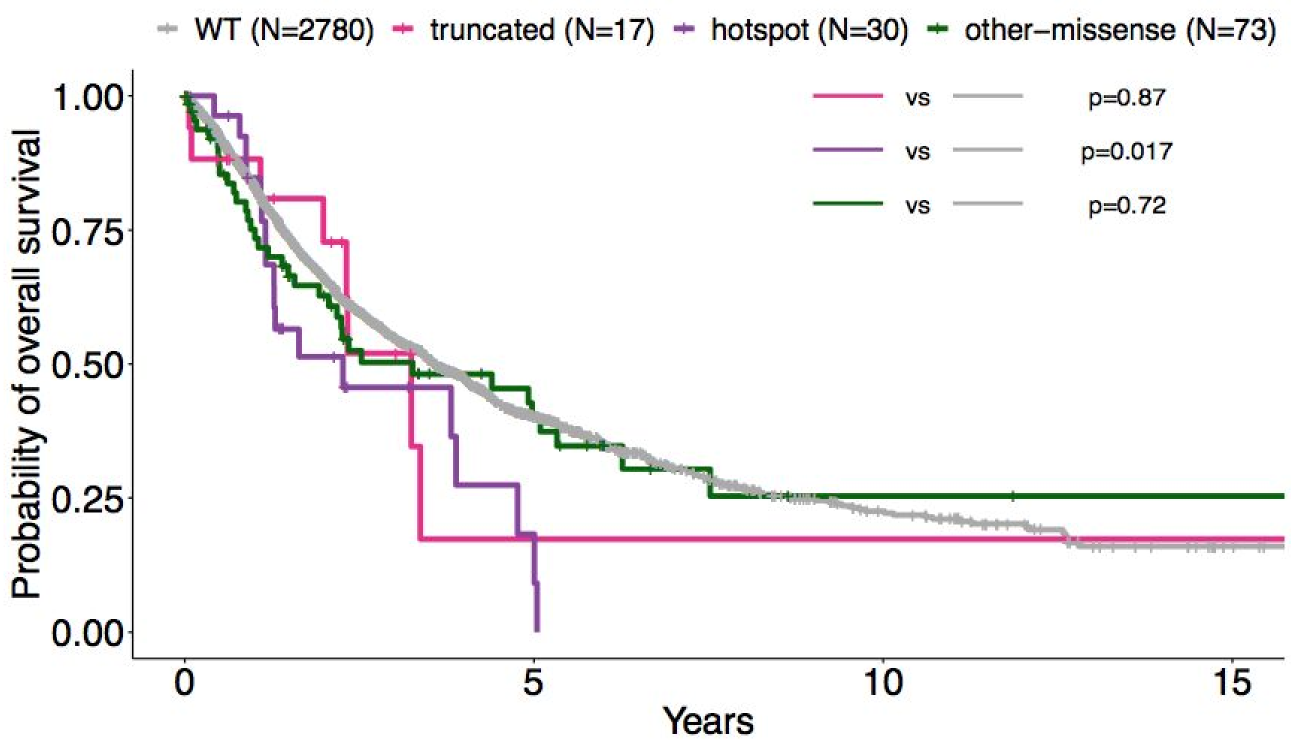
WT (N=2780)
truncated (N=17)
hotspot (N=30)
other−missense (N=73)
vs
p=0.87
vs
p=0.017
vs
p=0.72
Probability of overall survival
1.00
0.75
0.50
0.25
0.00
0
5
10
15
Years

**d.**

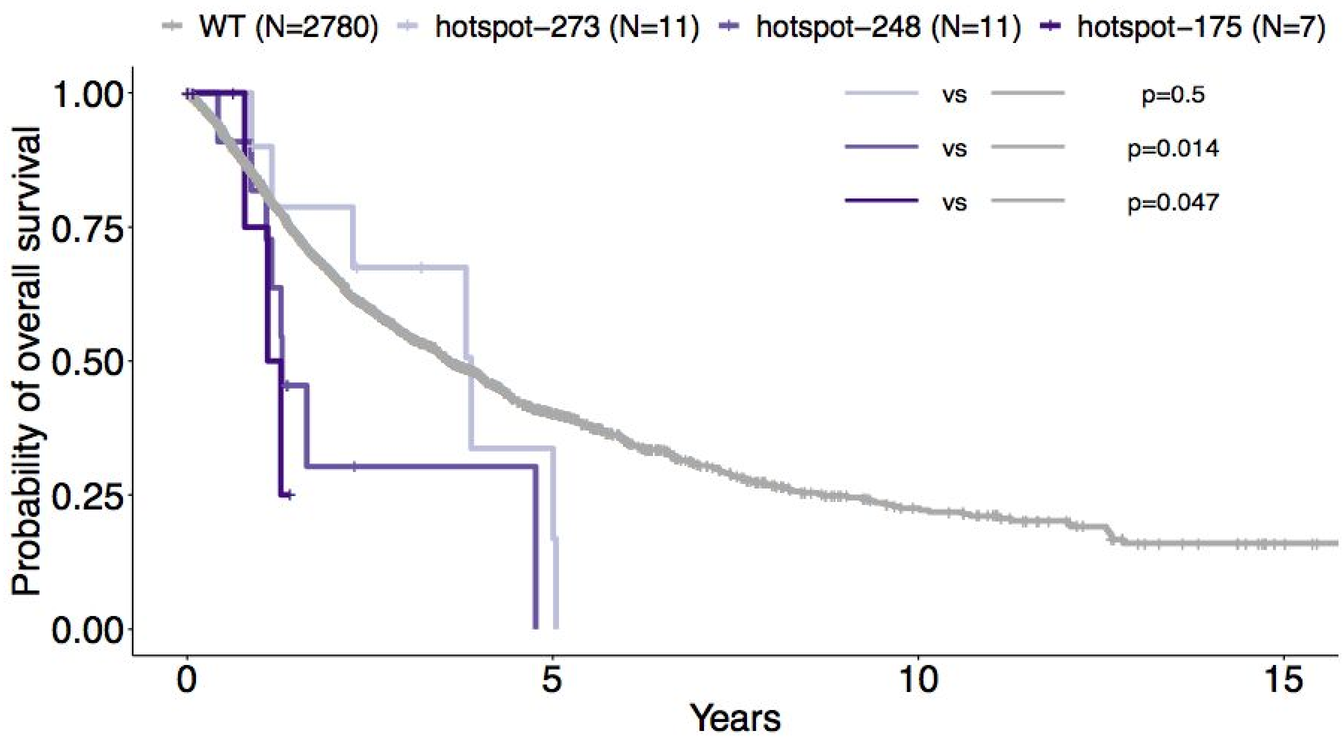
WT (N=2780)
hotspot−273 (N=11)
hotspot−248 (N=11)
hotspot−175 (N=7)
vs
p=0.5
vs
p=0.014
vs
p=0.047
Probability of overall survival
1.00
0.75
0.50
0.25
0.00
0
5
10
15
Years

#### Supplementary Figure 16: Genome stability across de-novo or therapy-related MDS and *TP53* state

Comparison of genome stability across type of chromosomal aberrations, to include rearrangement (rearr), gain and deletion (del), per *TP53* state of single gene mutation (1mut) or multiple hits (multi) and per type of MDS, i.e., de-novo MDS or therapy-related MDS. Annotated p-values are from the Wilcoxon rank-sum test. The y-axis represents the number per patient of unique chromosomes other than 17 with aberrations.

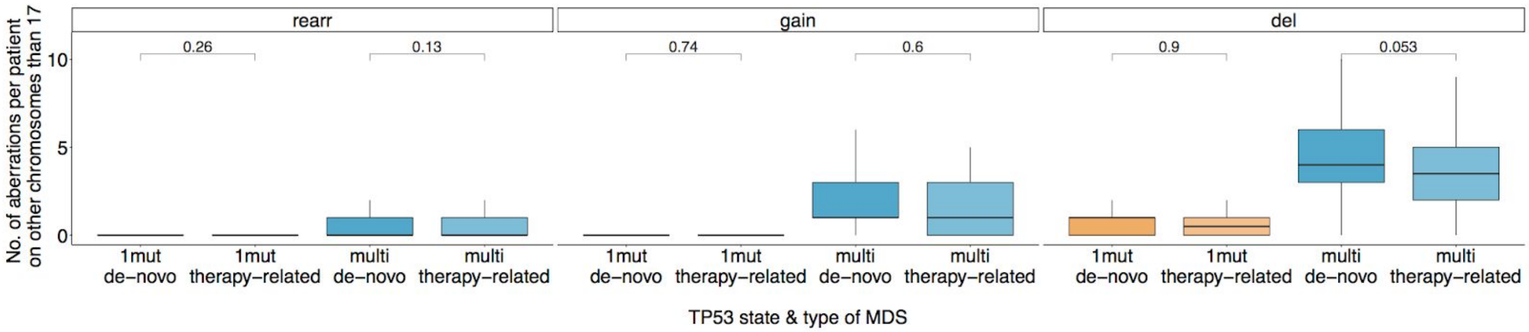

### Supplementary Figure 17: Clonal evolution from MDS to AML of *TP53* mutated patients

We present below the analysis of serial data from 12 MDS patients who progressed to AML with at least one *TP53* mutation in either disease phase. Patient sampling and clinical follow-up for this batch of data are described in (Roman et al. 2016) and in (Smith et al. 2018).

Each following panel corresponds to a patient and describes:

i) The likely oncogenic gene mutations at MDS and AML, with estimated variant allele frequency (VAF) at both disease stages, and corrected VAF to adjust for allele specific copy-numbers (deletion, gain, cnloh).

ii) CBA data at MDS and AML as well as the main chromosomal aberrations extracted from analysis of NGS derived copy-number profiles.

iii) NGS derived CNACS copy-number profiles at MDS and AML.

iv) Scatter plot of corrected VAF of the mutations at MDS (x-axis) and at AML (y-axis).

v) Estimation of clonal dynamics from MDS to AML, with schematics using the so-called fish plots (Miller et al. 2016). Multiple possibilities of clonal evolution leading to the observed pairs of corrected VAFs are depicted as multiple fish plots.

#### a. Patient 128470

The patient had multiple mutations at MDS and AML, including two clonal *TP53* mutations (i). The karyotype was complex at MDS and AML with deletions of 3p, 5q, 7 and 18p (ii and iii). The founding clone is characterized by bi-allelic targeting of *TP53* from two mutations, one missense C238Y mutation and one splice site mutation. It gave rise to two subclones, both with distinct *SETBP1* mutations (note that we could phase the two *SETBP1* mutations and they were observed in trans). One of the *SETBP1* subclone with *KRAS* A59 hotspot mutation shrank from MDS to AML; whereas the other *SETBP1* subclone containing itself two other subblones with *NF1* mutation and *NRAS* Q61 hotspot mutation expanded from MDS to AML. Small subclones with *PTPN11* and *CBL* mutations could originate from multiple parent clones. The multiple possibilities representing the clonal dynamics from MDS to AML are represented in v).

i)

| Gene | Protein change | Variant key | MDS VAF | MDS corrected VAF | AML VAF | AML corrected VAF |
|---|---|---|---|---|---|---|
| NRAS | p.Q61R | 1_115256529_T_C | 0.00 | 0.00 | 0.21 | 0.21 |
| CBL | p.? | 11_119149009_T_G | 0.04 | 0.04 | 0.01 | 0.01 |
| PTPN11 | p.N58Y | 12_112888156_A_T | 0.06 | 0.06 | 0.02 | 0.02 |
| KRAS | p.A59T | 12_25380283_C_T | 0.15 | 0.15 | 0.05 | 0.05 |
| NF1 | p.G629R | 17_29552152_G_A | 0.06 | 0.06 | 0.09 | 0.09 |
| TP53 | p.? | 17_7577156_C_T | 0.40 | 0.40 | 0.41 | 0.41 |
| TP53 | p.C238Y | 17_7577568_C_T | 0.43 | 0.43 | 0.40 | 0.40 |
| SETBP1 | p.D868N | 18_42531907_G_A | 0.25 | 0.25 | 0.12 | 0.12 |
| SETBP1 | p.G870S | 18_42531913_G_A | 0.18 | 0.18 | 0.29 | 0.29 |

ii)

| MDS NGS profile | MDS Cytogenetics | AML NGS profile | AML Cytogenetics |
|---|---|---|---|
| del3p,del5q,del7,del18p | missing | del3p,del5q,del7,del18p | 44,XY,der(3;18)(q10;q10),del(5)(q22q35),−7[10]/46,XY[1] |

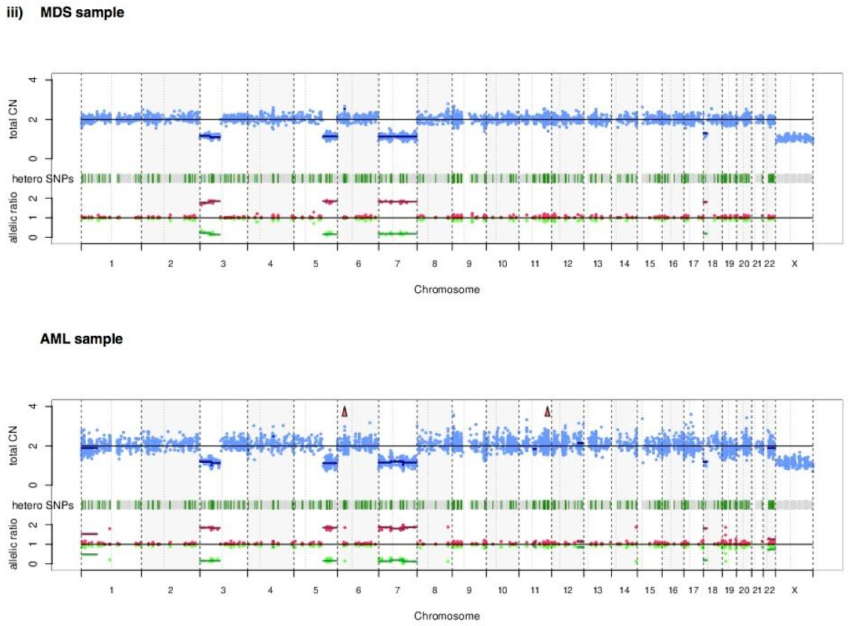


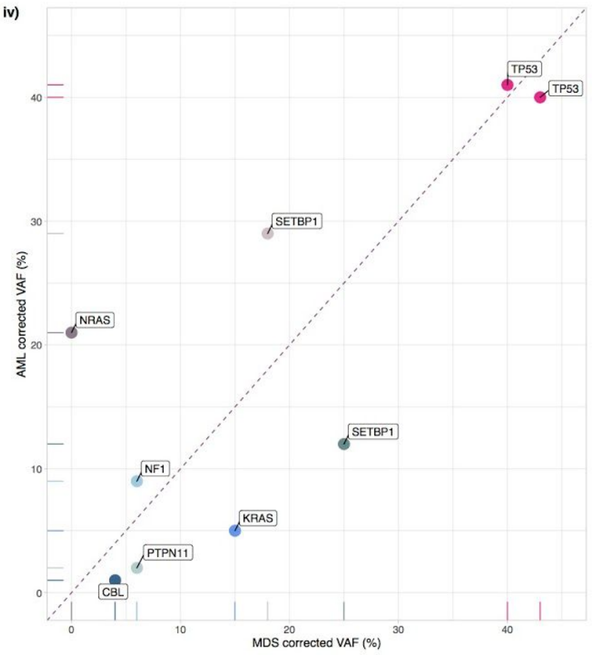

v)

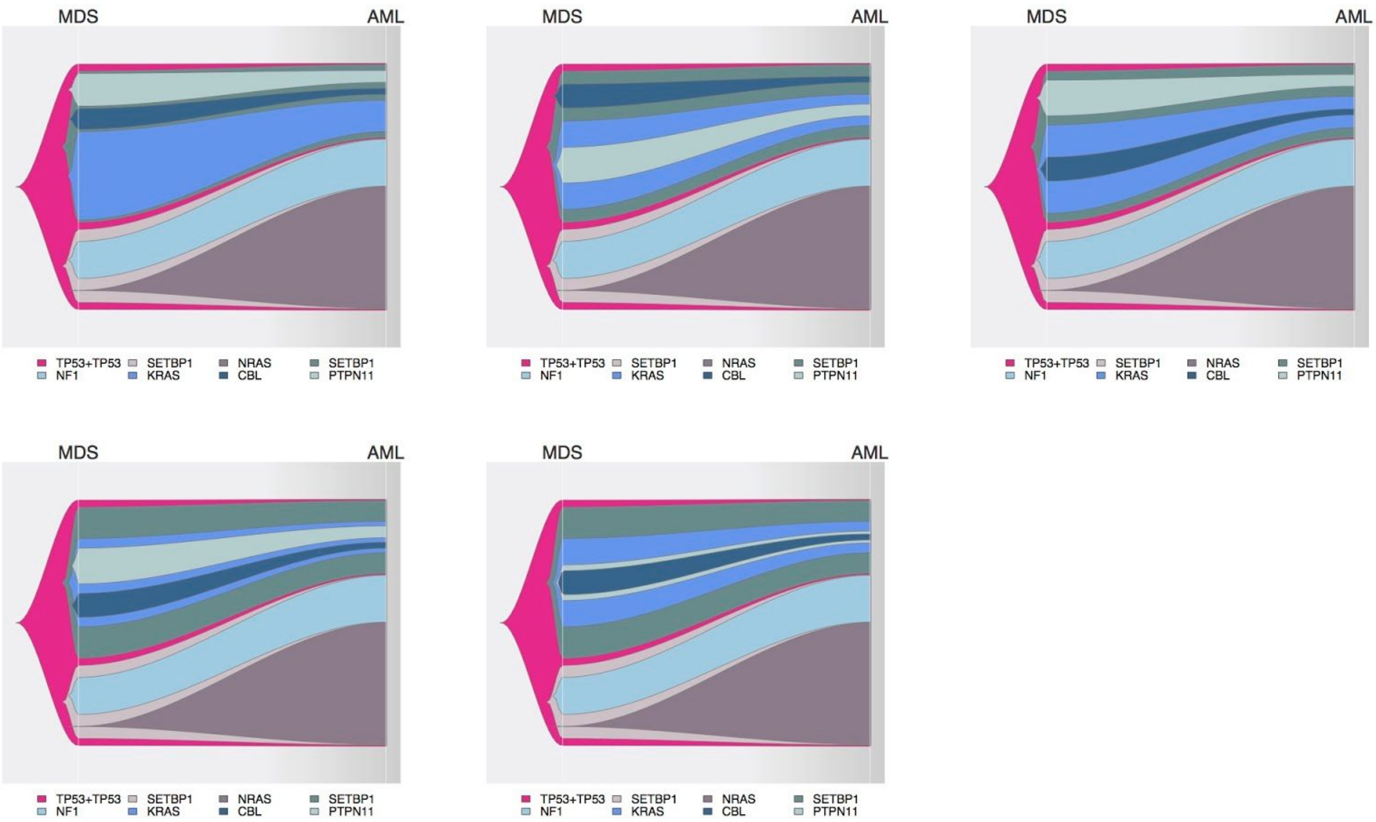
MDS
AML
TP53+TP53
SETBP1
NRAS
SETBP1
NF1
KRAS
CBL
PTPN11
MDS
AML
TP53+TP53
SETBP1
NRAS
SETBP1
NF1
KRAS
CBL
PTPN11
MDS
AML
TP53+TP53
SETBP1
NRAS
SETBP1
NF1
KRAS
CBL
PTPN11
MDS
AML
TP53+TP53
SETBP1
NRAS
SETBP1
NF1
KRAS
CBL
PTPN11
MDS
AML
TP53+TP53
SETBP1
NRAS
SETBP1
NF1
KRAS
CBL
PTPN11

##### b. Patient 129056

The patient had clonal mutations in *KMT2D* and *TP53* (i). The karyotype was complex at MDS and AML with deletions at 5q, 18 and 21 (ii and iii). Copy-neutral loss of heterozygosity (cnloh) of 17p was observed at both MDS and AML (iii), and *TP53* VAFs were corrected accordingly (i and iv). The founding clone is characterized by bi-allelic targeting of *TP53* with one point mutation at residue 220 and loss of the second wild-type allele by cnloh.

i)

| Gene | Protein change | Variant key | MDS VAF | MDS corrected VAF | AML VAF | AML corrected VAF |
|---|---|---|---|---|---|---|
| KMT2D | p.P496fs*3 | 12_49445979_GGA_G | 0.40 | 0.40 | 0.43 | 0.43 |
| TP53 | p.Y220C | 17_7578190_T_C | 0.72 | 0.36 | 0.85 | 0.43 |

ii)

| MDS NGS profile | MDS Cytogenetics | AML NGS profile | AML Cytogenetics |
|---|---|---|---|
| del5q,del18,plus21q,cnloh17p | 45~50,XY,del(5)(q?13q3),−7,−18,−21,+2~7mar[cp11] | del5q,del18,plus21q,cnloh17p | missing |

iii) MDS sample

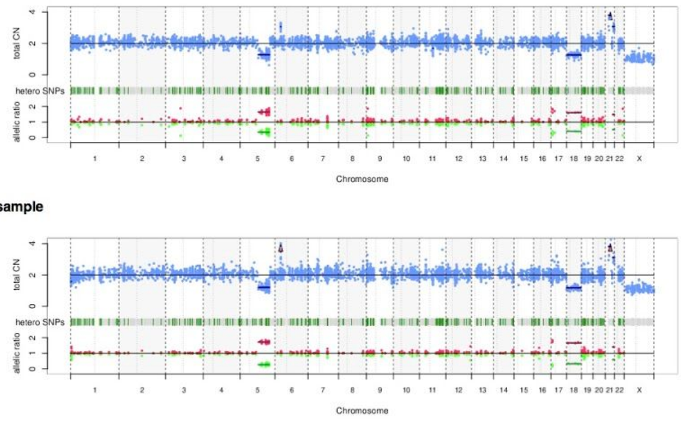


iv)

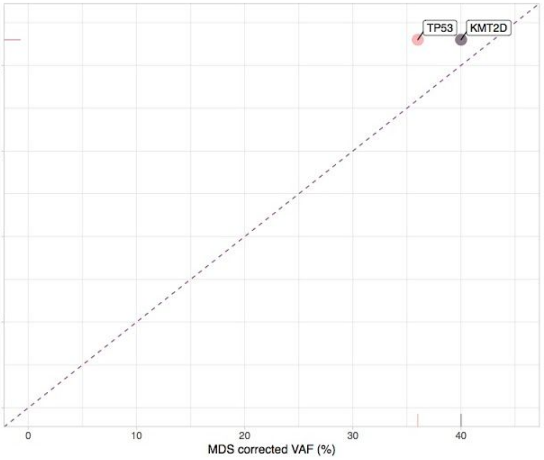


v)

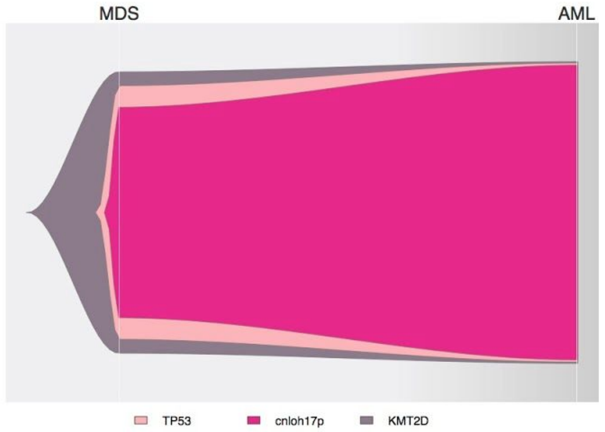

**c. Patient 129363**

The patient had clonal mutations in *DNMT3A* and *TP53* (i). The karyotype was complex at MDS and AML with multiple deletions (ii and iii). Deletion of 17p at the *TP53* locus was observed at both MDS and AML (iii), and *TP53* VAFs were corrected accordingly (i and iv). The founding clone is characterized by bi-allelic targeting of *TP53* with one point mutation at residue 238 and loss of the other allele by deletion. It gave rise to subclonal *KRAS* hotspot G12 mutation and *PTPN11* mutation. Two clonal dynamics possibilities are depicted in v).

**i)**

| Gene | Protein change | Variant key | MDS VAF | MDS corrected VAF | AML VAF | AML corrected VAF |
|---|---|---|---|---|---|---|
| PTPN11 | p.S502P | 12_112926884_T_C | 0.04 | 0.04 | 0.06 | 0.06 |
| KRAS | p.G12V | 12_25398284_C_A | 0.03 | 0.02 | 0.09 | 0.04 |
| TP53 | p.C238Y | 17_7577568_C_T | 0.77 | 0.38 | 0.91 | 0.45 |
| DNMT3A | p.Y528* | 2_25467492_G_C | 0.41 | 0.41 | 0.40 | 0.40 |

**ii)**

| MDS NGS profile | MDS Cytogenetics | AML NGS profile | AML Cytogenetics |
|---|---|---|---|
| del5q,del7,plus8,del12,del17p | 44~46,XY,add(4)(q3?),del(5)(q13q33),−7,add(7)(q?22), +8,−11,−12,del(13)(q12q22),−16,−17, −19,?add(21)(p11),+2~4mar[cp10] | del5q,del7,plus8,del12,del17p | 45~54,XY,add(4)(q3?),del(q13q33),−7,add(7)(q?22).+8, −12,del(13)(q12q22),der(17)t(7;17)(q11 23;p13), +mar[cp9]/46,XY[1] |

**iii)** MDS sample

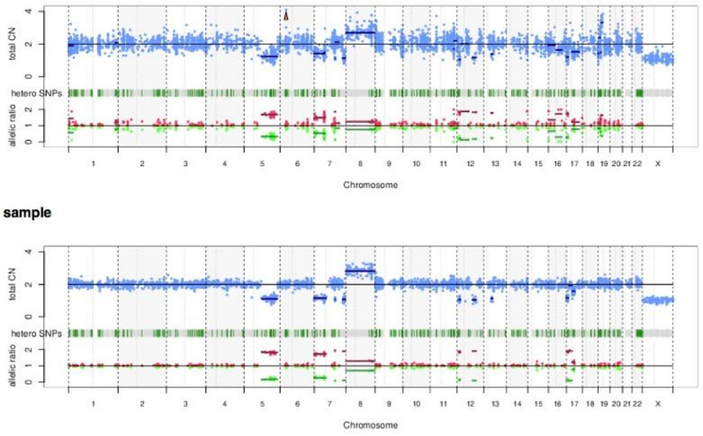


**iv)**

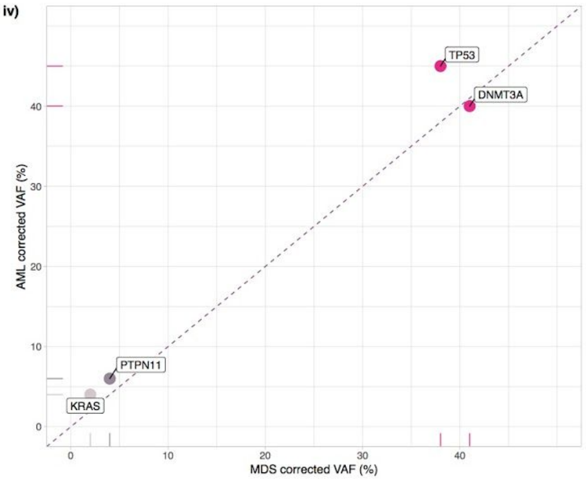


**v)**

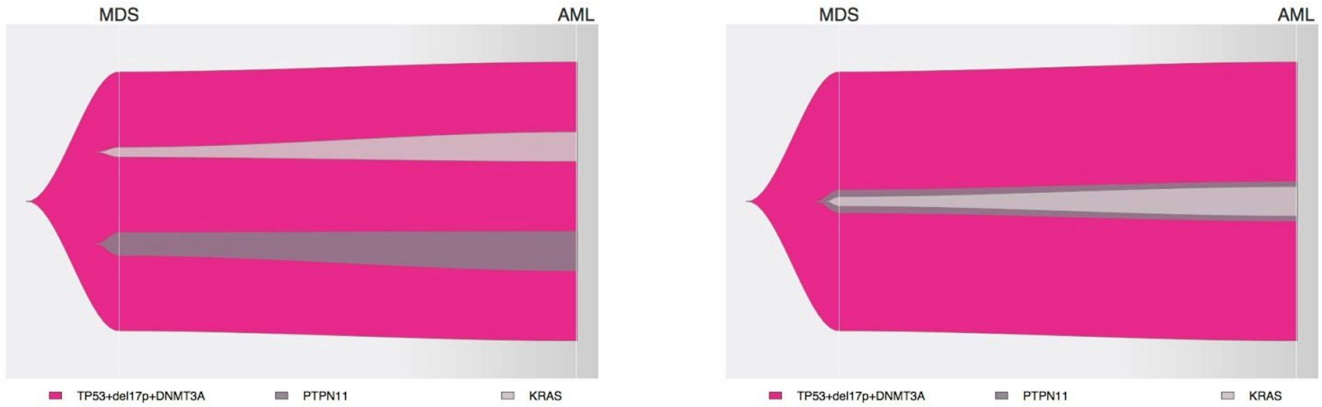

##### d. Patient 129525

The patient had a *TP53* mutation at MDS and AML and a *NF1* mutation observed at AML only with small VAF (i). The karyotype was complex at MDS and AML with multiple deletions (ii and iii). Deletion of 17p at the *TP53* locus was observed at both MDS and AML (iii), and *TP53* VAFs were corrected accordingly (i and iv). The founding clone is characterized by bi-allelic targeting of *TP53* with one point mutation at residue 193 and loss of the other allele by chromosomal deletion. It gave rise to a subclone with additional *NF1* mutation observed at AML.

i)

| Gene | Protein change | Variant key | MDS VAF | MDS corrected VAF | AML VAF | AML corrected VAF |
|---|---|---|---|---|---|---|
| NF1 | p.R1534* | 17_29588751_C_T | 0.00 | 0.00 | 0.08 | 0.04 |
| TP53 | p.H193L | 17_7578271_T_A | 0.63 | 0.32 | 0.83 | 0.42 |

ii)

| MDS NGS profile | MDS Cytogenetics | AML NGS profile | AML Cytogenetics |
|---|---|---|---|
| del5q,del16q,del17p,del18,plus22 | 40~41,X,–Y,–5,–6,–8,–15,–16,–17, –18,–21,–22,–22,+6mar[cp7]/46,XY[1] | del3q,del5q,del7,del16q,del17p,del18,plus22 | 40~21,X,–Y,–3,–5,–6,–7,–8,–15,–16,–17,–18, 21,–22,–22,+6~11mar[cp10]/46,XY[1] |

iii)

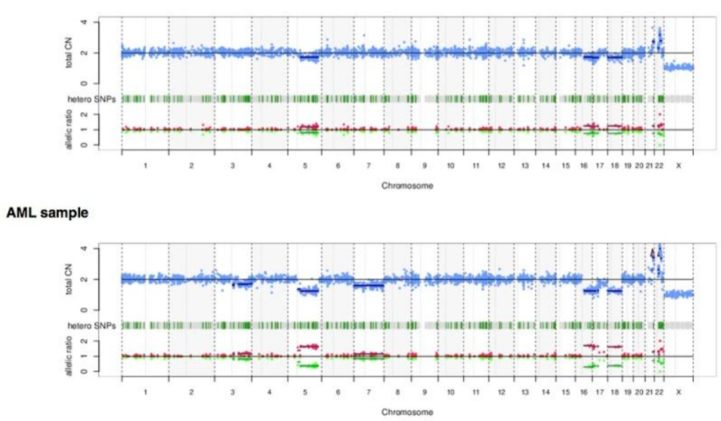


iv)

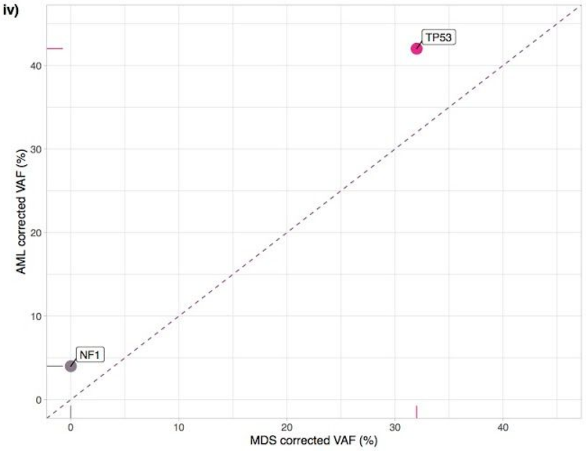


v)

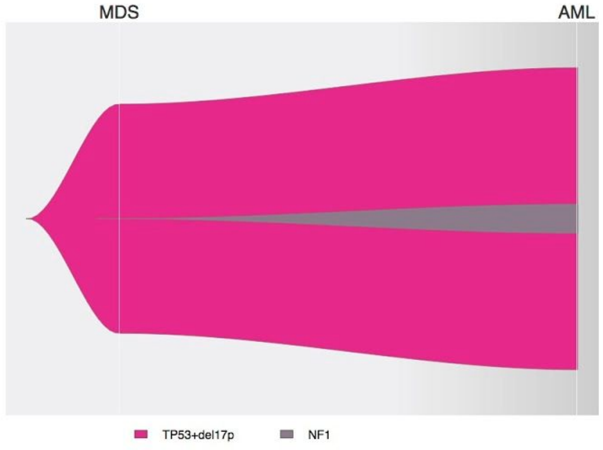

##### e. Patient 129602

The patient had two clonal *TP53* mutations at MDS and AML. It also had a subclonal *U2AF1* Q157 mutation in MDS that became clonal in AML (i). Three chromosomal aberrations were observed at both MDS and AML, i.e., plus1p, del5q and del22q (ii and iii). The founding clone is characterized by bi-allelic targeting of *TP53* with two missense mutations at residues 126 and 272. It gave rise to a *U2AF1* subclone with hotspot Q157 mutation that expanded in the course of the disease.

i)

| Gene | Protein change | Variant key | MDS VAF | MDS corrected VAF | AML VAF | AML corrected VAF |
|---|---|---|---|---|---|---|
| TP53 | p.V272M | 17_7577124_C_T | 0.46 | 0.46 | 0.43 | 0.43 |
| TP53 | p.Y126H | 17_7578554_A_G | 0.44 | 0.44 | 0.48 | 0.48 |
| U2AF1 | p.Q157R | 21_44514777_T_C | 0.26 | 0.26 | 0.42 | 0.42 |

ii)

| MDS NGS profile | MDS Cytogenetics | AML NGS profile | AML Cytogenetics |
|---|---|---|---|
| plus1p,del5q,del22q | 45,XY,add(1)(p?3),del(5)(q13),−22[3]/46,idem,+r[7] | plus1p,del5q,del22q | 45,XY,add(1)(p?3),del(5)(q13),−22[3]/46,idem,+r[7] |

iii) **MDS sample**

**AML sample**

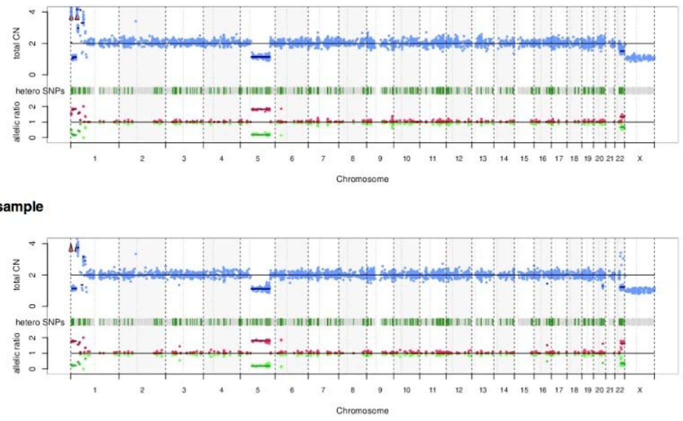


iv)

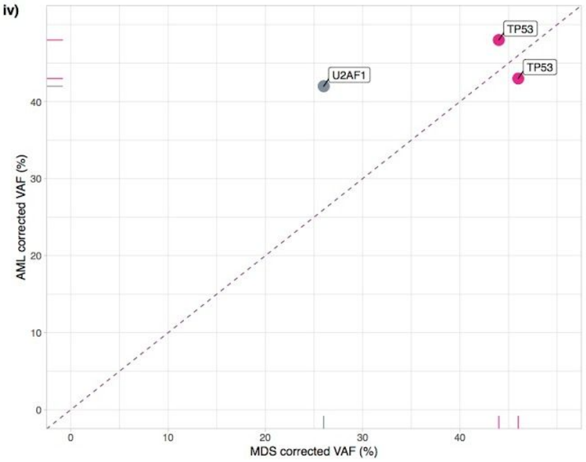


v)

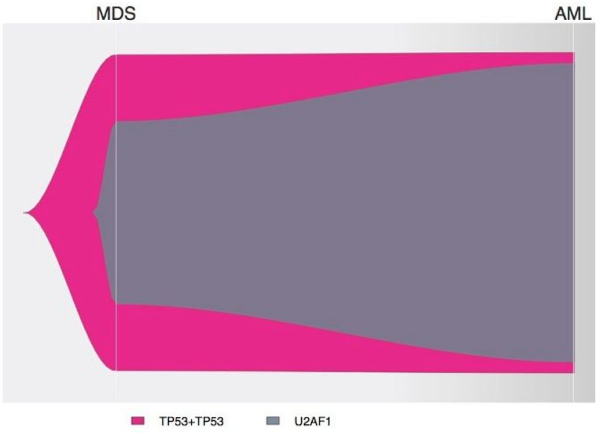

##### f. Patient 129662

The patient had a *TP53* mutation at MDS and AML and a *SRSF2* mutation observed at AML only with small VAF (i). The karyotype was complex at MDS and AML with multiple deletions (ii and iii). Copy-neutral loss of heterozygosity (cnloh) of 17p was observed at both MDS and AML (iii), and *TP53* VAFs were corrected accordingly (i and iv). The founding clone is characterized by bi-allelic targeting of *TP53* with one frameshift deletion and loss of the remaining wild-type allele by cnloh. It gave rise to a subclone with additional *SRSF2* mutation observed at AML. Note that the *SRSF2* mutation is not the canonical P95H/L/R hotspot.

i)

| Gene | Protein change | Variant key | MDS VAF | MDS corrected VAF | AML VAF | AML corrected VAF |
|---|---|---|---|---|---|---|
| SRSF2 | p.D97fs*27 | 17_74732955_C_CG | 0.00 | 0.00 | 0.07 | 0.07 |
| TP53 | p.Q167fs*3 | 17_7578428_GC_G | 0.78 | 0.39 | 0.93 | 0.47 |

ii)

| MDS NGS profile | MDS Cytogenetics | AML NGS profile | AML Cytogenetics |
|---|---|---|---|
| del5q,del6p,del12q,del15q, del16q,cnloh17p | 45,XX,del(5)(q13q33),del(6)(p33), −12,−15,−16,+2mar[10] | del5q,del6p,del7,del9q,del12q, del15q,del16q,cnloh17p | 45,XX,del(5)(q13q33),del(6)(p33),−12, −15,−16,+2mar[10] |

iii) MDS sample

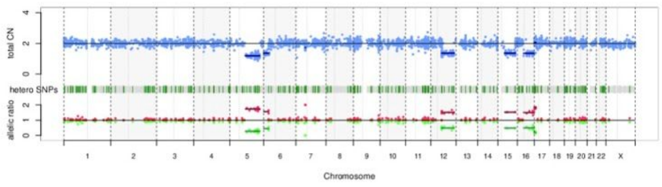


AML sample

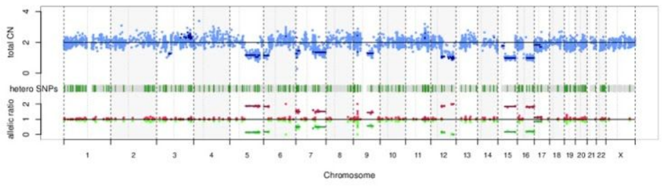


iv)

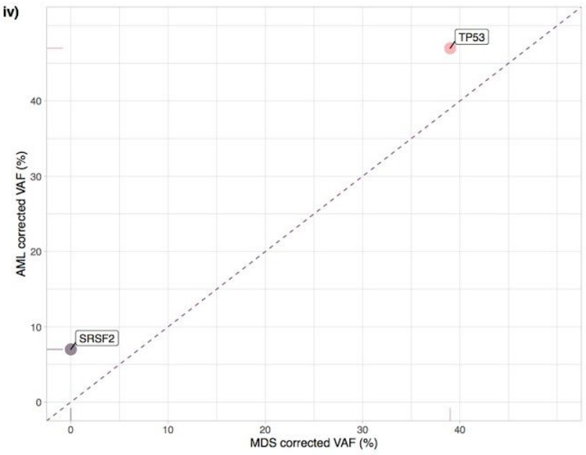


v)

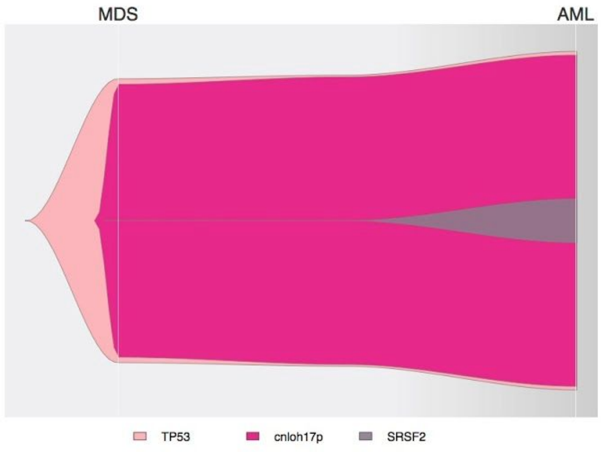

#### g. Patient 129763

The patient had 3 mutations at MDS and AML, one clonal *TP53* missense K139E mutation and additional mutations in *DNMT3A* and *NF1*. The karyotype was complex at MDS and AML, with multiple deletions alongside trisomy 8. Trisomy 11 is only observed at AML. Deletions of 17p at the *TP53* locus and at the *NF1* locus were observed at both MDS and AML (iii), and VAFs were corrected accordingly (i and iv). The founding clone is characterized by bi-allelic targeting of *TP53* with one point mutation at residue 139 and loss of the other allele by chromosomal deletion. It gave rise to nested *DNMT3A* and *NF1* subclones. Note that the subclone with *NF1* shrinks from MDS to AML as a subclone with trisomy 11 is likely to increase.

i)

| Gene | Protein change | Variant key | MDS VAF | MDS corrected VAF | AML VAF | AML corrected VAF |
|---|---|---|---|---|---|---|
| NF1 | p.R192* | 17_29497003_C_T | 0.56 | 0.28 | 0.17 | 0.08 |
| TP53 | p.K139E | 17_7578515_T_C | 0.93 | 0.46 | 0.89 | 0.44 |
| DNMT3A | p.E865* | 2_25458580_C_A | 0.36 | 0.36 | 0.50 | 0.50 |

ii)

| MDS NGS profile | MDS Cytogenetics | AML NGS profile | AML Cytogenetics |
|---|---|---|---|
| del5q,plus8,del16q,del17p | missing | del5q,plus8,plus11,del12q,del16q,del17p | missing |

iii) **MDS sample**

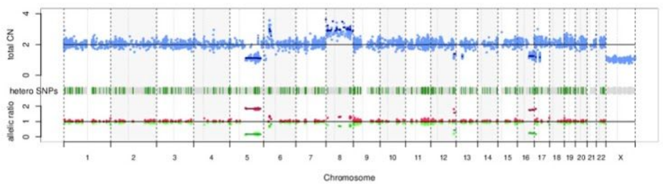


**AML sample**

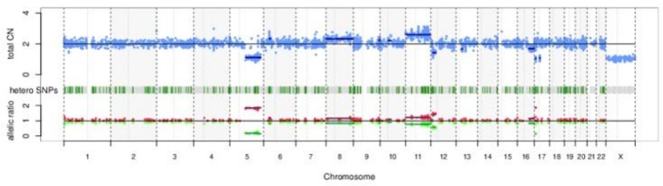


iv)

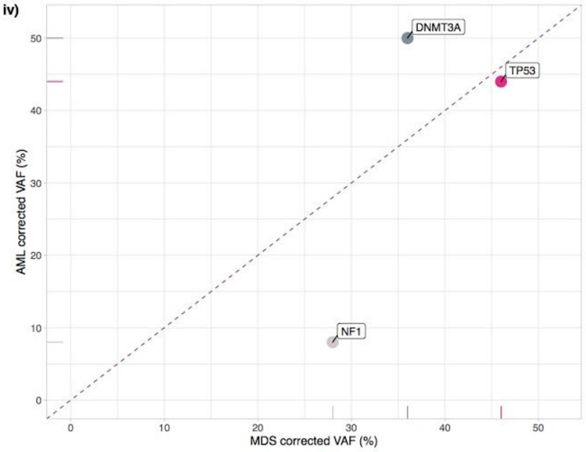


v)

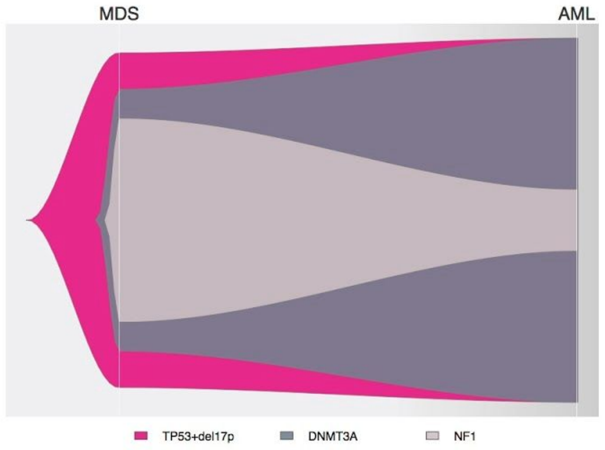

##### h. Patient 128715

The patient had a clonal *EZH2* mutation in MDS and AML, and a subclonal *TP53* mutation at AML only (i). Copy-neutral loss of heterozygosity (cnloh) of 7q was observed at MDS and AML (iii), and *EZH2* VAF was corrected accordingly (i and iv). No other chromosomal aberration was observed at MDS, whilst the karyotype was complex at AML with multiple gains and deletion of chromosome 16. Deletion of 17p at the *TP53* locus was present at AML (iii), and *TP53* VAF was corrected accordingly (i and iv). The founding clone is characterized by bi-allelic targeting of *EZH2* with point mutation at residue 288 and loss of the wild-type allele by cnloh. It gave rise to a subclone with *TP53* mutation and deletion of the *TP53* locus, i.e., bi-allelic targeting of *TP53*, in addition to multiple chromosomal alterations.

i)

| Gene | Protein change | Variant key | MDS VAF | MDS corrected VAF | AML VAF | AML corrected VAF |
|---|---|---|---|---|---|---|
| TP53 | p.R248W | 17_7577539_G_A | 0.00 | 0.00 | 0.52 | 0.26 |
| EZH2 | p.R288Q | 7_148523590_C_T | 0.88 | 0.44 | 0.93 | 0.47 |

ii)

| MDS NGS profile | MDS Cytogenetics | AML NGS profile | AML Cytogenetics |
|---|---|---|---|
| cnloh7q | missing | plus1,plus2,plus6,cnloh7q,plus8,plus13q, plus14q,plus15q,del16q,del17p,plus19,plus20,plusX | 54~56,XX,+X,+1,+1,?add(1)(p13),+2,+6,+8,+13, +14,+15,−16,der(17;19)(q10;q10)x2,+17,+19,+20,+r[cp8]/46,XX[2] |

iii) MDS sample

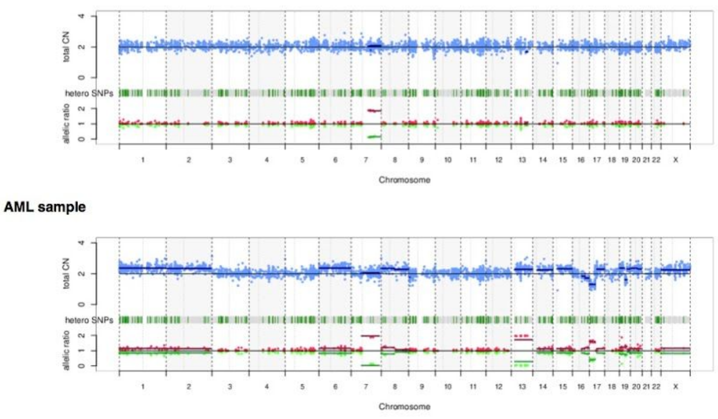


iv)

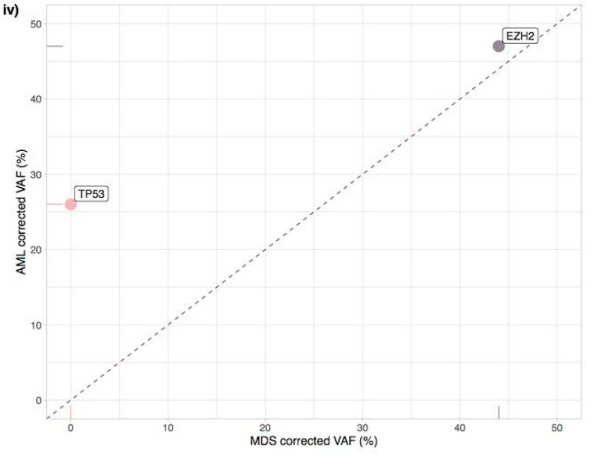


v)

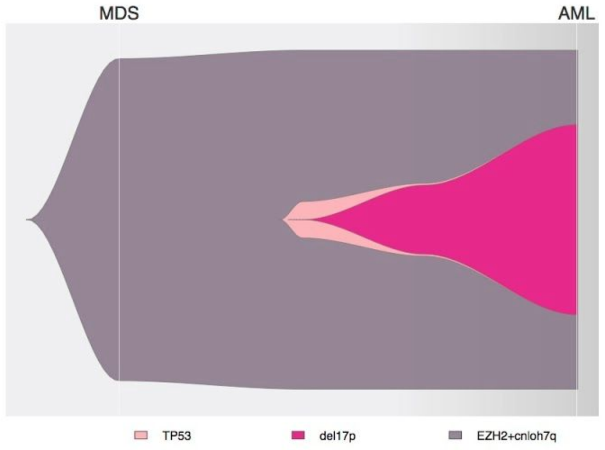

##### i. Patient 128829

The patient had a clonal *U2AF1* mutation at MDS and AML, along with an *ASXL1* mutation with much higher VAF at MDS than AML, *TET2* and *CBL* mutations with higher VAFs in MDS than AML as well, and *TP53* mutation with much higher VAF in AML than MDS (i). The karyotype was normal at MDS and complex at AML with multiple deletions (ii and iii). Deletion of 17p at the *TP53* locus was present at AML (iii), and *TP53* VAF was corrected accordingly (i and iv). The two possibilities shown in v) for clonal dynamics illustrate inter-clonal competition and attainment of clonal dominance of the clone with bi-allelic targeting of *TP53* with one point mutation at residue 245 and loss of the other allele by chromosomal deletion.

i)

| Gene | Protein change | Variant key | MDS VAF | MDS corrected VAF | AML VAF | AML corrected VAF |
|---|---|---|---|---|---|---|
| CBL | p.C384Y | 11_119148931_G_A | 0.19 | 0.19 | 0.01 | 0.01 |
| TP53 | p.G245C | 17_7577548_C_A | 0.05 | 0.05 | 0.86 | 0.43 |
| ASXL1 | p.C730fs1 | 20_31022703_TGCCTACTACA_T | 0.43 | 0.43 | 0.28 | 0.14 |
| U2AF1 | p.Q157H | 21_44514776_C_A | 0.47 | 0.47 | 0.47 | 0.47 |
| TET2 | p.D945fs8 | 4_106157930_AG_A | 0.1 | 0.1 | 0 | 0 |

ii)

| MDS NGS profile | MDS Cytogenetics | AML NGS profile | AML Cytogenetics |
|---|---|---|---|
| NK | missing | del7,del9q,del11,del12,del15q,del16,del17p,del18,del20 | missing |

iii) MDS sample

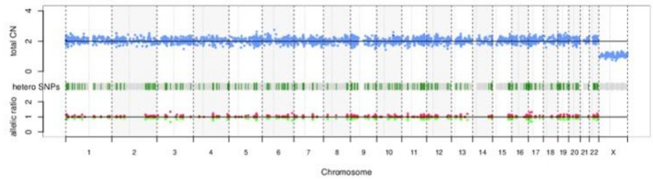


AML sample

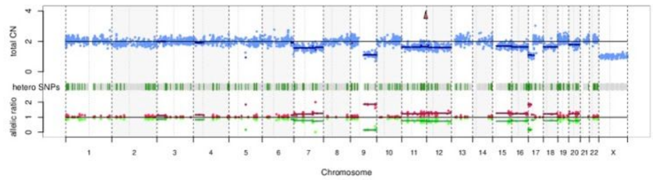


iv)

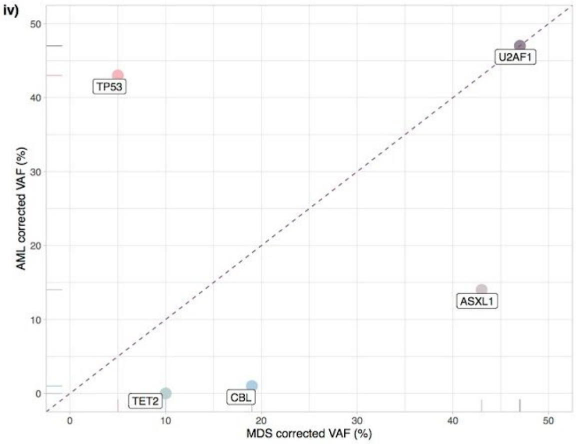


v)

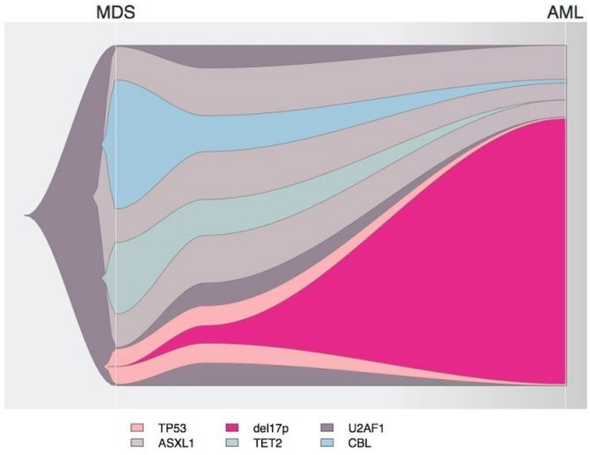


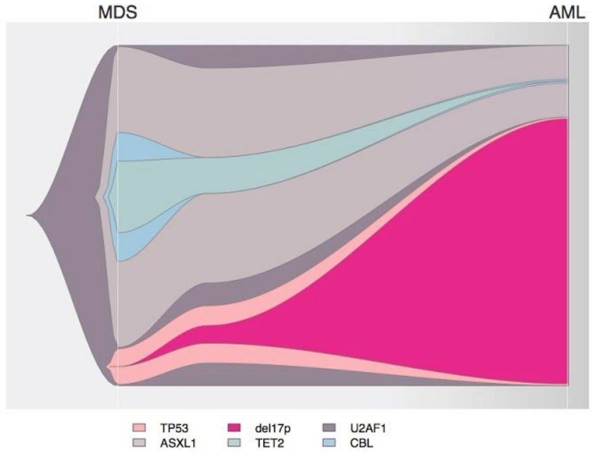

##### j. Patient 129119

The patient had clonal mutations in *TET2*, *DNMT3A* and *TP53* (i). The karyotype was complex at MDS and AML with multiple deletions (ii and iii). Loss of heterozygosity of 17p at *TP53* locus was difficult to assess here but was suggested by the CNACS profiles at a subclonal level (iii). The proposed clonal dynamics shown in v) is therefore a founding clone with *TET2* and *DNMT3A* that gave rise to a subclone with *TP53* point mutation at residue 242 that lost the second allele by chromosomal deletion during the course of the disease.

**i)**

| Gene | Protein change | Variant key | MDS VAF | MDS corrected VAF | AML VAF | AML corrected VAF |
|---|---|---|---|---|---|---|
| TP53 | p.C242Y | 17_7577556_C_T | 0.45 | 0.45 | 0.53 | 0.53 |
| DNMT3A | p.N489fs*162 | 2_25468898_TC_T | 0.44 | 0.44 | 0.47 | 0.47 |
| TET2 | p.L952fs*50 | 4_106157950_GCTCTAAGGTGGCATCT_G | 0.42 | 0.42 | 0.42 | 0.42 |

**ii)**

| MDS NGS profile | MDS Cytogenetics | AML NGS profile | AML Cytogenetics |
|---|---|---|---|
| del3p,plus4p,del5q,plus6 | 44,XY,−3,?del(4)(q?),−5,−6,−6,add(10)(q22),−14,+3mar[10] | del3p,plus4p,del5q,plus6,del17p | missing |

**iii) MDS sample**

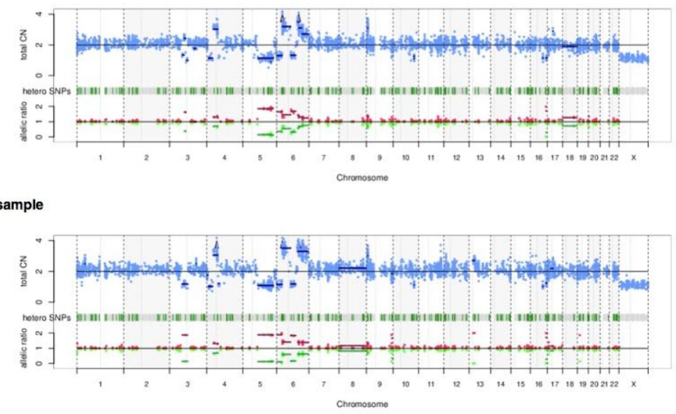


**iv)**

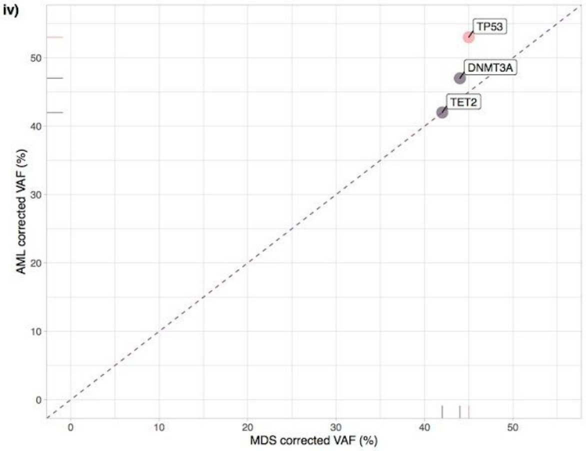


**v)**

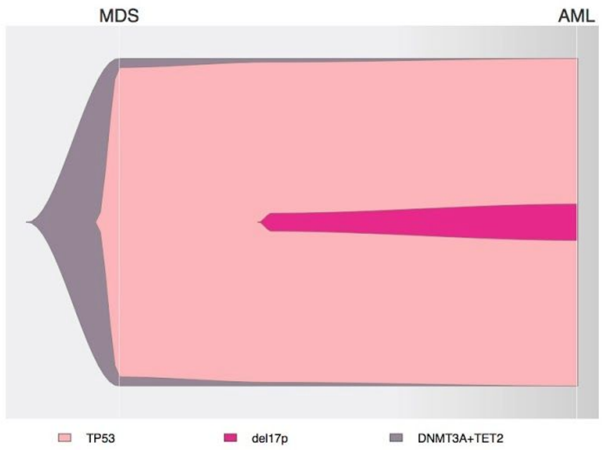

**k. Patient 129047**

The patient had three clonal mutations at MDS and AML (two frameshift mutations in *TET2* and one splice site mutation in *RUNX1)*, along with one *MPL* mutation with a higher VAF at AML than MDS, one *SH2B3* frameshift deletion observed at AML only, one *TP53* mutation observed at AML only at VAF 15%, and *KRAS* G12A hotspot mutation observed at AML only at VAF 4%. The karyotype was normal at both MDS and AML. The clonal dynamics from MDS to AML, with two possibilities represented in v), involves mono-allelic *TP53* with hotspot mutation R273P together with other high-risk genes such as *RUNX1* and *KRAS*.

i)

| Gene | Protein change | Variant key | MDS VAF | MDS corrected VAF | AML VAF | AML corrected VAF |
|---|---|---|---|---|---|---|
| MPL | p.Y591D | 1_43818306_T_G | 0.36 | 0.36 | 0.48 | 0.48 |
| SH2B3 | p.R147fs*50 | 12_111856386_TC_T | 0.00 | 0.00 | 0.93 | 0.46 |
| KRAS | p.G12A | 12_25398284_C_G | 0.00 | 0.00 | 0.04 | 0.04 |
| TP53 | p.R273P | 17_7577120_C_G | 0.00 | 0.00 | 0.15 | 0.15 |
| RUNX1 | p.? | 21_36231876_C_T | 0.47 | 0.47 | 0.53 | 0.53 |
| TET2 | p.C332fs*8 | 4_106156091_T_TA | 0.50 | 0.50 | 0.52 | 0.52 |
| TET2 | p.R1440fs*38 | 4_106193849_G_GA | 0.47 | 0.47 | 0.48 | 0.48 |

ii)

| MDS NGS profile | MDS Cytogenetics | AML NGS profile | AML Cytogenetics |
|---|---|---|---|
| NK | missing | NK | missing |

iii) **MDS sample**

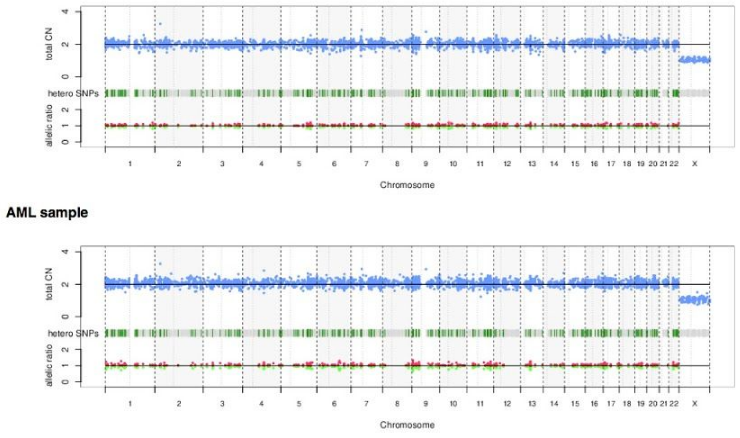


iv)

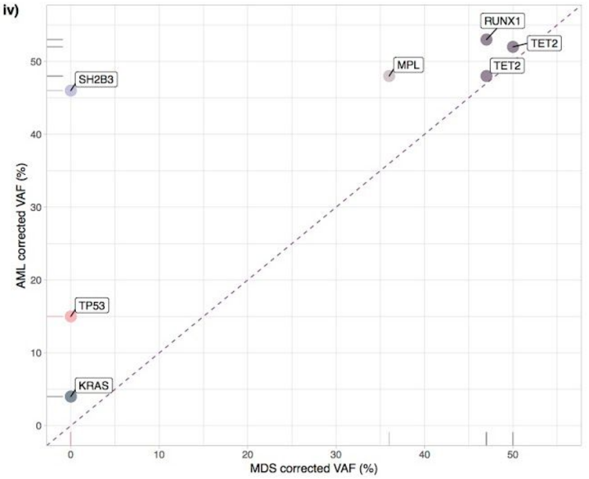


v)

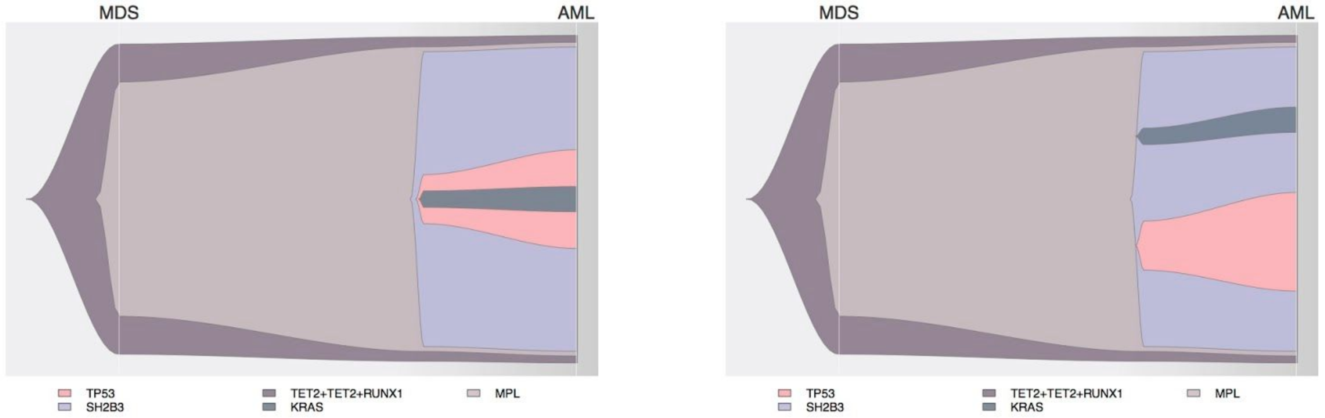

**I. Patient 130333**

The patients had numerous mutations at MDS and AML: clonal mutations in *TET2* and *SRSF2*, one *CBL* mutation with higher VAF in AML than MDS, one *TP53* mutation with slightly higher VAF in AML than MDS, and mutations in *ASXL1* and *KRAS* observed at AML only. Copy-neutral loss of heterozygosity (cnloh) of 11q was observed at MDS and AML (iii), and *CBL* VAF was corrected accordingly (i and iv). No other chromosomal aberration was observed. The clonal dynamics from MDS to AML, with many possibilities represented in v), involves mono-allelic *TP53* with point mutation C272F together with other high-risk genes such as *CBL* and *KRAS*.

i)

| Gene | Protein change | Variant key | MDS VAF | MDS corrected VAF | AML VAF | AML corrected VAF |
|---|---|---|---|---|---|---|
| CBL | p.K389E | 11_119148945_A_G | 0.22 | 0.11 | 0.49 | 0.25 |
| KRAS | p.A59E | 12_25380282_G_T | 0.00 | 0.00 | 0.09 | 0.09 |
| SRSF2 | p.P95H | 17_74732959_G_T | 0.44 | 0.44 | 0.46 | 0.46 |
| TP53 | p.C242F | 17_7577556_C_A | 0.07 | 0.07 | 0.13 | 0.13 |
| ASXL1 | p.R1415* | 20_31024758_C_T | 0.00 | 0.00 | 0.09 | 0.09 |
| TET2 | p.H1077fs*5 | 4_106158328_CA_C | 0.49 | 0.49 | 0.44 | 0.44 |
| TET2 | p.P1747fs*16 | 4_106196905_TC_T | 0.42 | 0.42 | 0.47 | 0.47 |

ii)

| MDS NGS profile | MDS Cytogenetics | AML NGS profile | AML Cytogenetics |
|---|---|---|---|
| cnloh11q | missing | cnloh11q | missing |

iii) **MDS sample**

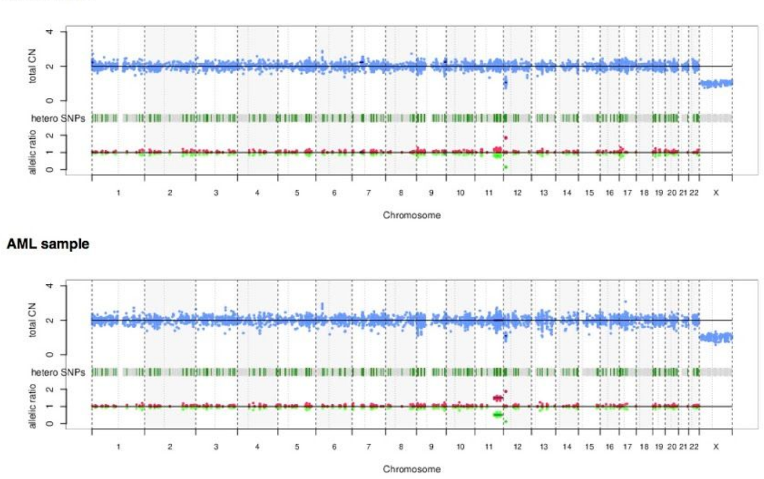


iv)

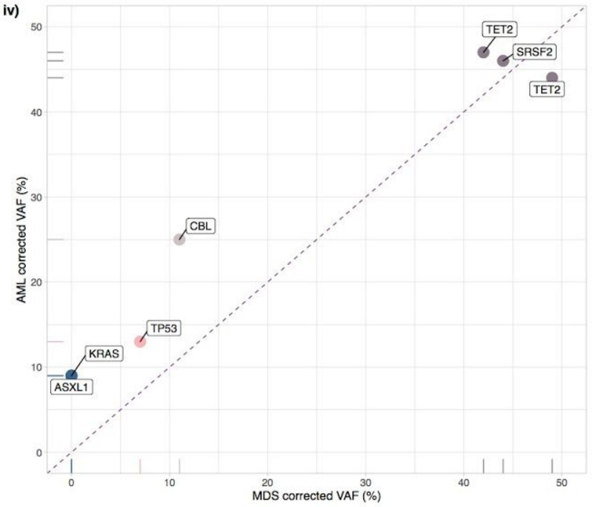

v)

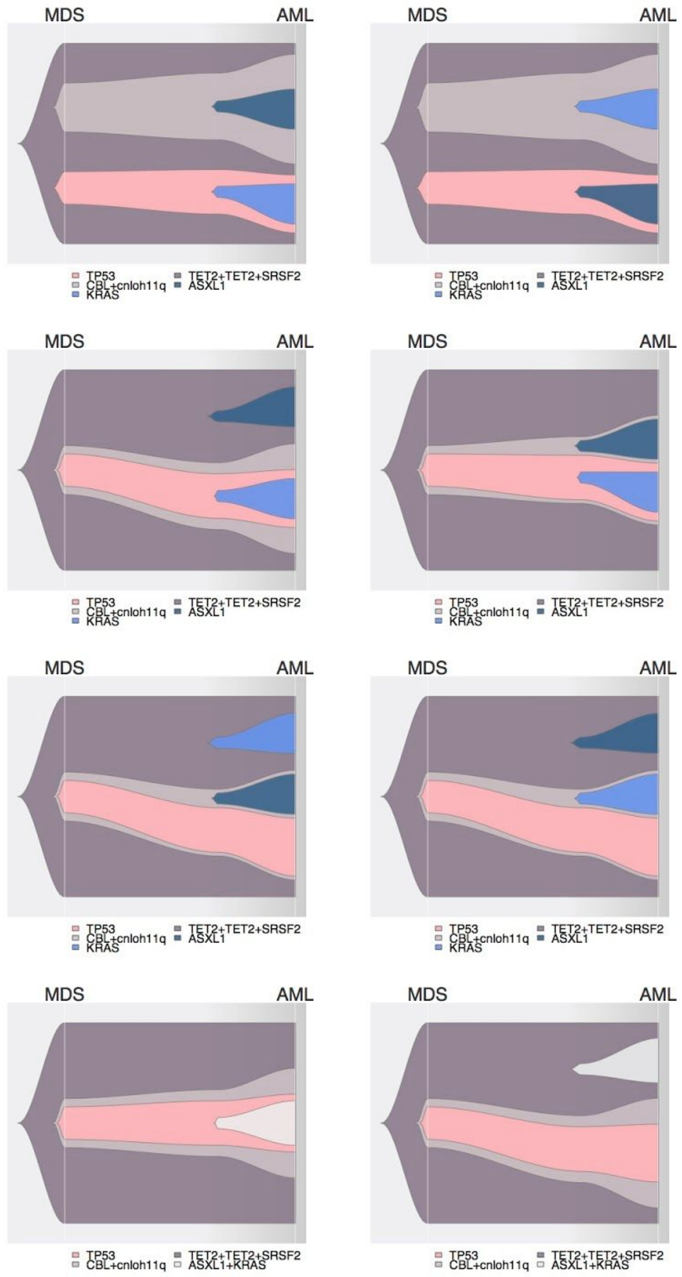
MDS
AML
TP53
CBL+cnloh11q
KRAS
TET2+TET2+SRSF2
ASXL1
MDS
AML
TP53
CBL+cnloh11q
KRAS
TET2+TET2+SRSF2
ASXL1
MDS
AML
TP53
CBL+cnloh11q
KRAS
TET2+TET2+SRSF2
ASXL1
MDS
AML
TP53
CBL+cnloh11q
KRAS
TET2+TET2+SRSF2
ASXL1
MDS
AML
TP53
CBL+cnloh11q
KRAS
TET2+TET2+SRSF2
ASXL1
MDS
AML
TP53
CBL+cnloh11q
KRAS
TET2+TET2+SRSF2
ASXL1
MDS
AML
TP53
CBL+cnloh11q
KRAS
TET2+TET2+SRSF2
ASXL1
MDS
AML
TP53
CBL+cnloh11q
KRAS
TET2+TET2+SRSF2
ASXL1
MDS
AML
TP53
CBL+cnloh11q
KRAS
TET2+TET2+SRSF2
ASXL1
MDS
AML
TP53
CBL+cnloh11q
KRAS
TET2+TET2+SRSF2
ASXL1
MDS
AML
TP53
CBL+cnloh11q
TET2+TET2+SRSF2
ASXL1+KRAS
MDS
AML
TP53
CBL+cnloh11q
TET2+TET2+SRSF2
ASXL1+KRAS
MDS
AML
TP53
CBL+cnloh11q
TET2+TET2+SRSF2
ASXL1+KRAS
MDS
AML
TP53
CBL+cnloh11q
TET2+TET2+SRSF2
ASXL1+KRAS

### Supplementary Figure 18: Representation of *TP53* subgroups and states in the validation cohort

**a.** Number of patients with 1, 2 or 3 mutations in *TP53* or with wild-type *TP53* but a chromosomal aberration at the *TP53* locus in the validation cohort. Colors represent the status of chromosome 17 at the *TP53* locus, to include copy-neutral loss of heterozygosity (cnloh), deletion (del) or no detected aberration (normal). **b.** Frequency of *TP53* subgroups within *TP53*-mutated patients in the validation cohort. The mono-allelic state of single gene mutation (1mut) represents 19% of *TP53* mutated patients, while the multi-hit state encompassing patients with multiple mutations (>1mut), mutation(s) and deletion (mut+del) or mutation(s) and cnloh (mut+cnloh) represent 81% of *TP53*-mutated patients. **c.** Density estimation of variant allele frequency (VAF) of *TP53* mutations across *TP53* subgroups (1mut, >1mut, mut+del, mut+cnloh from top to bottom).

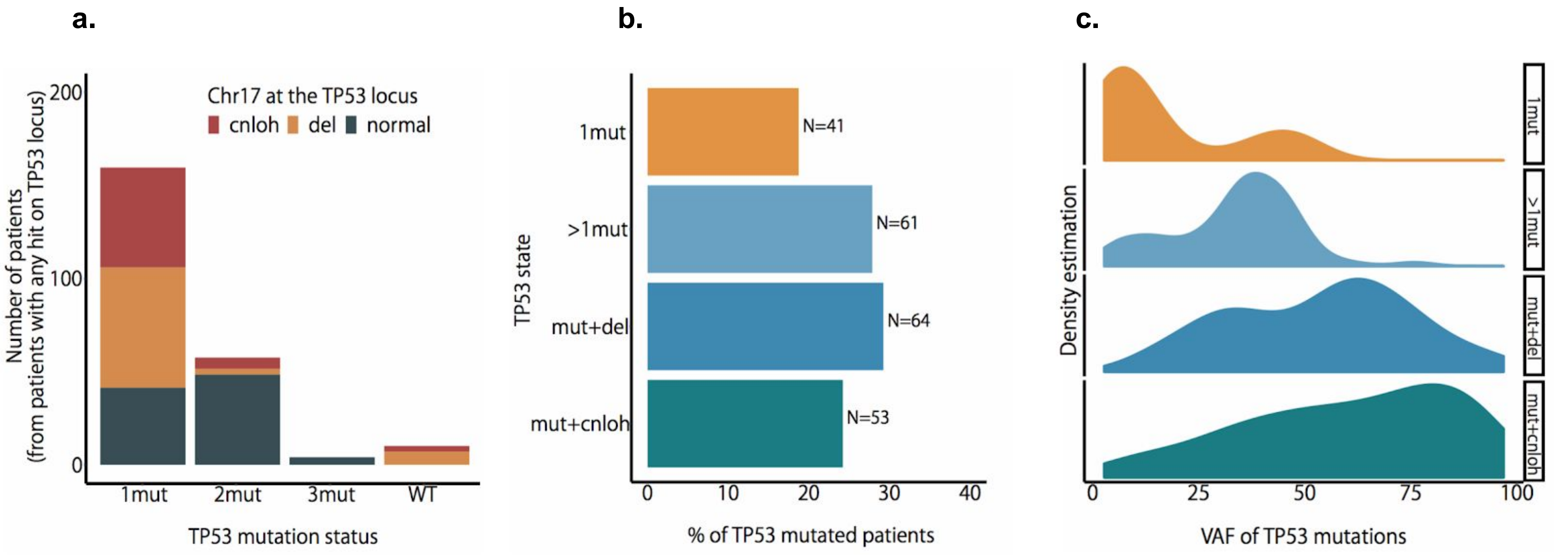

#### Supplementary Figure 19: Implications of *TP53* state to genome stability in the validation cohort

**a.** Number of unique chromosomes other than 17 affected by a chromosomal aberration per *TP53* subgroup in the validation cohort. Dots represent the median across patients, and lines extend from 25% to 75% quantiles. ****p<0.0001, Wilcoxon rank-sum test, compared to the 1mut group. **b.** Distribution of the number of chromosomal aberrations on other chromosomes than 17 per patient across *TP53* subgroups and types of aberrations (cnloh, gain, deletion). ****p<0.0001, Wilcoxon rank-sum test, each compared to the same aberration within the 1mut group. **c.** Heatmap of chromosomal aberrations per *TP53* subgroup. Aberrations include deletion (del), gain (amp), and copy-neutral loss of heterozygosity or equivalently uniparental disomy (upd).

**a.**

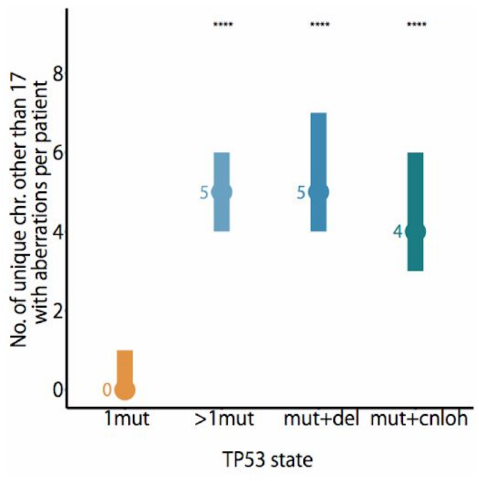


**b.**

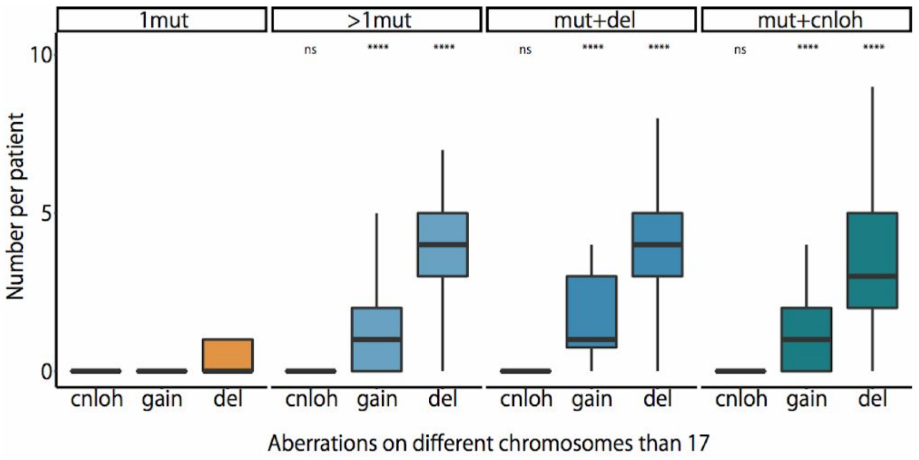

**c.**

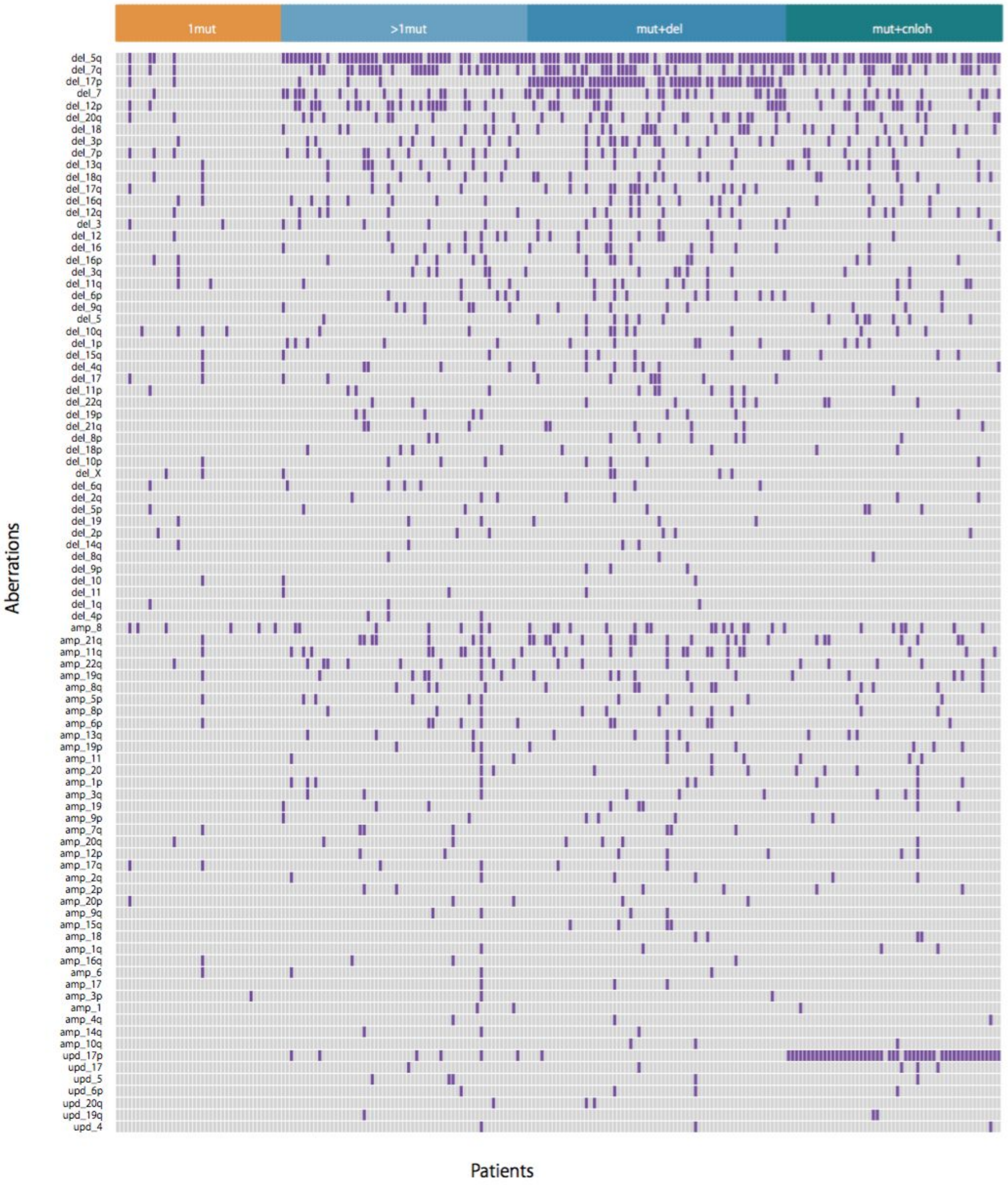
1mut
>1mut
mut+del
mut+cnloh
Aberrations
Patients
del_5q
del_7q
del_17p
del_7
del_12p
del_20q
del_18
del_3p
del_7p
del_13q
del_18q
del_17q
del_16q
del_12q
del_3
del_12
del_16
del_16p
del_3q
del_11q
del_6p
del_9q
del_5
del_10q
del_1p
del_15q
del_4q
del_17
del_11p
del_22q
del_19p
del_21q
del_8p
del_18p
del_10p
del_X
del_6q
del_2q
del_5p
del_19
del_2p
del_14q
del_8q
del_9p
del_10
del_11
del_1q
del_4p
amp_8
amp_21q
amp_11q
amp_22q
amp_19q
amp_8q
amp_5p
amp_8p
amp_6p
amp_13q
amp_19p
amp_11
amp_20
amp_1p
amp_3q
amp_19
amp_9p
amp_7q
amp_20q
amp_12p
amp_17q
amp_2q
amp_2p
amp_20p
amp_9q
amp_15q
amp_18
amp_1q
amp_16q
amp_6
amp_17
amp_3p
amp_1
amp_4q
amp_14q
amp_10q
upd_17p
upd_17
upd_5
upd_6p
upd_20q
upd_19q
upd_4

#### Supplementary Figure 20: Clinical correlates of *TP53* state in the validation cohort

**a-c.** Violin plots indicative of the levels of cytopenias per *TP53* state of a single gene mutation (1mut) or multiple hits (multi), respectively hemoglobin in panel a., platelets in panel b. and absolute neutrophil count (ANC) in panel c. in the validation cohort. Black dots represent the median across patients and black lines extend from 25% to 75% quantiles. **p<0.01 Wilcoxon rank-sum test. **d.** Percentage of bone marrow blasts per *TP53* state of a single gene mutation (1mut) or multiple hits (multi). **e.** Kaplan-Meier probability estimates of overall survival per *TP53* state of wild-type *TP53* (WT), mono-allelic *TP53* per single gene mutation (1mut) and multiple *TP53* hits (multi) in the validation cohort. **f.** Kaplan-Meier probability estimates of overall survival per *TP53* state within the MDS-SLD WHO subtype. **g.** Kaplan-Meier probability estimates of overall survival per *TP53* state within the MDS-EB1 and MDS-EB2 WHO subtypes merged together as MDS-EB1/2.

**a.** **b.** **c.** **d.**

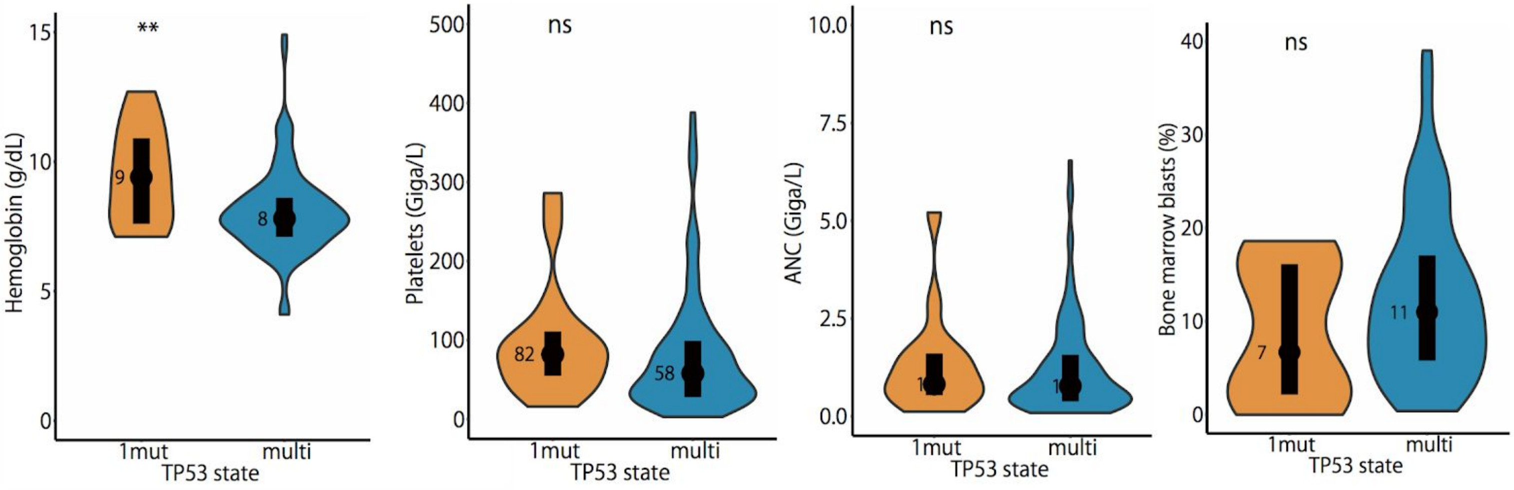

e.

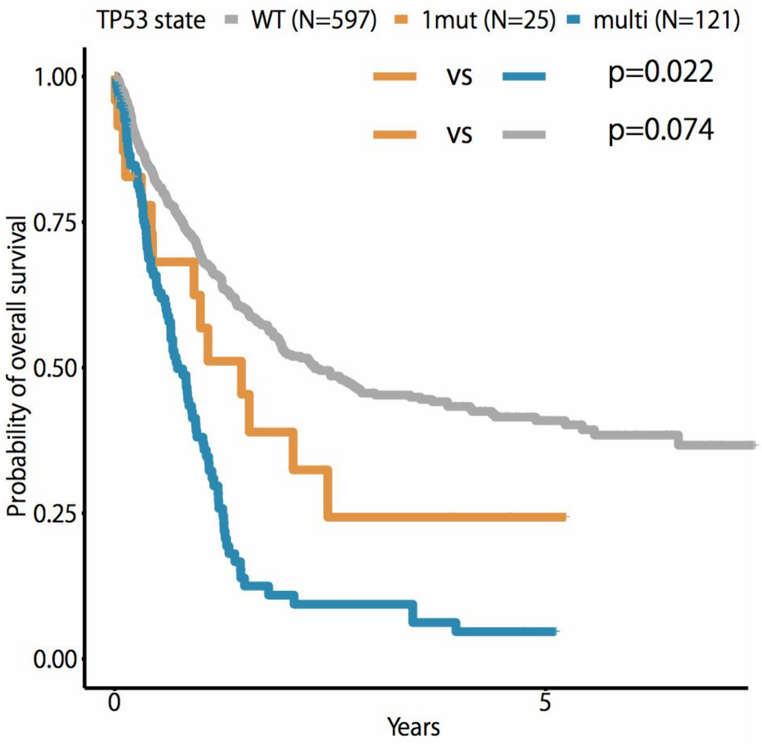


f.

MDS-SLD

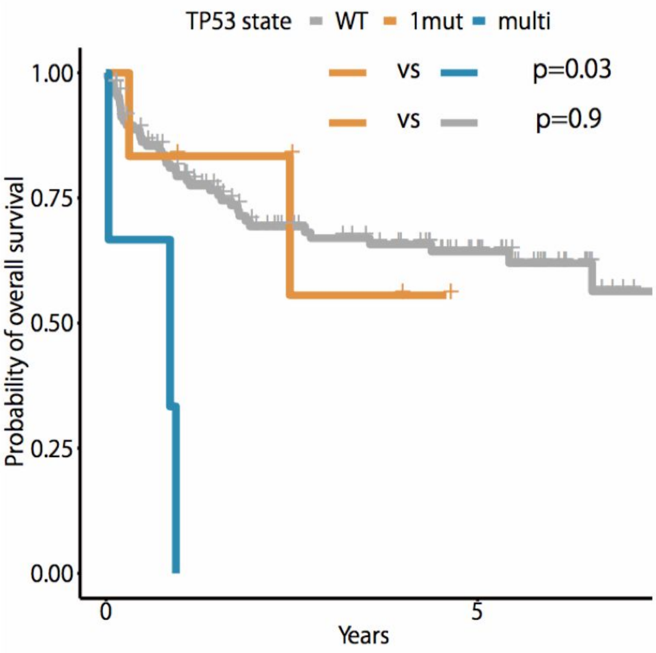


g.

MDS-EB1/EB2

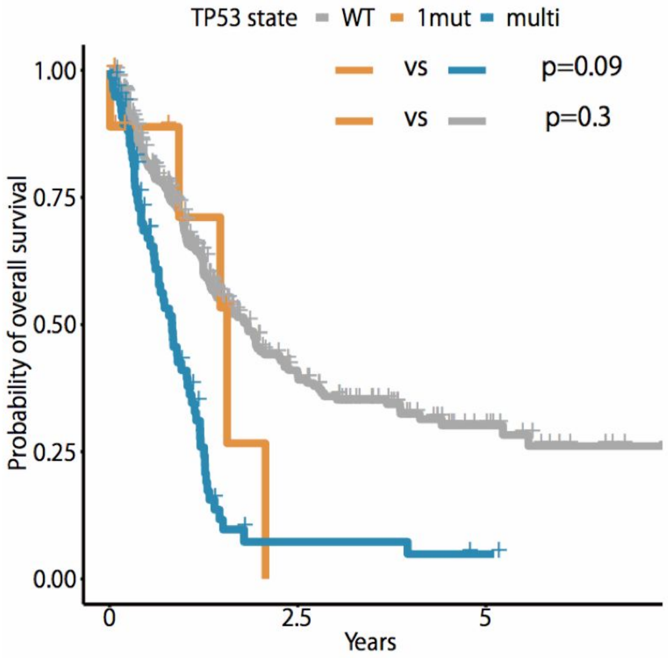
